# Bounded ecological memory in coral reefs: divergent demographic pathways under recurrent heatwaves and collapse under extreme thermal stress

**DOI:** 10.64898/2026.08.21.746008

**Authors:** Tooba Varasteh, Andy P. Curran, Olivia Mathews, Shanna L. Davidson, George Warfel, Fiona Warsfold, Kayla Shen, Vadim Backman, Luisa A. Marcelino

## Abstract

Anthropogenic warming exposes coral reefs to recurrent marine heatwaves (MHWs), yet responses to comparable thermal stress remain highly variable. A central challenge is determining whether reduced bleaching reflects acclimatization-like persistence of existing colonies or mortality-driven filtering that produces demographically depleted assemblages. Using colony-level surveys across the Florida Reef Tract (2014–2023), we show bleaching sensitivity declined from 2014 through 2022, consistent with ecological memory. A spatial matched-event framework integrating bleaching severity and adult colony density resolved these reduced-bleaching outcomes into two distinct pathways: demographic persistence (stable/increasing density) and mortality-filtered tolerance (declining density). Persistence pathways were maintained under predictable exposure regimes characterized by lower interannual thermal variability. This persistence was driven by weedy life-history taxa (e.g., *Porites spp.*) replacing historical framework builders, indicating stability reflects ecological reassembly rather than uniform increases in thermal tolerance. Under the unprecedented extremes of the 2023 MHW, this acclimatization-like buffering broke down, causing widespread sensitization and collapse of previously persistent networks. These results support a bounded ecological memory framework; prior exposure reduces bleaching sensitivity under predictable thermal regimes but fails under extreme heat stress. Consequently, modern refugia are best defined by adult standing stock retention representing systems undergoing selective ecological reassembly rather than full functional recovery.

## INTRODUCTION

Anthropogenic ocean warming is increasing the frequency, duration, and intensity of marine heatwaves (MHWs), exposing coral reef communities to recurrent thermal stress before populations can fully recover [1–4]. These events can trigger coral bleaching, disrupting the coral-algal symbiosis that underpins growth, survival, and reef ecosystem function [5–7]. Yet similar levels of thermal stress can produce markedly different bleaching and demographic outcomes across space and time [4, 8, 9]. Consequently, predicting reef responses to recurrent heat stress remains a major challenge.

A growing body of work suggests that these divergent outcomes reflect ecological memory, whereby prior exposure influences the response to subsequent disturbances [8, 10, 11]. This history dependence spans colony-level environmental memory [10, 11] and emergent, ecosystem-level ecological memory [4, 8]. At the ecosystem scale, repeated heat stress can modify bleaching sensitivity and generate responses not predictable from single events alone [4, 8]. In contrast, colony-level memory is mechanistically diverse, spanning host physiology, symbiosis, microbiomes, and epigenetic regulation [10, 11].

History dependence is not uniformly beneficial. Prior exposure may reduce bleaching sensitivity under some conditions, but exacerbate responses when recovery is incomplete [9, 11–13]. Increasing evidence suggests these effects are bounded, such that buffering during moderate events may weaken or reverse under more severe heatwaves [4, 14]. Together, these findings imply that carryover effects operate within physiological and ecological limits, especially as heat stress intensifies and recovery intervals shorten [4, 9, 14].

A central unresolved question is whether reduced bleaching following repeated heat exposure reflects true persistence of coral populations or arises through selective mortality that reshapes community composition. Similar bleaching outcomes can emerge through fundamentally different processes: acclimatization-like buffering that preserves established colonies, or mortality-filtered tolerance in which susceptible individuals are lost, leaving a more tolerant but demographically depleted assemblage [12, 13, 15, 16]. Distinguishing these pathways is essential to forecasting reef futures: demographic persistence preserves reproductive capacity and potential for functional continuity, whereas mortality-driven filtering can stabilize a degraded assemblage while eroding reef-building capacity and long-term recovery potential [9, 17–19].

Here, we address this challenge by integrating bleaching responses with demographic change using a matched-event framework that classifies reef trajectories based on joint changes in bleaching severity and adult colony density. Focusing on adult standing stock allows us to distinguish among sensitization, demographic persistence, and mortality-filtered tolerance, and to test whether reduced bleaching reflects persistence or demographic filtering. We further ask whether ecological memory depends not only on cumulative heat exposure but also on the temporal structure of repeated disturbances, evaluating whether variability among successive marine heatwaves influences the persistence of acclimatization-like responses. This approach builds on evidence that bleaching outcomes depend on multiple aspects of thermal exposure, including variability and recovery dynamics, rather than single threshold metrics alone [20–23].

The Florida Reef Tract (FRT) provides an ideal system to test these ideas because it has experienced multiple marine heatwaves spanning a wide range of intensities [18, 19], while maintaining extensive long-term colony-level bleaching and demographic monitoring (Disturbance Response Monitoring-Florida Reef Resilience program, DRM-FRRP; [24]) The combination of repeated disturbances, heterogeneous environmental conditions, and large-scale observational data enables evaluation of bleaching sensitivity and demographic change across space and time.

Here, we use the extreme 2023 marine heatwave as a natural stress test of ecological memory. Specifically, we ask whether: (1) prior heat exposure modifies bleaching sensitivity, consistent with ecosystem-scale ecological memory [4, 8]; (2) reduced bleaching reflects demographic persistence or mortality filtering, representing acclimatization-like retention of established colonies versus selective removal of heat-susceptible individuals; (3) how these responses depend on the temporal structure of heat exposure, particularly whether repeated heat stress reinforces persistence or instead leads to breakdown under extreme conditions; and (4) persistent refugia reflect broad community uplift or selective gains among a subset of taxa.

## METHODS

### DRM-FRRP coral bleaching dataset and marine heatwaves selection

We analyzed coral bleaching and demographic data from Disturbance Response Monitoring/Florida Reef Resilience Program (DRM-FRRP), across the 251 km² Florida Reef Tract (FRT) [24], between 2005 and 2023. Surveys followed a randomized design that sampled two 10 × 1 m belt transects per site across habitats and subregions without tracking fixed colonies or permanent sites through time. Colony condition was recorded for colonies ≥4 cm including bleaching and mortality states. We focused on five focal marine heatwaves (2014, 2015, 2020, 2022, 2023) selected based on heat-stress intensity (≥ 30% of sites exposed to DHW ≥ 4.0) and sampling coverage (≥150 sites; Table S1; Supplementary SM1-2).

### Bleaching severity index and Bleaching severity

Following Swain et al. (2016) we converted categorical colony condition observations to quantitative tissue-affected ranges and assigned midpoint values [25]. For each species *j* at site *k*, mean bleaching severity (*B_jk_*) was weighted by the proportion of colonies in each category. Site-level assemblage severity (*B_site_*) was computed as the unweighted mean of *B_jk_* to avoid dominance effects across the FRT’s spatial gradients in species composition. Severe bleaching was defined as *B_site_* ≥ 0.30 [8] (Supplementary SM3).

### Heat Stress exposure (NOAA CRW-DHW) and spatial alignment

Thermal stress exposure was quantified using NOAA Coral Reef Watch (CRW) Degree Heating Weeks (DHW, expressed as °C-weeks) at 5 km (0.05°) resolution (v3.1) [26]. DRM-FRRP survey sites were mapped to the nearest 5 km grid cell and DHW_site_ was defined as the maximum DHW within a 14-day window preceding the survey date. Because surveys span August–November, differences in survey timing produce multiple DHW_site_ values within a single grid-year. Thus, we utilized two-step averaging, where site-level metrics were averaged to compute grid-level DHW_grid_ and *B_grid_* (Supplementary SM4–SM5). Differences in non-normally distributed heat stress among years were evaluated using Kruskal-Wallis (for independent, full-tract comparisons) and Friedman (for paired, repeatedly surveyed grids) tests, followed by Bonferroni-adjusted Wilcoxon tests.

### Severe-bleaching reaction norms

To quantify shifts in thermal sensitivity across successive MHWs, year-specific bleaching reaction norms were modeled as a function of DHW using a binomial Generalized Linear Model (GLM) with a logit link function [8], where bleaching severity was binary-coded (0 = non-severe, *B_site_* <0.3; 1 = severe, *B_site_* ≥0.3]).

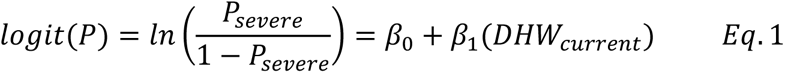

where *P_severe_* is the probability of severe bleaching, *β_0_* is the baseline bleaching probability intercept, *β_1_* captures thermal sensitivity (i.e., shifts in the slope of the heat stress–bleaching relationship), and DHW_current_ represents the DHW of a surveyed site in the year under examination. Models were fit across all FRT sites in each focal year. Sensitivity experiments validated these trends across (i) repeatedly surveyed grids to reduce sensitivity to spatial heterogeneity and year-to-year shifts in sampled sites, and (ii) grid-level aggregates, to evaluate spatial aggregation. Species-specific reaction norms were fit analogously for eleven coral species with sufficient sample sizes across the FRT. Statistical significance of the relationship between heat stress and bleaching was assessed using Wald z-tests on the *β_1_* coefficient for each year-specific model (Supplementary SM6).

### Ecological Memory interaction models

To test whether prior exposure modifies bleaching sensitivity, we applied Hughes et al. 2021 [4] ecological memory framework that combines “Year Alone” model (Eq. 1) with an “Interaction” between current-year and baseline (2014) MHW, for each pairwise window (2014–2015, 2014-2020, 2014–2022, and 2014–2023):

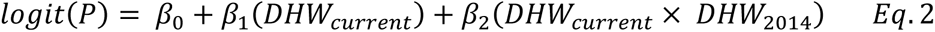

Where β_1_ represents the main effect of heat stress during the current anomaly, and β_2_ represents the effect of baseline exposure on current bleaching sensitivity (ecological memory; negative β_2_). Following Hughes et al. (2021) [4], the DHW_2014_ main effect was omitted to constrain intercepts, ensuring *β*_2_ tests whether prior exposure modifies bleaching sensitivity (the slope) rather than shifting baseline bleaching probability. Model support was evaluated using ΔAIC and McFadden’s pseudo-R² and the significance of the ecological memory effect (β_2_) was assessed using Wald z-tests, with 95% confidence intervals (Supplementary SM7).

### Matched-Event trajectory analysis

We isolated demographic patterns by conducting pairwise matched-event analyses, comparing repeatedly surveyed grid cells in a given subsequent year directly to the 2014 baseline. For each pairwise window, we computed grid-level change in bleaching severity (ΔB_grid_) and adult colony densit (CD_adult,grid_) for a pooled 13-species assemblage:

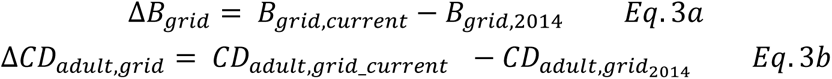

(Supplementary SM8-SM9). To ensure thermal comparability, grid-level DHW_current_ was evaluated relative to DHW_2014_ using a regression envelope (±1.5 SD of residuals) to flag and exclude thermally atypical trajectories. Retained grids were classified into three non-mutually exclusive processes: (1) sensitization (H2), displaying increased bleaching severity (ΔB_grid_ > 0), (2) persistence (H3), showing reduced bleaching (ΔB_grid_ ≤ 0) accompanied by stable or increasing adult density (ΔCD_grid,adult_ ≥ 0), consistent with physiological acclimatization and (3) mortality-filtered tolerance (H4), showing reduced bleaching (ΔB_grid_ ≤ 0) accompanied by declining adult density (ΔCD_grid,adult_ < 0), consistent with selective mortality of heat-sensitive colonies.

### Demographic modeling and robustness checks

Adult colony density served as a proxy for persisting standing stock, because juvenile recruitment can increase even while adult populations decline, potentially masking demographic loss. Adult status was assigned using species-specific size thresholds reflecting life-history growth strategies [17, 27]: (e.g., W5/S10 (weedy adults ≥5 cm; stress-tolerant/generalist adults ≥10 cm diameter). Adult density was calculated for a pooled assemblage of 13 abundant and ecologically important species, using true zeros for surveyed sites lacking adult colonies before aggregating to grid-year means (Supplementary SM9). To test whether density trajectories differed significantly between H3 and H4 grids, we fitted negative binomial generalized linear models to site-year adult colony counts:

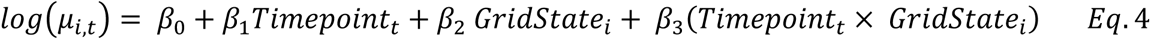

where µ_i,t_ is the expected adult colony density at site *i* and timepoint *t*; β_0_ represents baseline density, β_1_ captures temporal change, β_2_ represents baseline differences between H3 and H4 grids, and β_3_ quantifies divergence in demographic trajectories between persistence and mortality-filtered networks (Wald z-tests, 95% CI). GridState was assigned at the grid level and inherited by site-year observations. Robustness was evaluated across adult-size thresholds and juvenile-density models (SM9). Parameter estimates are reported with 95% Wald confidence intervals.

### Stable and Ultimate network definitions and baseline characterization

Stable networks were defined using only pre-2023 trajectory classifications (2014-2015, 2014-2020, 2014-2022), requiring consistent assignment to H3 or H4 across at least two windows without conflicting classifications (e.g., H3 grids: ≥2 windows and never H2/H4; Supplementary SM10). “Ultimate” networks were the subset that retained their classification during the 2023 MHW. Baseline demographic and thermal metrics were compared using Mann-Whitney U tests, with Cliff’s δ (95% CI) as an effect-size metric.

### Species decomposition and sensitivity analysis

To identify species driving divergence between H3 and H4 networks, we quantified species-specific changes in adult density (ΔCD_adult,grid_) for 22 taxa, and compared groups using Welch’s t-tests to account for unequal variances. Sensitivity analyses on four focal taxa (*PAST, PPOR, SSID, MCAV*) used independent pre-2023 Stable network classifications (Supplementary SM11).

## RESULTS

### Heat stress and severe bleaching across the five focal marine heatwaves

Coral assemblages across the Florida Reef Tract (FRT; 4,546 sites mapped to 1,712 5 x 5 km grid cells) experienced substantial variation in cumulative heat stress between 2005 and 2023, (Fig. 1; Table S1). Thermal stress was generally low prior to 2014, and severe bleaching (B_site_ ≥ 0.30) was uncommon (<5% of sites in most years; Fig. S1; Table S1). The 2014 marine heatwave (MHW) produced widespread severe bleaching (∼50% of sites; Fig. 2). Subsequent events in 2015, 2020, and 2022 reached moderate to high heat stress intensities, but resulted in substantially lower severe bleaching prevalence (Figs. 1B, 2).

**Figure 1.**
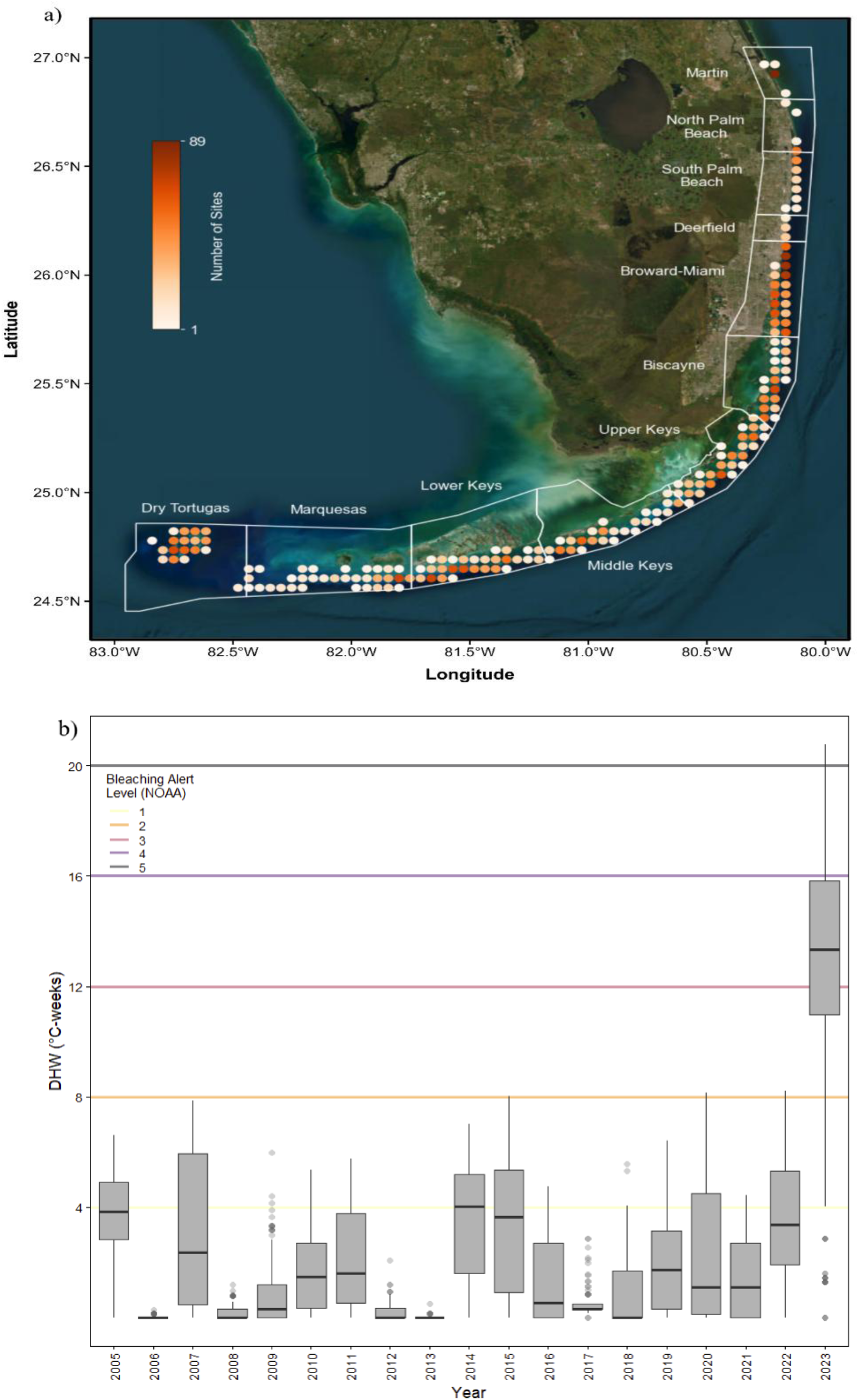
Spatial survey coverage and heat-stress context across the Florida Reef Tract (2005–2023). (a) DRM–FRRP survey effort mapped to NOAA Coral Reef Watch (CRW) 5 x 5 km grid cells, showing site sampling intensity [1 site (white) to 89 sites (dark orange) per grid]. (b) Distribution of CRW Degree Heating Weeks (DHW, °C-weeks) across survey years. DHW represents the maximum 14-day pre-survey exposure. Horizontal lines indicate CRW Bleaching Alert Levels 1 (4 ≤ DHW < 8) through 5 (DHW ≥ 20) (Table S1).

**Figure 2.**
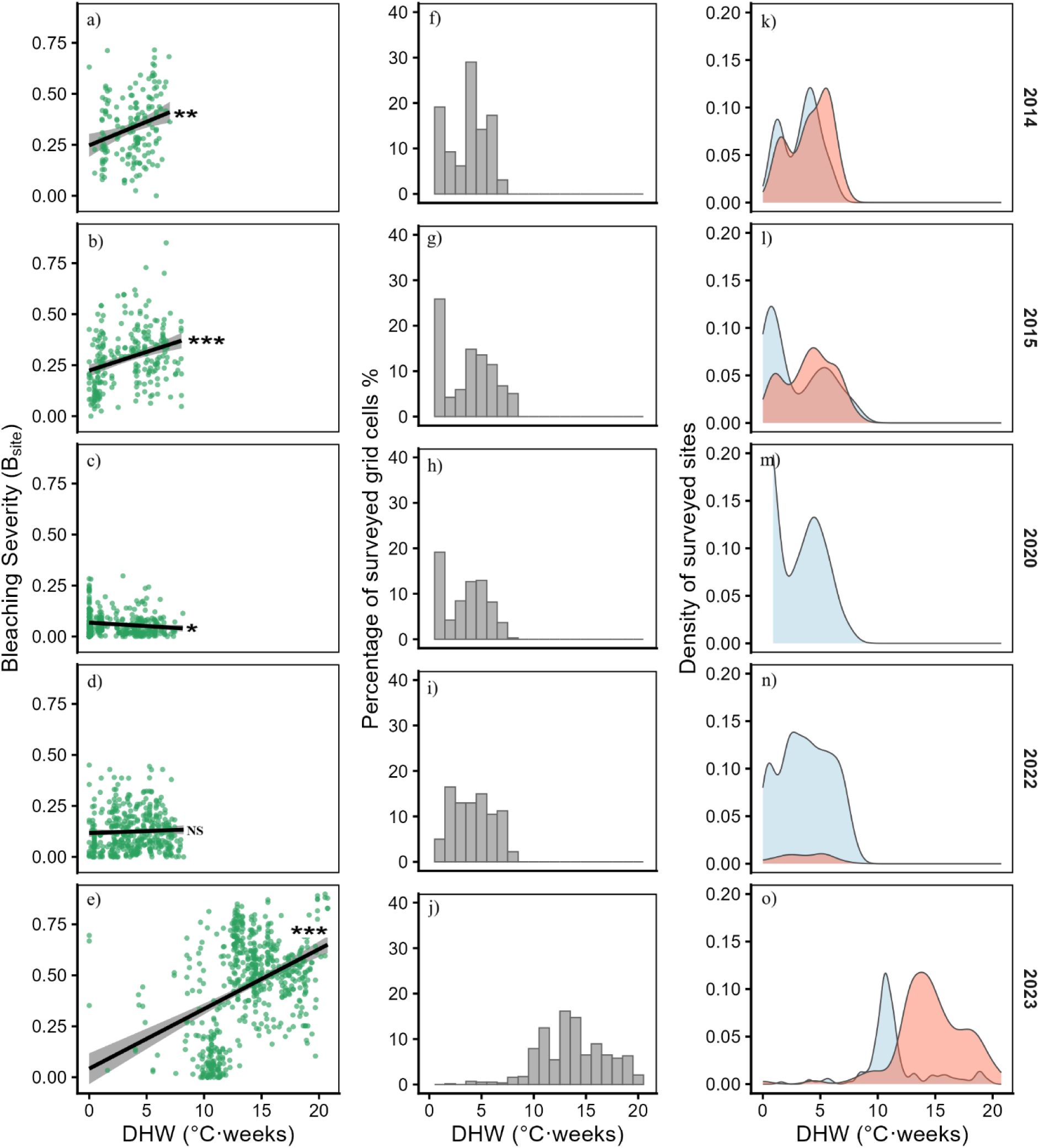
Five focal marine heatwaves (MHWs) show contrasting bleaching responses under measured heat stress. Bleaching severity and heat exposure for 2014, 2015, 2020, 2022, and 2023. (a–e) Site-level bleaching severity (B_site_) versus heat stress (DHW, °C-weeks; NS p> 0.05; * p< 0.05; ** p< 0.01; *** p<0.001). (f–j) Frequency distributions of surveyed grid cells across binned DHW. (k–o) Distribution of severe (orange; B_site_ ≥ 0.30) versus non-severe (blue) site-level bleaching outcomes (Fig. S1; Table S1).

To quantify these shift, we modeled year-specific bleaching reaction norms as the probability of severe bleaching (*P_severe_*) as a function of DHW (Eq. 1; Fig. 3A; Table S2). Reaction norms shifted markedly over time. In 2014, a 70% probability of severe bleaching occurred at 6.5 °C-weeks, whereas in 2015, this threshold shifted to 8.6 °C-weeks. In 2020 and 2022, reaction norms flattened because severe bleaching was rare (Fig. 3A; Table S2). Although the 2020 event was cooler than 2014 (p<0.05; Table S3), 2022 exhibited comparable heat stress yet severe bleaching remained rare, supporting the interpretation of a reduction in bleaching sensitivity consistent with memory effects (Fig. 3B; Tables S2-S3). The 2023 MHW exposed reefs to unprecedented thermal anomalies (8 –20°C-weeks) and resulted in widespread severe bleaching and raising the 70% severe bleaching threshold to ∼12.9 °C-weeks (Figs. 2–3).

**Figure 3.**
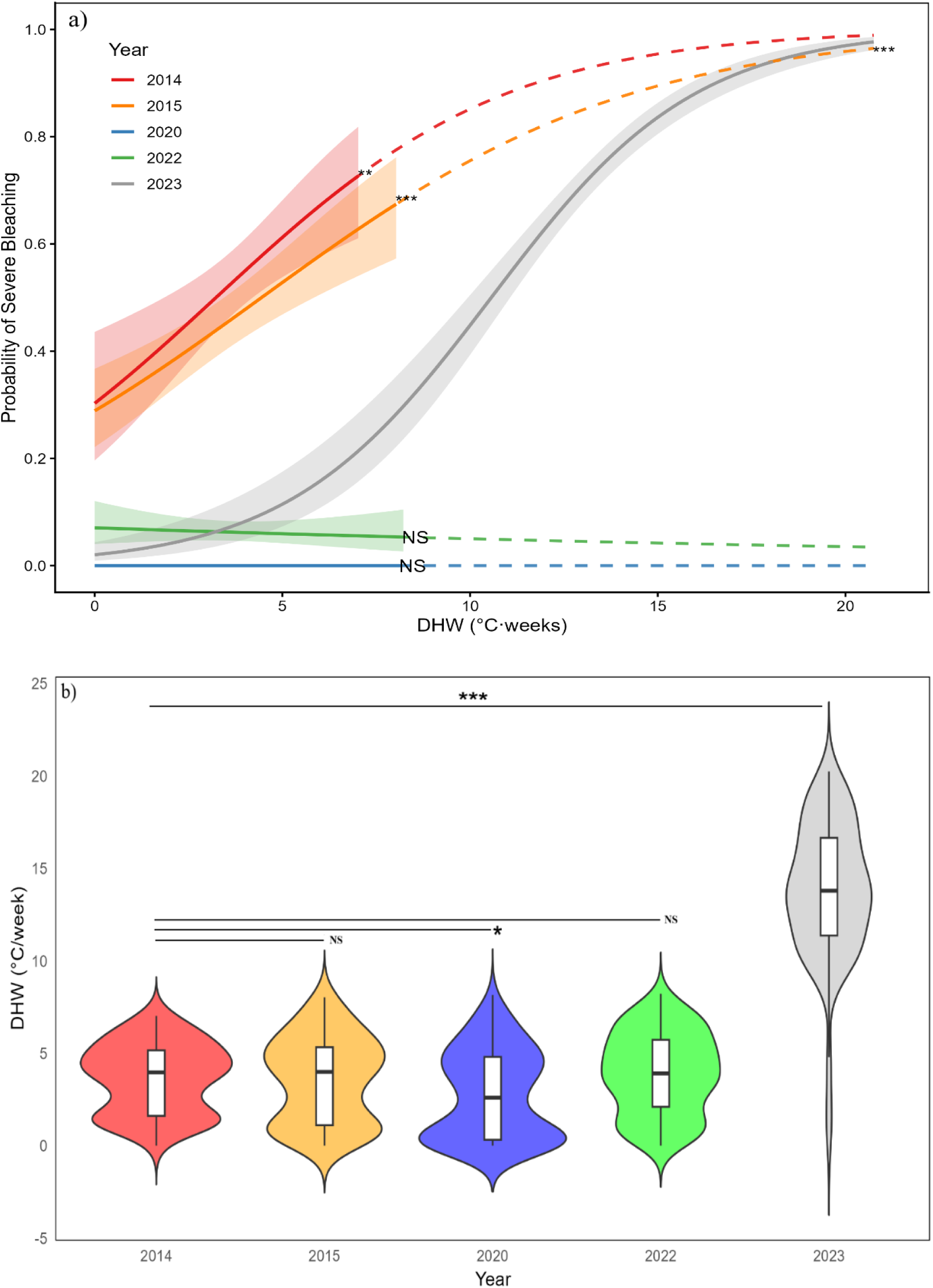
Tract-wide severe-bleaching reaction norms shift across successive MHWs, with 2023 as an extreme stress test. (a) Year-specific binomial GLMs relating the probability of severe bleaching (B_site_ ≥ 0.30) to DHW across all surveyed sites in the Florida Reef Tract across the five focal MHWs. Solid lines with ±1 SD shading denote the observed DHW range; dashed lines represent extrapolated model fits beyond observed values (Eq.1; Wald-z test). (b) Violin plots of the distributions of site-level DHW for each focal year (Kruskal-Wallis, Bonferroni-adjusted Wilcoxon tests; Tables S2-S3).

Sensitivity analysis restricted to repeatedly surveyed grid cells (n= 48 grids; 928 sites) produced simlar reaction norm shifts (Fig. S2), indicating tract-wide patterns were not driven by spatial sampling variation. Grid-level (5 x 5 km) models also reproduced these trends, confirming robustness to spatial aggregation (Fig. S3; Table S2).

Species-level analyses showed similar directional patterns across some of the most abundant coral taxa (*SSID, PAST, SINT, MCAV, AAGA, PPOR, SRAD, OFAV, DSTO, PSTR, CNAT*), with reduced severe bleaching in 2015-2022 and increased bleaching under 2023 conditions observed for most species (Fig. 4; Table S2), althought statistical power was limited for some species due to smaller sample sizes.

**Figure 4.**
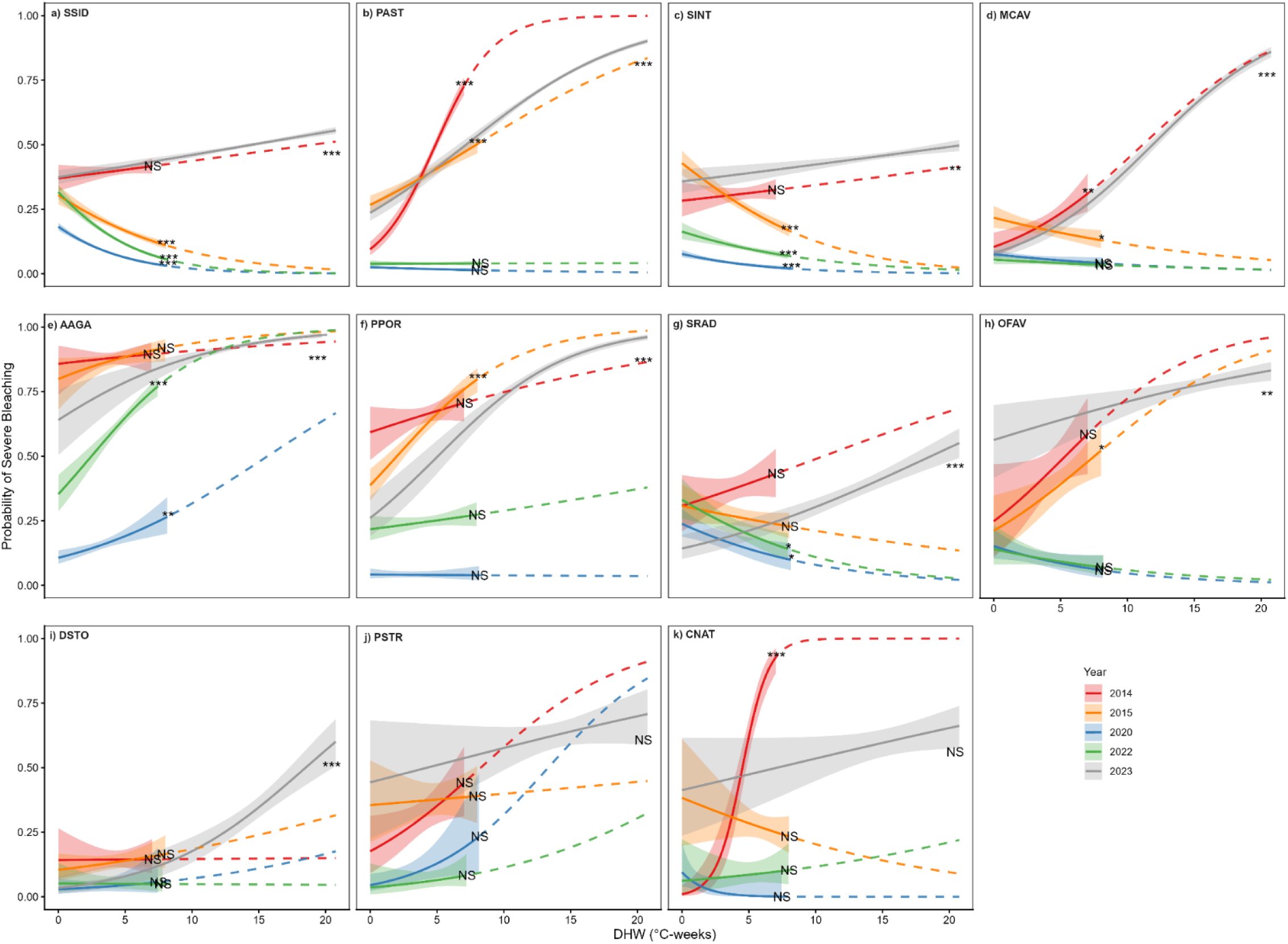
Species-specific severe-bleaching reaction norms across five MHWs for 11 abundant taxa. Year-specific binomial GLMs relating the probability of severe bleaching to DHW for eleven taxa with sufficient colony counts. Solid and dashed lines, and shading as in Fig. 3 (Eq. 1; Wald z-test; Table S2).

### Ecological memory alters bleaching sensitivity to current heat stress

To test whether prior heat exposure affected bleaching sensitivity, we compared models with and without an interaction between current heat stress (DHW_current_) and baseline exposure during the 2014 MHW (DHW_2014_).

During the 2014–2015 back-to-back events, severe bleaching was common (79/180 sites), and inclusion of the interaction term improved model fit (ΔAIC = −3.25; Table 1). The interaction coefficient was significantly negative (β₂ = −0.09, 95% Wald CI [0.32 - 1.34], p < 0.05), indicating a lower probability of severe bleaching at sites with higher DHW_2014_. In 2020, following a period of relatively low thermal stress (2016 through 2019), severe bleaching was absent (0/251 sites), precluding statistical estimation of binomial slopes and interaction effects (Table 1). In 2022, severe bleaching remained low (21/290 sites) and the Interaction model again outperformed the Year Alone model (ΔAIC = −6.66) with a significantly negative interaction term (β₂ = −0.15, 95% Wald CI [0.24 - 1.56], p < 0.01; Table 1).

**Table 1.** Ecological memory interaction models show that baseline exposure reduces bleaching sensitivity and reverses under the 2023 extreme. Severe bleaching probability (B_site_ ≥ 0.30) was modeled via site-level binomial GLMs comparing 2014 to subsequent MHWs in repeated surveyed grids (n = 63, 68, 71, and 74). A “Year-Alone” model (Eq.1, DHW_current_) was compared to an “Interaction model” (Eq., 2, DHW_current_ × DHW_2014_). Metrics include sample size, model fit (McFadden’s pseudo-R², ΔAIC), and coefficient, with 95% Wald CIs. Negative β₂ indicates reduced sensitivity.

| Pairwise Window | Sample size |  | Year Alone Model |  |  | Interaction model |  |  |  |  | model comparison |
| --- | --- | --- | --- | --- | --- | --- | --- | --- | --- | --- | --- |
| | Number Sites current | Number Severe Sites current ($B_{\text{site}} \geq 0.3$) | $DHW_{\text{current}} (\beta_1)$<br>[95% CI] | p-value | Pseudo-$R^2$ | $DHW_{\text{current}} (\beta_1)$<br>[95% CI] | p-value | Pseudo-$R^2$ | Interaction ($\beta_2$) | p-value | $\Delta AIC$ (Alone - Interaction) |
| 2014-2015 | 180 | 79 | 0.328 [0.186, 0.47] | <0.001 | 0.101 | 0.814 [0.324, 1.304] | 0.001 | 0.122 | -0.085 | 0.028 | -3.248 |
| 2014-2020 | 251 | 0 | NA | NA | NA | NA | NA | NA | NA | NA | NA |
| 2014-2022 | 290 | 21 | 0.013 [-0.192, 0.219] | 0.898 | 0.002 | 0.901 [0.239, 1.563] | 0.008 | 0.059 | -0.146 | 0.007 | -6.66 |
| 2014-2023 | 367 | 241 | 0.448 [0.322, 0.573] | <0.001 | 0.239 | -0.108 [-0.277, 0.062] | 0.214 | 0.515 | 0.11 | <0.001 | -128.24 |

In 2023, severe bleaching prevalence was high (241/367 sites), and the Interaction model strongly outperformed the Year Alone model (ΔAIC = −128.24). However, the interaction coefficient reversed sign and became significantly positive (β₂ = +0.11, 95% Wald CI [-0.28 -0.06], p < 0.001, Table 1).

### Spatial heterogeneity in trajectories reveals sensitization, persistence, and mortality-filtered pathways

Using the 5 x 5 km grid framework, we evaluated spatial variation bleaching severity and demographic change across matched-event comparisons relative to 2014. Thermally comparable grids were identified based on the relationship between DHW_2014_ and DHW_current_ (±1.5 SD envelope), with thermally atypical grids excluded from classification (Fig. 5A-C; Table S4).

**Figure 5.**
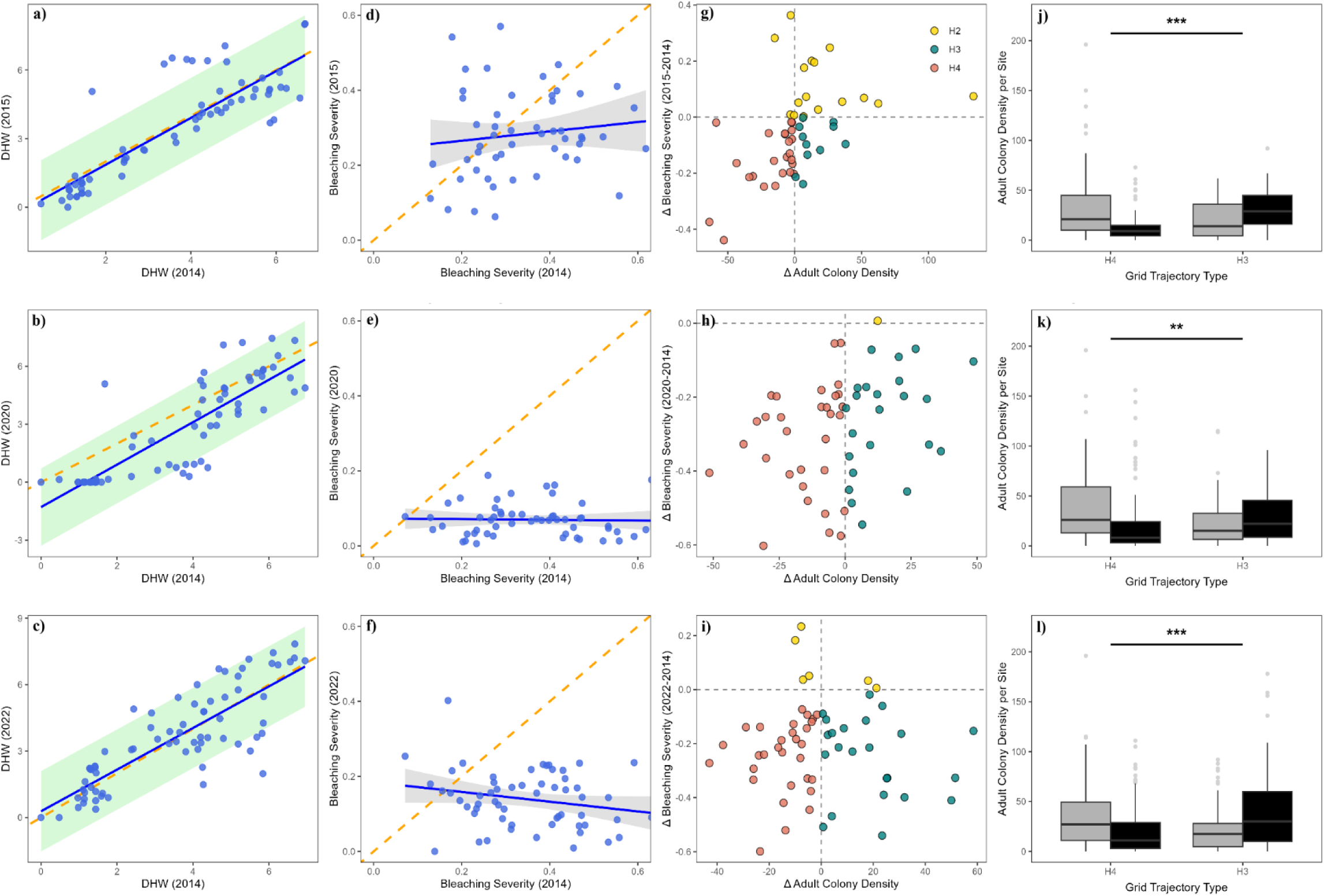
Pre-2023 matched-event framework distinguishes sensitization, persistence, and mortality-filtered tolerance under comparable heat stress. Pairwise classifications relative to 2014 (2015: 314 sites/63 grids; 2020: 406/68; 2022: 450/71). (a-c), DHW comparability; thermally typical grids (within ±1.5 SD envelopes) are retained. (d-f), Grid-level bleaching outcomes (B_grid_). Solid lines denote linear fits (R^2^ = 0.017; 0.001; 0.053, p> 0.05) vs. 1:1 expectations (dashed). (g-i) Trajectory classes (Eq. 3): H2 sensitization (ΔB_grid_ > 0), H3 persistence (ΔB_grid_ ≤ 0 and ΔCD_adult,grid_ ≥ 0), or H4 mortality-filtered (ΔB_grid_ ≤ 0 and ΔCD_adult,grid_ < 0). Adult colony density (CD_adult_) Adult density using W5/S10 thresholds. (j-l) CD_adult,site_ for H3 and H4 grids in 2014 (grey) and current year (black). Trajectory divergence assessed using negative binomial GLMs (Eq.4; ** p<0.01; *** p<0.001; Tables S4-S5).

Across thermally comparable grids, joint changes in bleaching severity (ΔB_grid_, Eq.3a) and adult colony density (ΔCD_adult, grid_, Eq.3b) identified three distinct trajectory classes: sensitization (H2; ΔB_grid_ > 0), persistence (H3; ΔB_grid_ ≤ 0 and ΔCD_adult, grid_ ≥ 0), and mortality-filtered tolerance (H4; ΔB_grid_ ≤ 0 and ΔCD_adult, grid_ < 0) (Fig. 5G-I; Table S4).

Bleaching trajectories varied across pairwise windows (Fig. 5D-F). In 2014–2015, responses were heterogeneous, with both increases and decreases in bleaching observed across grids (57% ΔB_grid_ ≤ 0; 26% ΔB_grid_ > 0). In contrast, bleaching severity was overwhelmingly reduced in 2014-2020 and 2014-2022, with >85% of grids exhibiting ΔB_grid_ ≤ 0 relative to 2014. Consistent with tract-wide tolerance gains during these later events (Fig. 5E–F). However, grids with reduced bleaching remained partitioned into H3 and H4 based on demographic change. In 2014–2015, reduced-bleaching grids were split between persistence (H3) and declining adult density (H4), and this partitioning persisted in later windows despite near-universal reductions in bleaching severity. Sensitization (H2) was common in 2014–2015 but was rare in subsequent comparisons (Fig. 5G–I; Table S4).

To test whether trajectory classes reflect statistically distinct demographic dynamics, we modeled site-level adult colony density for a 13-species assemblage using negative binomial GLM models with a Timepoint × GridState interaction term (β_3_, Table S5; Fig. 5J- L). In this framework (Eq. 4), β_3_ quantifies whether temporal changes in adult colony density differ between persistence (H3) and mortality-filtered (H4) grids. Across all pairwise comparisons, β_3_ was statistically significant, indicating divergence in adult density trajectories between H3 and H4 grid (Table S5). Adult densities remained stable or increased in H3 grids but declined in H4 grids relative to 2014. These patterns were consistent across all adult-size thresholds (W5/S10, W10/S15, and no-threshold) and supported by complementary analyses of juvenile density (Table S5), suggesting adult persistence in H3 grids rather than recruitment-driven replacement alone.

### Baseline predictors of stable persistence and mortality-filtered networks

Stable trajectory networks were defined using only pre-2023 matched-event classifications (2014–2015, 2014–2020, 2014–2022; Table S4). Grids consistently classified as H3 (“Stable H3”, n=12) or H4 (“Stable H4”; n=13), while Stable H2 was rare (n=3) and treated descriptively. Stable H3 and H4 networks showed similar depth and survey effort, but differed in baseline community structure, with Stable H3 networks showing lower densities of weedy adults (MWU p=0.011; Cliff’s δ <0 with 95% CI excluding 0, Table S6). Stress-tolerant/generalist adult densities were also lower on average in Stable H3 networks, though this contrast was weaker and power-limited. Thermal variability was lower on average in Stable H3 networks, although confidence intervals overlapped zero (Table S6).

### Trajectory responses and persistence under the 2023 stress test

To evaluate how trajectory dynamics change under extreme thermal stress, we applied the matched-event framework to the 2014–2023 comparison (Fig. 6A–D). Unlike earlier windows, thermal comparability is limited due to the unprecedented magnitude of the 2023 marine heatwave; results are therefore interpreted as a stress test of the framework rather than a strictly matched comparison.

**Figure 6.**
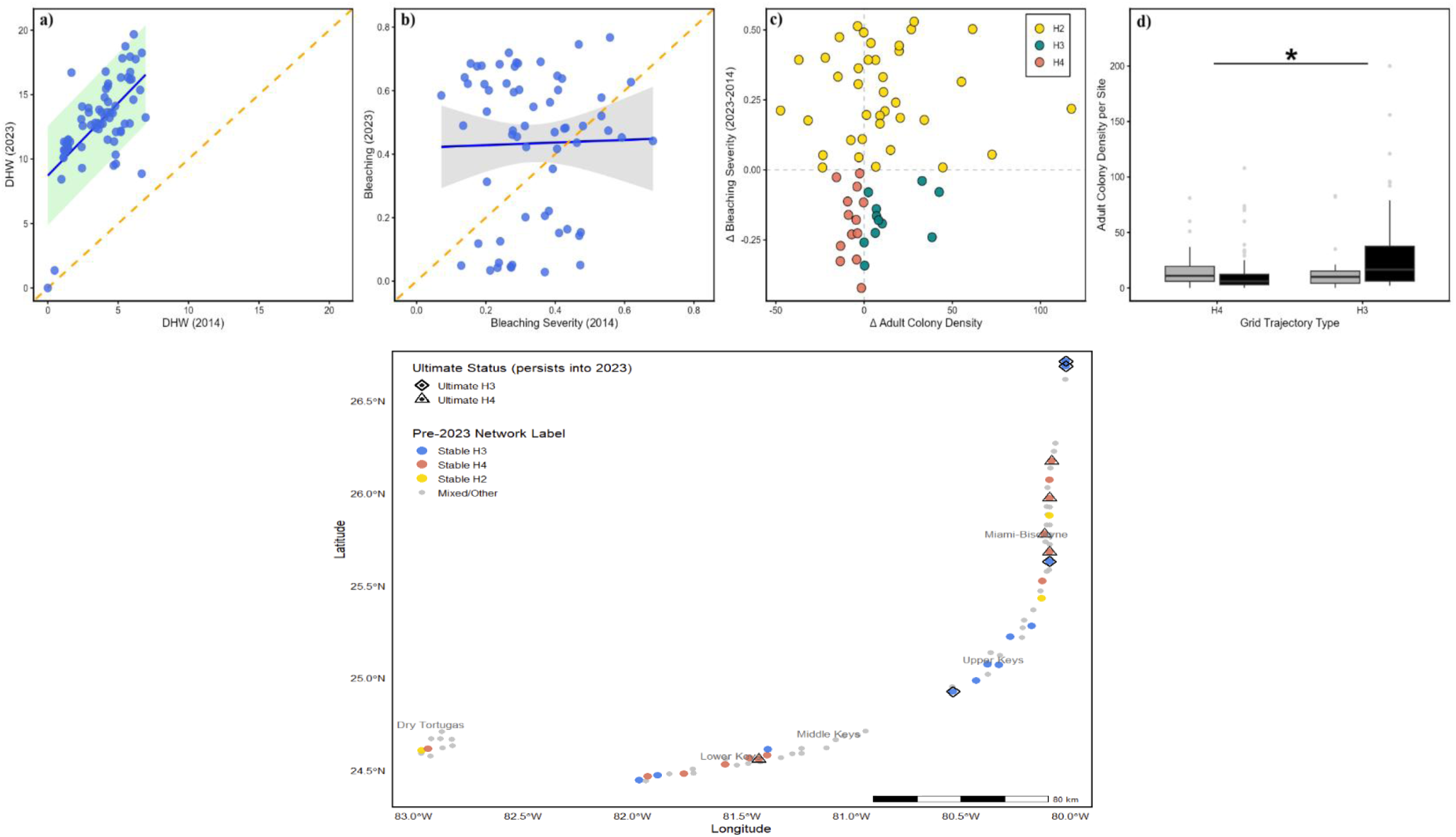
2014–2023 stress test and spatial distribution of stable and ultimate refugia. Trajectory diagnostics for the 2014–2023 window are shown using the matched-event framework (Fig. 5). (a) DHW context, (b) bleaching outcomes relative to 2014, (c) H2/H3/H4 trajectory classification (W5/S10 threshold). (d) Boxplots of CD_adult,site_ for H3 v and H4 grids in 2014 (grey) and 2023 (black). Solid line denotes linear fits (R^2^ = 0.001, p> 0.05) vs 1:1 expectation (dashed). Trajectory divergence calculated as in Fig. 5. (e) Spatial distribution of “Stable” (pre-2023 grids) and “Ultimate” (grids retaining demographic persistence in 2023) networks. See Tables S4, S6-S7.

Under 2023 conditions, bleaching severity increased across most grids relative to 2014 (Fig. 6B–C), and sensitization (H2) became the dominant trajectory. Reduced-bleaching trajectories (H3 and H4), which characterized earlier events, were rare under these conditions. Where reduced bleaching occurred, grid-level responses remained split between persistence (H3) and declining adult density (H4) (Fig. 6C), although sample sizes were limited.

To identify networks that retained their trajectory identity under extreme stress, we defined “ultimate” networks as the subset of Stable H3 and Stable H4 grids (defined using pre-2023 windows) that maintained their respective classifications in 2023 (Fig. 6E; Table S4; Table S7). Because these criteria were stringent, sample sizes were small (Ultimate H3: n = 4; Ultimate H4: n = 5), and results are interpreted using effect sizes and confidence intervals.

Ultimate H3 networks were associated with lower multi-event thermal variability compared to Ultimate H4 networks (Mann-Whitney U, p< 0.05; Table S7). Ultimate H3 networks also experienced lower 2023 DHW on average, although this contrast was uncertain given small sample sizes.

### Species-level decomposition reveals selective drivers of network stability

To identify taxa contributing to divergence between persistence (H3) and mortality-filtered (H4) networks, we decomposed assemblage-level change into species-specific adult density trajectories (Δ ΔCD_grid,adult_, 2014–2023) for 23 coral species (Fig. 7; Table S8).

**Figure 7.**
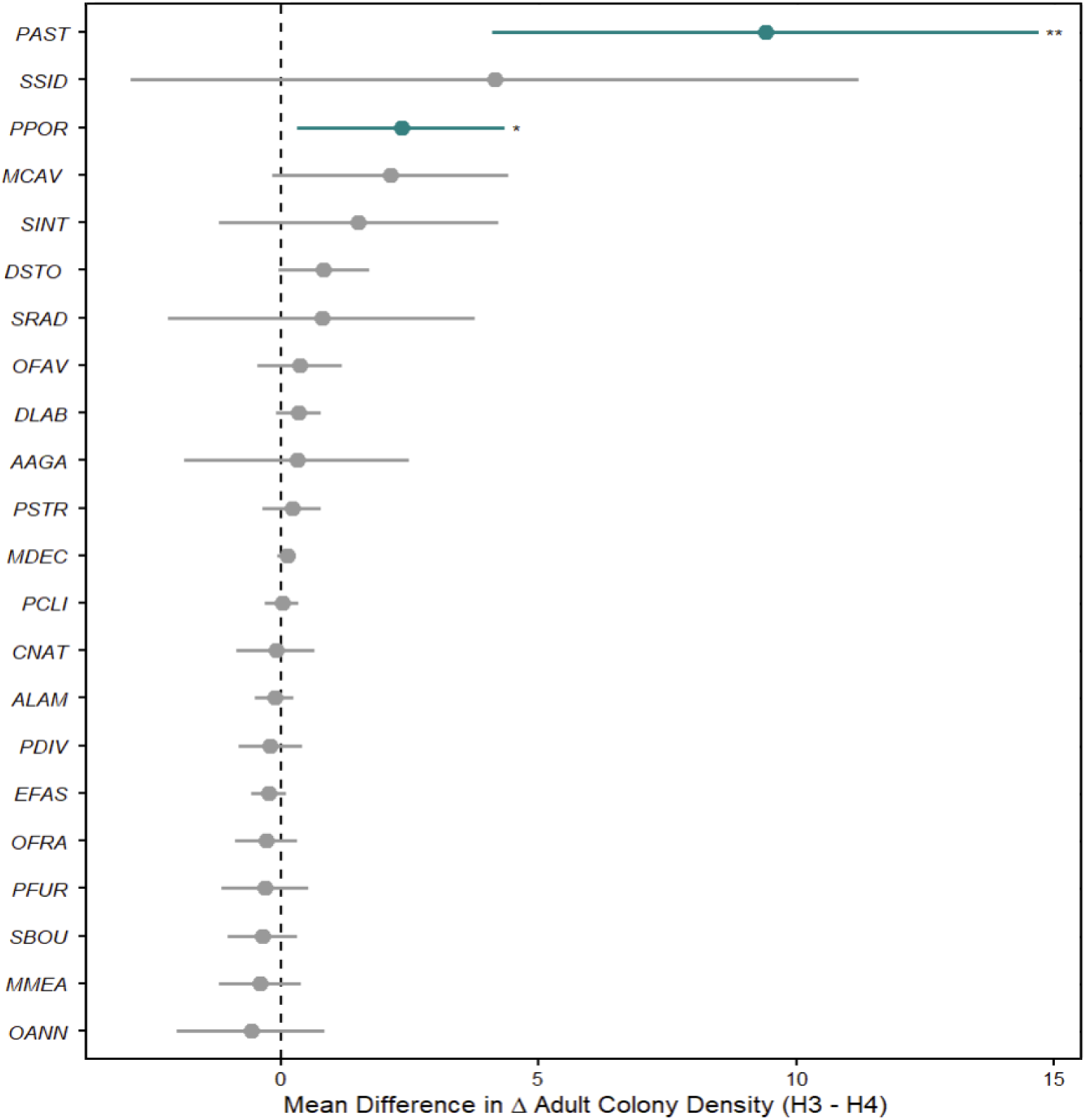
Species-level decomposition of H3–H4 divergence reveals selective contributors to network-scale stability. Forest plot of species-specific shifts in adult colony density (ΔCD_adult,grid_ = H3 - H4; 2014–2023 window; Table S4) comparing H3 (persistence) versus H4 (mortality-filtered) grids. (Welch’s t-test; * p<0.05; ** p<0.01; Tables S8-S9).

The H3–H4 divergence was driven primarily by gains in a small subset of taxa, with the strongest H3 advantages observed in opportunistic/weedy corals (e.g., PAST and PPOR, both significantly higher in H3; Table S8). In contrast, several resilient taxa increased broadly across both grid types (e.g., SSID), contributing to regional persistence without strongly differentiating trajectories.

To ensure these patterns are not driven by trajectory labels that incorporate 2023 outcomes, we repeated the analysis for four focal taxa (PAST, PPOR, MCAV, SSID) using pre-2023 stable network classification (Stable H3 vs Stable H4). Results were consistent: PAST and PPOR retained clear H3 advantages, SSID increased across both network types, and MCAV showed weak directional separation (Table S9).

## DISCUSSION

### Ecological memory is detectable across the Florida Reef Tract during successive marine heatwaves

Across the Florida Reef Tract (FRT), bleaching sensitivity declined from 2014 through 2022, despite moderate–high heat stress in subsequent years. This pattern was supported by both year-specific reaction norms (Fig. 3) and ecological memory interaction models (Table 1). Similar reductions were recovered in a repeatedly surveyed subset of grids and across eleven taxa with sufficient sample size, indicating that the memory signal is driven by spatial sampling variation or a single dominant species.

These findings align with ecological (environmental) memory frameworks, in which species and ecosystem responses to a disturbance depend on recent disturbance history rather than being determined solely by contemporaneous heat stress [4, 8, 9, 11, 28].

### Acclimatization-like persistence vs mortality-filtered tolerance trajectories

Reduced bleaching following repeated heat stress can arise from distinct demographic processes that can produce similar bleaching outcomes [4, 8]. Our trajectory framework resolves this ambiguity by coupling bleaching responses with changes in adult standing stock within thermally comparable grids. Significant Timepoint × GridState interactions indicate divergence between persistence (H3) and mortality-filtered (H4) trajectories (Fig. 5; Table S5), and this pattern is robust across adult-size threshold and juvenile analyses. Stable H3 and H4 networks differed only modestly in baseline conditions, suggesting that divergence emerges primarily through responses to repeated disturbances rather than strongly initial differences. Consistent with this interpretation, ultimate H3 networks were characterized more clearly by lower multi-event thermal variability than by large baseline demographic differences (Table S7). Together, these results are consistent with ecological memory contributing to alternative demographic outcomes under repeated disturbances, with persistence more likely under more predictable exposure regimes.

In H3 networks, reduced bleaching coincides with stable or increasing adult densities, consistent with acclimatization-like persistence [9, 28–31]. In H4 networks, reduced bleaching occurs despite declining adult densities, consistent with mortality filtering via selective loss of susceptible individuals producing a demographically depleted but apparently more tolerant assemblage [12, 13, 16]. These pathways imply different futures: persistence maintains reproductive capacity and structural continuity, whereas filtering can stabilize altered communities while undermining long-term recovery and reef-building function [10, 11, 18].

Most reef networks on the FRT did not maintain a single trajectory across successive events, shifting among sensitization (H2), persistence (H3), and mortality-filtered tolerance (H4) (Fig. 5; Table S4). H2 represents the detrimental side of prior heat exposure, where incomplete recovery or accumulated stress increases subsequent bleaching risk [11, 29]. The trajectory classes capture distinct expressions of history dependence, consistent with previous studies [4, 8], while the rarity of stable networks highlights the limited number of reef systems that maintain consistent responses across repeated heatwaves.

### 2023 stress test: limits, and refugia defined by adult standing-stock persistence

The 2023 MHW provides an extreme stress test of ecological memory. Interaction models reversed sign (Table 1), indicating that prior exposure no longer reduced bleaching sensitivity under extreme heat. Consistent with this result, sensitization dominated the 2014–2023 trajectory comparison, whereas persistence pathways became rare (Fig. 6). Together, these findings suggest that the mechanisms associated with persistence under moderate events do not scale to unprecedented thermal stress, consistent with evidence that resilience declines as recovery intervals shorten and heat stress exceeds historical limits [1, 20, 32, 33].

“Ultimate” refugia, defined as stable H3 or H4 networks that retained their classification in 2-023 (Fig. 6E; Table S7), provide an out-of-sample test of persistence under extreme stress. Persistence (H3) was more strongly associated with lower multi-event thermal variability than with baseline demographic differences, suggesting that persistence under extreme conditions is contingent on exposure history rather than initial state.

Because bleaching prevalence was extremely high during the 2023 event, we define “refugia” by retention of adult standing stock rather than low bleaching, recognizing that bleaching, survival, and longer-term performance can decouple. This distinction is supported by recent studies where mortality and long-term impairment may be driven by thermal maxima and post-bleaching recovery dynamics rather than bleaching prevalence alone [34–36].

Even where adult standing stock persists, extreme events can impose substantial legacy costs through delayed partial mortality and prolonged physiological recovery trajectories, such that visually intact reefs may experience reduced functional performance over multiple years [9, 35, 37]. In this context, “ultimate” refugia should be interpreted not as systems resistant to bleaching but as those that avoid immediate demographic collapse under extreme stress, consistent with broader evidence that persistence and full functional recovery often diverge following successive marine heatwaves, particularly under extreme thermal regimes such as 2023 [4, 9, 35].

### Predictable exposure regime hypothesis

Our results support a bounded ecological-memory framework in which prior exposure can reduce bleaching sensitivity under recurrent, moderate-high heat stress, but only within a limited range of environmental conditions. Because history-dependent responses may depend on the structure of repeated exposure [22] as well as cumulative heat-stress, we interpret our results through a “predictable exposure regime” lens, defined as reduced interannual variance in marine heatwave intensity within a reef network.

Several lines of evidence support this interpretation. Reaction norm and interaction models indicate reduced bleaching sensitivity following prior exposure (Fig. 3; Table 1), while trajectory analyses show that persistence pathways (H3) are maintained across repeated events (Fig. 5). At the network scale, systems that retain demographic persistence under extreme stress (“ultimate” H3 networks) were more strongly associated with lower multi-event thermal variability than with differences in baseline community structure (Table S7). Together, these results suggest that persistence is more likely where successive heatwaves occur within a relatively consistent exposure regime. This interpretation complements recent work showing that bleaching is shaped by multiple aspects of thermal exposure, including variability, temporal structure, and recovery dynamics, rather than on single-threshold metrics alone [21, 22, 38]. Importantly, the variability considered here operates at the interannual scale, in contrast to studies linking reduced bleaching risk to high-frequency (daily to seasonal) temperature variability [23, 39, 40]. These processes are not mutually exclusive. Reef systems may experience short-term thermal fluctuations while remaining embedded within relatively predictable interannual heat-stress regimes [21, 23, 41]. Our results suggest that ecological memory can accumulate under such regimes but weakens when thermal anomalies exceed the historical range of exposure, consistent with the widespread sensitization observed during the extreme 2023 marine heatwave.

### Species-level consequences: selective beneficiaries, altered assemblages, and functional implications

A key question is whether acclimatization-like persistence reflects a community-wide response or is driven by a subset of taxa. Our results indicate that divergence between persistence and mortality-filtered networks is driven primarily by gains in a small number of species, rather than uniform responses across the assemblage (Fig. 7; Table S8). In particular, weedy life-history strategies (e.g., *PAST*, *PPOR*) exhibited the strongest H3 advantages, consistent with their fast growth, early maturation, and high recruitment potential [17, 19]. In contrast, several stress-tolerant taxa (e.g., *SSID, SINT;* and to a lesser extent, *MCAV*), and some weedy taxa (*SRAD, AAGA*) increase in both network types, contributing to regional persistence without strongly differentiating trajectories.

Sensitivity analyses using independent pre-2023 network classifications yielded similar results, confirming that key patterns were not driven by classifications incorporating 2023 outcomes (Table S9). Together, these findings indicate that demographic persistence emerges through ecological reassembly and differential persistence among taxa, rather than community-wide increases in thermal tolerance. [19, 42, 43, 44].

This interpretation aligns with long-term shifts in Florida coral assemblages away from historical dominance by framework-building *Orbicella* and *Acropora* toward weedy and stress-tolerant taxa [18, 19]. Consequently, demographic stability does not necessarily imply functional recovery. Persistence may maintain coral cover and local population viability, but assemblages increasingly dominated by opportunistic taxa are unlikely to rebuild historical reef framework, consistent with projections of declining carbonate production and increasing net erosion under recurrent heat stress [18, 45].

Consequently, refugia are best understood not as places that escape heat stress, but as networks that continue to retain adult standing stock despite repeated disturbance. Persistence therefore reflects a dynamic process of ecological reassembly rather than recovery of historical reef states. This perspective supports conservation strategies that prioritize demographic resilience, environmental heterogeneity, and connectivity under increasing thermal stress [46–49, 44].

### Caveats and future work

Several caveats frame this interpretation. First, stable-network and ultimate-refugia classifications are intentionally stringent, they yield small sample sizes; accordingly we emphasize effect sizes, confidence intervals over strict *significance* thresholds. Second, while our framework distinguishes demographic persistence from mortality filtering, it does not identify the physiological or ecological mechanisms underlying these patterns (e.g., epigenetic memory or symbiont shifts) [10, 35]. Third, randomized sampling necessitates grid-based analyses that reduces bias but limits resolution for rare taxa. Future work combining spatial-network analyses with physiological, demographic and molecular profiling of stable refugia will be needed to distinguish among alternative mechanisms underlying persistence, filtering, and recovery.

### Outlook and implications

As marine heatwaves intensify, reefs face increasing risks of demographic and functional decline and net-erosional states [1, 45], with persistence becoming unlikely without rapid emissions reductions [50, 51]. Our results show that persistence is neither uniform nor guaranteed, but depends on exposure history, demographic trajectories, and species composition. Modern refugia are dynamic systems best defined by retention of adult standing stock rather than low bleaching alone and often reflect selective ecological reassembly rather than recovery of historical reef structure or function. Sustaining reefs will require reducing climate-driven thermal stress maintaining demographic resilience, biodiversity, and network connectivity. These efforts should prioritize not just survival, but the preservation of ecological function and positive carbonate budgets under increasingly extreme thermal regimes [46–49].

## Author’s contributions

T.V.: conceptualization, data curation, analysis, methodology, writing

A.P.C.: data curation, analysis, methodology;

O.M.: data curation, methodology;

Sh.L.D.: writing, and editing;

G.V.: conceptualization, writing and editing;

F.V.: data curation, data analysis;

K.Sh.: data curation, data analysis;

V.B.: conceptualization, methodology, editing;

L.A.M.: conceptualization, formal analysis, funding acquisition, methodology, writing and editing.

All authors gave final approval for publication and agreed to be held accountable for the work performed therein.

## Acknowledgements

The authors thank Dr. Timothy D. Swain and Avery Wallace for insightful discussions that contributed to the development of this work.

