## Supplementary Information for "Bounded ecological memory in coral reefs: divergent demographic pathways under recurrent heatwaves and collapse under extreme thermal stress"

#### SM1. Dataset, data processing and site aggregation

We analyzed data from the Disturbance Response Monitoring (DRM) program, administered by the Florida Fish and Wildlife Conservation Commission under the Florida Reef Resilience Program (FRRP) framework. This dataset comprises surveys of 216,080 coral colonies representing about 45 species, conducted between 2005 and 2023 across the 251 km² Florida Reef Tract (FRT), spanning from Martin County (northern extent: ~27.1° N, 80.4° W) to the Dry Tortugas (southern extent: ~24.3° N, 82.7° W). Surveys followed a randomized design, where each transect within a selected geographic region and reef habitat was sampled only once during the study period [1-3]. This approach does not track individual colonies or fixed sites over time. Instead, it enables repeated sampling of hundreds of coral assemblages across diverse reef habitats and subregions over multiple years, allowing for the identification of broad, system-wide trends that are not tied to conditions observed only at specific reefs. DRM-FRRP surveys consist of two 10×1 m belt transects per site. At each site, two 10 × 1 m belt transects were surveyed to identify colonies to species (for colonies ≥4 cm), measure colony size (height and maximum diameter), and record colony condition including bleaching category (non-bleached, *NB*; pale, *P*; partially bleached, *PB*; bleached, *B*), mortality (recent mortality, *RM*, attributable to bleaching; old mortality, *OM*, attributable to disease, physical damage, or other causes), and disease observations. Species richness per site (8.2 ± 0.28) and colony density per species (5.65 ± 0.23) varied among sites (Supplementary Table S1), reflecting well-documented biogeographic gradients across FRT subregions [4]. Survey effort varied annually, with greater spatial coverage in later years and during periods of elevated thermal anomalies (Supplementary Table S1).

#### SM2. Data exclusion criteria and selection of marine heatwaves.

To relate bleaching severity to heat stress while reducing confounding from other stressors and survey timing, we applied several filters. First, we retained surveys conducted from August through November, when bleaching observations most consistently follow peak seasonal heat stress, and excluded winter/post-bleaching surveys. Second, to isolate acute thermal responses from concurrent biological stress, we excluded colonies exhibiting active signs of disease. This exclusion is conservative but justified because disease incidence was shown to be consistently low (about 8 to 50x lower) relative to bleaching incidence in focal heatwave years (Figure S5). This exclusion also improves comparability across years because disease reporting protocols changed substantially in 2018 following the increased incidence of Stony Coral Tissue Loss Disease (SCTLD) and other coral diseases [5, 6], with new categories and separation of disease-associated mortality that are not directly comparable with earlier surveys. Excluding diseased colonies therefore reduces analytical noise and avoids incomparable disease labels across the 2005–2023 period. Third, we combined observations from both transects at each site and excluded sites where fewer than three species were recorded. Fourth, to evaluate changes in bleaching severity across significant heatwaves, we restricted our analysis to years where: (i) ≥ 30% of the reef sites were exposed to DHW ≥ 4.0, with more than half of those exceeding DHW 4.0; and (ii) at least 150 sites were surveyed. Based on these criteria, the marine heatwaves (MHWs) included in this study were those occurring in 2014 (174 sites; 47 species; 7,850 colonies), 2015 (249; 44, 10,613), 2020 (388; 45; 19,589), 2022 (444; 47; 19,943), and 2023 (606; 48; 33,391, Supplementary Table S1, and Supplementary SM2). The 2014 MHW was used as the reference event for subsequent comparisons of thermotolerance at site and species scales.

Analyses were conducted across multiple hierarchical levels. Site-level models were used to quantify bleaching responses and demographic change, where each site retains a unique heat stress value ($DHW_{site}$) due to survey timing. Grid-level analyses (5 × 5 km) were used for trajectory classification and repeated spatial inference by averaging site-level metrics within each grid-year. Species-level bleaching models were fitted at the colony or species-within-site level, while species-level demographic changes were evaluated at the grid level. This multi-level framework reduces pseudo-replication while preserving biologically meaningful variation across spatial scales.

### SM3. Bleaching severity index computation

We computed bleaching severity following Swain et al. by mapping categorical bleaching states to tissue-affected intervals and using midpoints as scores [7]. The mapping used: non-bleached (0%), pale (1–29%), partially bleached (30–69%), bleached (70–100%), and recent mortality (100%). Species-level severity at site k (Bjk) was computed by weighting each category midpoint by the proportion of colonies in that category for species j at site k. Importantly, any occurrence of recent mortality (RM) was treated as complete tissue loss (100% affected), irrespective of the proportion of the colony exhibiting recent mortality. Thus, the presence of RM contributes a value of 1.0 to the bleaching severity calculation, effectively penalizing Bjk to the maximum possible value for that category. This mathematical penalty reflects the biological interpretation that any recent mortality observed during bleaching surveys represents irreversible loss of coral tissue that is not comparable to partial bleaching alone and ensures that lethal thermal stress is consistently recorded as a more severe response than severe, yet potentially survivable, bleaching.

Site-level assemblage severity ($B_{site}$) was computed as the unweighted mean of Bjk across species present, avoiding disproportionate influence of dominant taxa and enabling cross-site comparisons across a highly heterogeneous tract. While some bleaching severity metrics weigh bleaching susceptibility scores by the relative abundance of each taxon [8, 9], and others focus on the few most abundant coral taxa with abundance-weighted susceptibility scores [10], we chose an unweighted assemblage-level index, also used in other studies [11]. This avoids bias toward dominant species and facilitates valid cross-site comparisons through multiple bleaching years in a system where species composition varies substantially over the large spatial scale of the FRT and temporal scale (2014 - 2023).

**SM4. Heat stress extraction and spatial alignment**

Cumulative thermal stress was quantified using NOAA Coral Reef Watch's (CRW) Version 3.1 Degree Heating Weeks (DHW) at 5 km (0.05°) resolution (v3.1) [12, 13]. This product provides global heat stress metrics at a 5 km (0.05°) resolution, and measures both the magnitude and duration of heat stress above the maximum monthly mean (MMM) climatological sea surface temperature for a given location, where daily positive temperature anomalies ≥ 1 °C above MMM are accumulated over a rolling 12-week window, expressed as °C-weeks. DHW is a robust predictor of coral bleaching severity, with established thresholds linking heat stress intensity to bleaching and mortality risks [14, 15]. To align biological survey data with environmental stress, DRM-FRRP survey sites were spatially mapped to the nearest 5 km grid cell center (lat_pixel_center ±0.025°). For sites falling exactly on a grid boundary, we followed a standardized convention assigning the coordinate to the southern or eastern adjacent cell. To capture the thermal exposure most physiologically relevant to the bleaching response, we extracted the maximum DHW value observed within a 14-day window preceding the survey date for each site. This window accounts for the potential lag between peak thermal accumulation and the visible manifestation of bleaching or mortality in the field and resulted in unique DHW values associated with each site for a given location and survey month ($DHW_{site}$).

Sites that overlapped with land-masked pixels (DHW code = 253) were excluded from the analysis. For spatial comparisons, site-level bleaching severity values ($B_{site}$) were averaged within each 5 x 5 km grid cell to derive a grid-level bleaching metric ($B_{grid}$) per year calculated as:

$$\mathrm{B}_{grid} = \frac{1}{n}\sum_{i=1}^{n} B_{site,i}$$

where $n$ is the number of sites surveyed within the cell and $B_{site,i}$ is the bleaching severity at site $i$. Due to DRM-FRRP's randomized survey design, this means that when comparing the same grid cell across years, $\mathrm{B}_{\mathrm{grid}}$ represents the spatial average of randomly surveyed sites, species compositions and individual colonies within that cell (Supplementary Table S1).

Additionally, because surveys occur over a seasonal window (August–November), sites mapped to the same grid cell within a given year can experience different thermal exposure values depending on survey timing, resulting in multiple $\mathrm{DHW}_{\mathrm{site}}$ values within a grid-year. Thus, when comparing heat stress exposure at grid cell-level for specific years, the $\mathrm{DHW}_{\mathrm{grid}}$ was computed by averaging across all the $\mathrm{DHW}_{\mathrm{site}}$ values within each grid-year (two-step averaging), thereby integrating site-specific exposure differences arising from survey timing.

*Statistical Evaluation of Thermal Differences.* Differences in $\mathrm{DHW}_{\mathrm{grid}}$ among the focal years were evaluated using non-parametric tests, as the $\mathrm{B}_{\mathrm{site}}$/ $B_{grid}$ values exhibited non-normal distributions. This evaluation was performed at two distinct spatial scales to match our biological analyses. For the full Florida Reef Tract, where grid-level observations were treated as independent across sampling years, overall differences were assessed using Kruskal-Wallis tests, followed by post-hoc pairwise Wilcoxon rank-sum tests with a Bonferroni correction. Conversely, for the subset of repeatedly surveyed grid cells where spatial observations were paired longitudinally through time, differences in thermal exposure were evaluated using Friedman tests, followed by post-hoc pairwise Wilcoxon signed-rank tests with a Bonferroni correction (see Figure 3, Supplementary Figure S2, and Supplementary Table S3).

**SM5. Two-step averaging and true-zero handling (pseudo-replication control)**

Pairwise grid trajectories between 2014 and subsequent survey years (2015, 2020, 2022, 2023) were classified using a two-step averaging framework to avoid colony level pseudo replication (described in SM4 for DHW). Specifically, sites mapped to a specific grid-year may have different DHW values due to the survey time, so site-level values were averaged to calculate a mean DHW value for a given grid-year. For bleaching, colony observations were aggregated to $Bjk$ within site-year, then to $B_{site}$, and finally averaged across sites in each grid-year to obtain $B_{grid}$. For demography, colony counts were aggregated to site-year densities and then averaged to grid-year densities. For species-specific demography, we retained “true zeros” by explicitly including surveyed sites where a focal species (or demographic category) was absent, ensuring that grid-level means reflect both presence and absence rather than conditioning on occurrence. Colony density was calculated as colonies per site. Since a site used two 10 x 1 belt transects, colony density was calculated as the number of colonies per 20 square meters. This approach preserves the randomized sampling design while enabling repeated spatial comparisons using grid cells as the consistent unit. Grids with no sites surveyed in a given year were classified as ND_missing (NA) and excluded from biological trajectory assignment.

### SM6. Reaction norm GLMs and plotting conventions

We examined shifts in bleaching severity at the level of the coral assemblages (site- or grid-level) and at the level of individual species across 5 MHWs (2014, used as reference, 2015, 2020, 2022 and 2023), using a binary tipping-point definition of severe bleaching after Hughes et al. (2019) [14], which captures a meaningful ecological threshold: when severe bleaching prevalence is low, recovery is more likely, whereas widespread severe bleaching increases the risk of mass mortality and structural collapse. Shifts in bleaching severity at the level of coral assemblages were calculated at two distinct ecological scales using two different metrics. First, we modeled the probability of severe bleaching ($P_{severe}$) in each site-year as a function of heat stress (DHW) using Generalized linear models (GLM; Eq. 1) adapting the "Ecological memory" framework [14] and applied to the entire FRT (n = 174, 249, 388, 444, and 606 sites for 2014, 2015, 2020, 2022 and 2023, respectively; statistical significance assessed using Wald z-tests on the $\beta_1$ coefficient for each year-specific model Supplementary Table S2**;** Figure 3). However, there is a well described spatial heterogeneity across the region which could mask high variability in exposure and stress response including mortality due to regional differences in water temperature and in species diversity and abundance. Additional variability could be introduced by the DRM-FRRP random surveying structure, where sites are randomly surveyed every year, so that a stable colony density could just be sampling error (i.e., surveying high-density sites in later years). To account for this, we subsequently examined a smaller sub-set of 48 repeatedly surveyed 5 x 5 km grid cells (n= 107, 153, 190, 225, 253 sites for 2014, 2015, 2020, 2022 and 2023, respectively; statistical significance was assessed using Wald z-tests on the $\beta_1$ coefficient for each year-specific model Supplementary Table S2; Figure S3). Second, we determined the change in bleaching severity ($\Delta B_{grid}$) in pairwise comparisons of 5 x 5 km grid cells exposed to DHW intensities in the current year comparable to those in the 2014 MHW (Eq. 3a in methods, see matched-event analysis section below). Finally, to determine if assemblage-level trends were conserved at the taxonomic level, we utilized GLMs to model colony-specific bleaching severity (Bjk) for the 11 species with sufficient sample sizes across the FRT. Most of the 11 examined species exhibited low abundance at many sites resulting in very wide confidence intervals and non-significant results due to small sample sizes. Most other species exhibited low abundance at many sites resulting in very wide confidence intervals and non-significant results due to small sample sizes. $P_{severe}$ was modeled as a binomial response variable (0 = non-severe, $B_{site}$ <0.3; 1 = severe, $B_{site}$ ≥0.3], Eq. 1 in methods. Statistical significance was assessed using Wald z-tests on the $\beta_1$ coefficient for each year-specific model Supplementary Table S2; Figure 4). All models were fitted using the *glm* function in R (R Core Team, 2023).

<u>Plotting conventions and Interpretation:</u>

To accurately represent the thermal regime experienced by coral assemblages in each year, we adopted a hybrid visualization approach for the reaction norms. Within the observed DHW range for each specific survey year (e.g., 0–7 DHW for 2014; 0–20 DHW for 2023), reaction norms are plotted as solid lines bounded by ribbons representing ± 1 Standard Deviation. This convention visualizes the dispersion of the empirical data and the magnitude of the trajectory shift between years. Portions of the curve extending beyond the observed thermal range (extrapolations) are rendered as faint dashed lines without ribbons. These extrapolations are provided solely to visualize theoretical trajectory shifts and are explicitly labeled to prevent over-interpretation of unobserved thermal states.

Formal statistical significance of the relationship between heat stress and bleaching within each year was assessed using Wald z-tests on the $\beta_1$ coefficient for each year-specific model ($p<0.05$). The visual divergence of the modeled reaction norms and their corresponding dispersion ribbons at standardized thermal stress levels (e.g., DHW = 5, 6, 7) illustrates the magnitude of the temporal shift in thermal tolerance relative to the 2014 baseline.

**SM7. Ecological Memory interaction models.**

To test whether initial exposure to the 2014 baseline MHW modifies bleaching sensitivity in subsequent events, we applied the 'Ecological Memory' modeling framework established by Hughes et al. [16, 14]. The probability of severe bleaching at the site level was modeled using binomial generalized linear models (GLMs) with a logit link function, where severe bleaching was binary-coded (1 when $B_{site}$>= 0.3; 0 otherwise. For each pairwise temporal window (2014 –2015, 2014 –2020, 2014 –2022 and 2014 –2023), we compared two models. The 'Year Alone' model predicts severe bleaching as a function of the cumulative heat stress experienced during the current anomaly ($DHW_{current}$, Eq. 1 in methods). The 'Interaction' model incorporated a multiplicative interaction term described in Eq. 2 (see methods). Following Hughes et al. [16], the main effect for historical heat stress ($DHW_{2014}$) was explicitly omitted from the Interaction model. This ensures a shared intercept at 0 °C-weeks, forcing the model to purely evaluate how historical baseline exposure alters the rate (slope) of the current bleaching response. Model performance was evaluated using McFadden's pseudo-$R^2$, which measures the proportional improvement in log-likelihood relative to a null model and is appropriate for logistic regression. Model comparison between the 'Year Alone' and 'Interaction' formulations was based on ΔAIC, where a lower AIC (Akaike Information Criterion) indicates better support after penalizing for model complexity (Table 1). Together, these metrics assess both goodness-of-fit and parsimony, ensuring that any improved fit reflects a meaningful ecological signal rather than mathematical overfitting.

**SM8. Matched-event analysis and ND_thermal envelope definition**

To isolate the influence of prior thermal history on subsequent bleaching responses, we conducted a matched-event analysis across repeatedly surveyed grid cells. Reef sites aggregated into 5 × 5 km grid cells were paired across years relative to the 2014 marine heatwave (MHW), which was used as the baseline event. To maximize sample size given lower survey coverage in earlier years, we conducted pairwise comparisons between 2014 and each subsequent MHW year: 2014 - 2015 pair (n = 63 grids encompassing 134 and 180 sites in 2014 and 2015, respectively), 2014- 2020 pair (n = 68 grids with 155 and 251 sites in 2014 and 2020, respectively), 2014- 2022 (n = 71 grids with 160 and 290 sites in 2014 and 2022, respectively), and 2014- 2023 (n= 74 grids with 165 and 367 sites in 2014 and 2023, respectively, Supplementary Table S1).

Thermal comparability between years was evaluated using the relationship between baseline heat stress ($DHW_{2014}$) and heat stress in the focal year ($DHW_{current}$). A linear regression was fitted for each pairwise comparison, and a regression envelope (±1.5 standard deviations of the residuals) was used to identify thermally typical grid cells. Grids falling within this envelope were retained for biological classification, whereas grids outside the envelope were classified as ND_thermal (thermally atypical) and excluded from downstream analysis. Grids lacking survey data in one of the paired years were classified as ND_missing. This approach controls for differences in thermal magnitude between events, while accounting for spatial heterogeneity in heat exposure, enabling comparison of bleaching severity under broadly comparable thermal conditions. Biological trajectory classes were then defined for thermally typical grids based on (i) their change in bleaching severity ($\Delta B_{grid}$) which distinguishes increased versus reduced bleaching severity relative to 2014, and (ii) their change in adult colony density ($\Delta CD_{adult,grid}$), which captures persistence or decline in adult standing stock, allowing to identify patterns consistent with physiological acclimatization from those consistent with mortality-driven filtering. Using Eq. 3 (see methods), we evaluated three non-mutually exclusive processes driving $\Delta B_{grid}$ dynamics: *Sensitization* (H2), where bleaching severity increases due to incomplete recovery from prior stress ($\Delta B_{grid} > 0$). Reduced bleaching ($\Delta B_{grid}$ =< 0), may arise through two distinct processes: *acclimatization/persistence-like tolerance (H3),* defined as reduced bleaching accompanied by stable or increasing adult colony density ($\Delta CD_{adult,grid} \geq 0$) due to environmental memory of prior exposure and physiological adjustment, and *mortality-filtered tolerance (H4)*, defined as reduced bleaching accompanied by declining adult colony density ($\Delta CD_{adult.grid} < 0$) due to higher mortality of heat-sensitive colonies in prior heat stress events, leaving a more resistant assemblage. These patterns cannot be distinguished using bleaching alone; therefore, we incorporated changes in adult colony density ($\Delta CD_{adult,grid}$; see Demographic modeling section) to separate patterns consistent with demographic persistence from mortality-driven filtering.

**SM9. Demographic modeling and threshold sensitivity**

Colony density was evaluated at the site level for a pooled assemblage of thirteen abundant and ecologically important taxa in the Florida Reef Tract (FRT), selected to ensure sufficient sample size for robust inference. To isolate the demographic signature of acclimatization from post-disturbance recruitment dynamics, analyses were restricted to adult colonies, which serve as a proxy for individuals that persisted through earlier marine heatwaves [17]. Including juvenile colonies could confound interpretation by masking adult mortality with recruitment-driven increases in density. Focusing on adults also increases the likelihood that counted colonies were present through earlier heatwaves, rather than being dominated by post-event recruits.

Colony size (maximum diameter, cm) was used as a proxy of age and reproductive status, with species assigned to life-history strategy groups reflecting differences in growth rate and maturation size, with weedy species typically exhibiting faster growth rates, earlier reproductive age and smaller mature adult sizes than stress-tolerant species [18] using data accessed via the Coral Trait Database [19] (Figure S4). Stress-tolerant and generalist life-history strategies [*Siderastrea siderea* (SSID), *Montastraea cavernosa* (MCAV), *Stephanocoenia intersepta* (SINT) Dichocoenia stokesii (DSTO), Diploria labyrinthiformis (DLAB), Pseudodiploria strigosa (PSTR), Colpophyllia natans (CNAT); Generalists: Orbicella annularis (OANN), Orbicella faveolata (OFAV)], were grouped together (ST/Gen), while weedy taxa [*Porites astreoides* (PAST), *Agaricia agaricites* (AAGA), *Porites porites* (PPOR), and *Siderastrea radians* (SRAD)] formed a separate group (W). Thus, colonies > 10 cm were identified as experienced adults for stress tolerant species due to their slower growth rate of about 0.5 – 1 cm/ year, while colonies > 5 cm were defined as adults for weedy species, due to their faster growth rate of about 1- 1.5 cm/ year [18]. We also evaluated two additional thresholds; Weedy adults >= 10 cm and Stress-tolerant/Generalist adults >= 10 cm and, no threshold, allowing for all colonies to be included in the delta-colony density metric (Figure S4). Site-level adult colony density (including true zeros) was first calculated and then averaged across sites within each grid cell and year to obtain grid-level adult density ($CD_{adult,grid}$) following the two-step averaging framework (*Supplementary SM4 and SM5*). True zeros were assigned to surveyed sites lacking adult colonies before aggregating to grid-year means. To test whether colony density trajectories differed between trajectory classes, we fit negative binomial generalized linear models (GLMs) to site-year adult colony density using a Timepoint × GridState interaction (Density ~ Timepoint × GridState) with a log link (implemented using glm.nb in the MASS package in R, Eq. 4 in methods). GridState (H3 vs H4) was assigned at the grid level (Supplementary Table S4), and site-year observations inherited their grid's state. The interaction term tests whether temporal changes in adult density differ between trajectory classes, rather than reflecting baseline differences alone. Full model coefficients, confidence intervals, and diagnostics are reported in Supplementary Table S5.

Robustness of demographic inference was evaluated across adult size thresholds and through complementary analyses of juvenile colony density (colonies below adult size thresholds), allowing us to distinguish adult standing-stock persistence from recruitment-driven replacement.

**SM10. Stable pre-2023 labels: computation and statistics**

Stable network labels were derived from Table S8 pre-2023 trajectory assignments (2014–2015, 2014–2020, 2014–2022) using the following criteria: For each grid cell, we counted the number of times it was classified as H2, H3, or H4 across these windows. A grid was labeled Stable H3 if it was classified as H3 in at least two of the three windows and was never classified as H2 or H4; Stable H4 if classified as H4 in at least two windows and never H2 or H3; and Stable H2 if classified as H2 in at least two windows and never H3 or H4. Grids not meeting these criteria were labeled Mixed and excluded from baseline predictor inference. Baseline predictors were computed from 2014 surveys only using two-step averaging with true zeros. For baseline thermal metrics, $DHW_{grid}$ values for each MHW year were computed by averaging $DHW_{site}$ across sites in each grid-year; thermal variability metric Standard Deviation (SD) summarized the distribution of DHW across the four pre-2023 MHWs. Baseline comparisons between Stable H3 and Stable H4 used Mann-Whitney U tests (equivalent to the Wilcoxon rank-sum tests), and Cliff's δ (95% CI) was computed as a distribution-free effect size (Supplementary Table S6). Interpretation emphasized a priori focal predictors (baseline adult densities and thermal variability) because Supplementary Table S6 is intended as contextual characterization rather than a multi-hypothesis discovery screen. "Ultimate" grids (i.e., Stable H3 or Stable H4 based on pre-2023 trajectories) that retained their trajectory classification in the 2014–2023 comparison were also evaluated (Supplementary Table S7), and group comparisons between ultimate H3 (4 grids) and ultimate H4 (5 grids) were compared using similar statistical tests described for stable grids above.

**SM11. Species decomposition and out-of-sample sensitivity**

For species-level decomposition, adult density for each of 22 taxa was computed at the site-year level with true zeros for surveyed sites lacking the species, then aggregated to grid-year means via two-step averaging (SM4 and SM5). Grid membership (H3 vs H4) was held fixed using the 2014–2023 trajectory classifications from Supplementary Table S4; this analysis is interpreted as decomposition of community-level divergence rather than prediction. Species-specific density change was calculated as $\Delta CD_{grid,adult} = \Delta CD_{grid,adult\ ,2023} - \Delta CD_{grid,adult\ ,2014}$ (Eq.3b), and differences in ΔCD between H3 and H4 grids were evaluated using Welch's two-sample t-tests (unequal variances), reporting mean differences and 95% confidence intervals (Supplementary Table S8).

To ensure that key species-level patterns are not dependent on trajectory labels that incorporate 2023 outcomes, we performed an out-of-sample sensitivity analysis using the pre-2023 stable network labels described in the previous section. We restricted this analysis a priori to four focal taxa representing distinct ecological roles and key patterns from the main species decomposition: *PAST* and *PPOR* as opportunistic drivers, *MCAV* as a stress-tolerant framework builder, and SSID as a broadly resilient taxon. $\Delta CD_{grid,adult}$ was recomputed for four focal taxa (*PAST, PPOR, MCAV, SSID*) and compared between Stable H3 and Stable H4 using Welch tests (Supplementary Table S9).

**Figure S1. Full-year bleaching–heat stress relationships across the 2005–2023 record**

Relationships between site-level bleaching severity ($B_{site}$) and heat stress ($DHW_{site}$) are shown for all 19 survey years, providing full temporal context for selection of focal marine heatwave (MHW) events (2014, 2015, 2020, 2022, 2023). Bleaching severity ($B_{site}$) was computed as the unweighted mean of species-level severity (Bjk) within each site, and $DHW_{site}$ represents the CRW maximum Degree Heating Weeks (°C-weeks) within a 14-day window preceding each survey. Points represent individual sites, spanning heterogeneous spatial sampling across the Florida Reef Tract (FRT). The figure demonstrates that strong positive relationships between heat stress and bleaching are largely restricted to major MHWs, justifying the focus on selected events for subsequent ecological memory and trajectory analyses. See descriptive statistics and linear regression significance in Table S1.

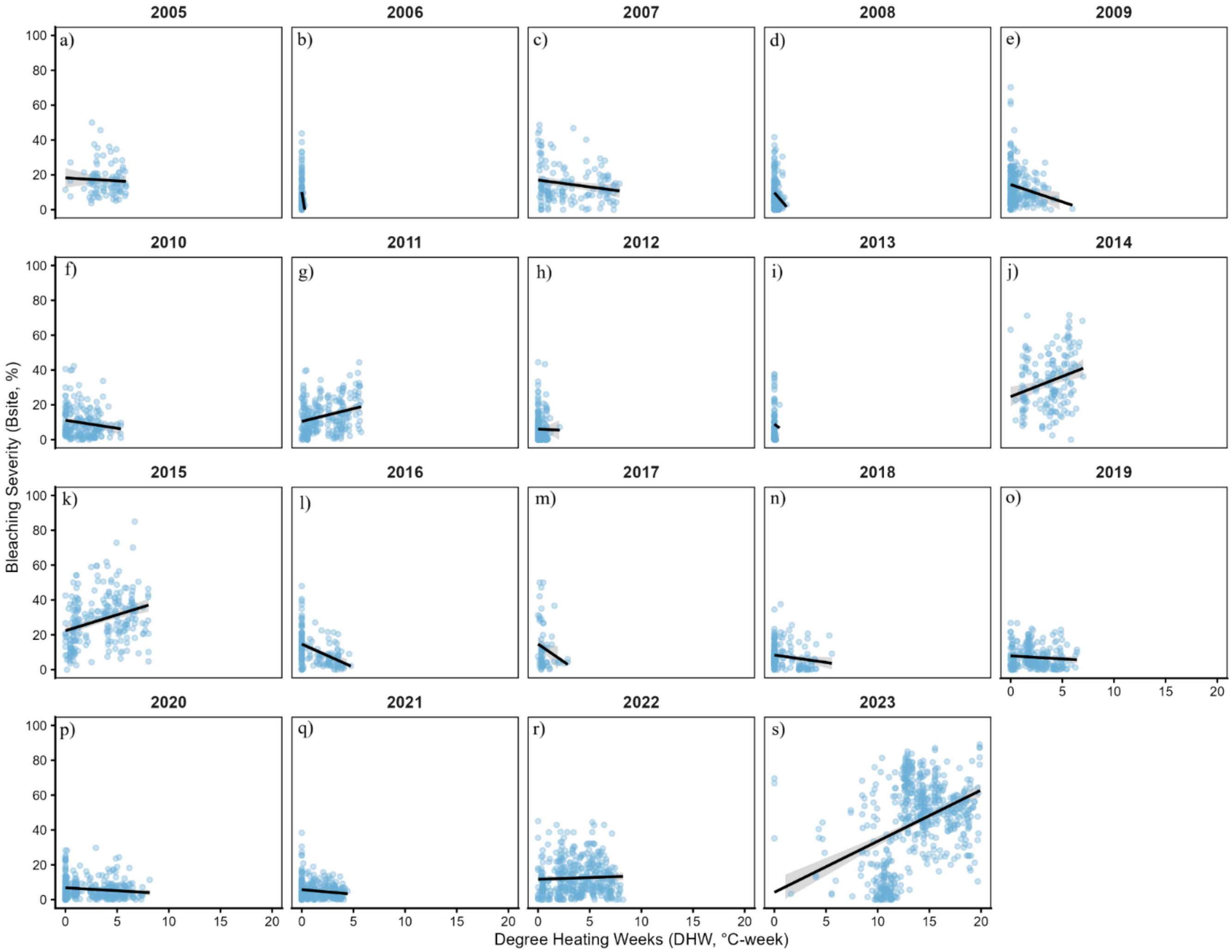

**Figure S2. Reaction norms in repeatedly surveyed grid cells (paired spatial subset)**

Severe-bleaching reaction norms restricted to sites within repeatedly surveyed 5 × 5 km grid cells (n = 48) across 2014, 2015, 2020, 2022, and 2023 (107, 153, 190, 225, 253 sites, respectively). (a) Site-level binomial GLMs (as in Figure 3, Eq.1) were fitted within this spatially constrained subset to reduce the influence of spatial heterogeneity and shifting site composition across the years. Similar shifts in bleaching sensitivity as the full dataset are observed, supporting the interpretation that ecological memory signals are not artifacts of uneven spatial sampling but reflect consistent changes within recurrently sampled reef areas. Solid lines with ±1 SD shading are shown only within the observed DHW range for each year, dashed lines display extrapolated model fits beyond observed values (Table S2) (b) Distributions of CRW site-level DHW among surveyed sites for each focal grid-year. The DHW distributions in 2015 and 2022 did not differ significantly from the 2014 baseline; however, thermal stress in 2020 was significantly lower, and 2023 was significantly higher than all other years (Friedman, Bonferroni-adjusted Wilcoxon tests; Table S3). (c) Spatial distribution of grids repeatedly surveyed in 2014, 2015, 2020, and 2022 (n=50; n= 48 grids when 2023 is included; Table S1).

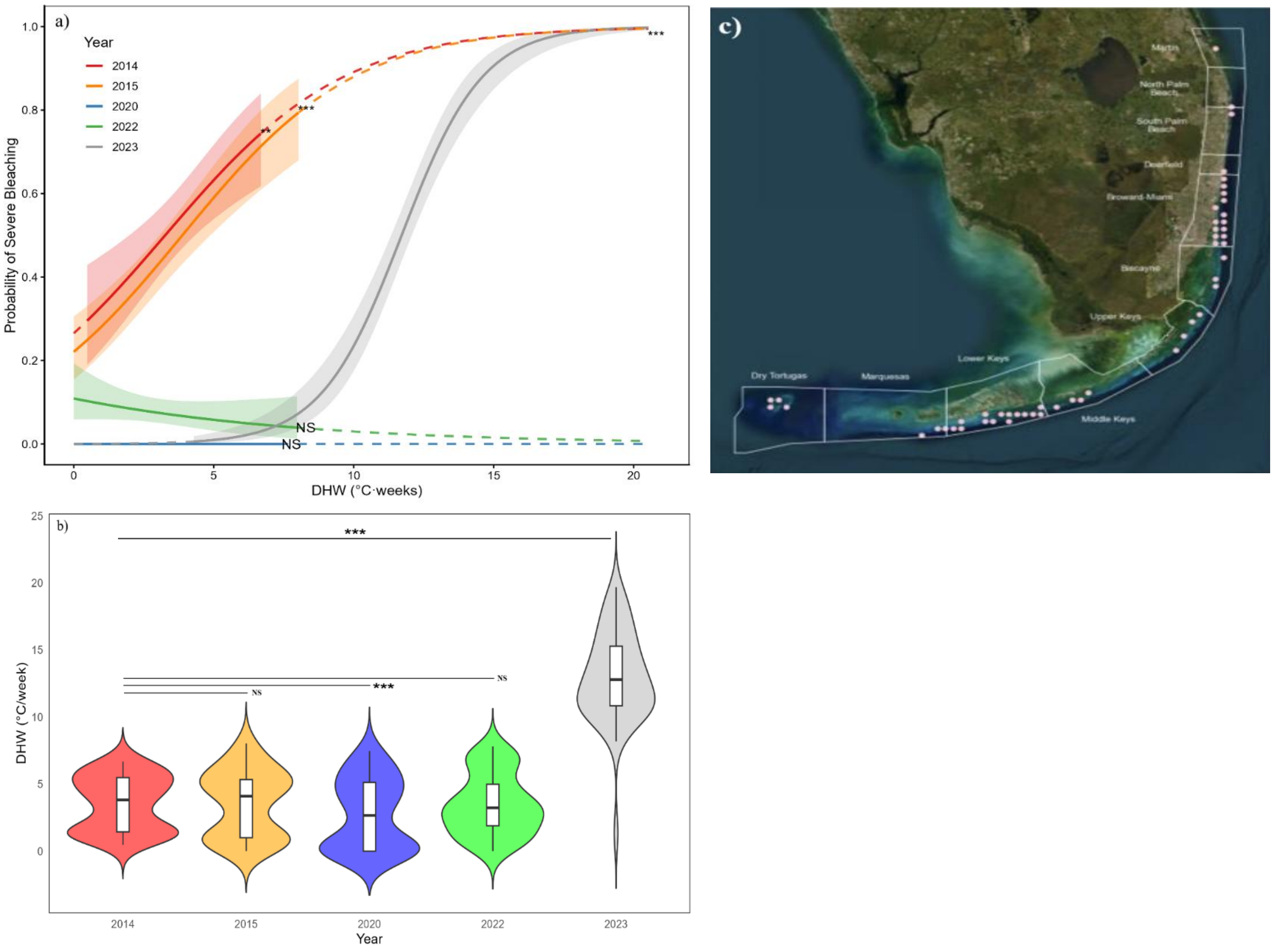

**Figure S3. Sensitivity analysis of reaction norms using grid-level aggregation**

Sensitivity analysis of severe-bleaching reaction norms derived from grid-level (5 × 5 km) averages rather than site-level data for 2014, 2015, 2022 and 2023 (80, 91, 136, 148 grids, respectively). Bleaching severity ($B_{grid}$) and heat stress ($DHW_{grid}$) were calculated using a two-step averaging approach with true zeros retained (Supplementary SM4). Binomial GLMs relating probability of severe bleaching ($B_{grid} \geq 0.30$) to DHW were fitted as in the main analysis (Figure 3; Eq. 1, Wald-z test). Compared with site-level models (Figure 3), grid-level aggregation reduces sample size and increases uncertainty (wider confidence intervals) but preserves the direction and magnitude of temporal shifts in bleaching sensitivity, confirming that tract-wide patterns are robust to spatial aggregation and not driven by site-level sampling variability (Solid lines, dashed lines and shading as in Figure 3 and Figure S2). See descriptive and model statistics in Table S1 and Table S2.

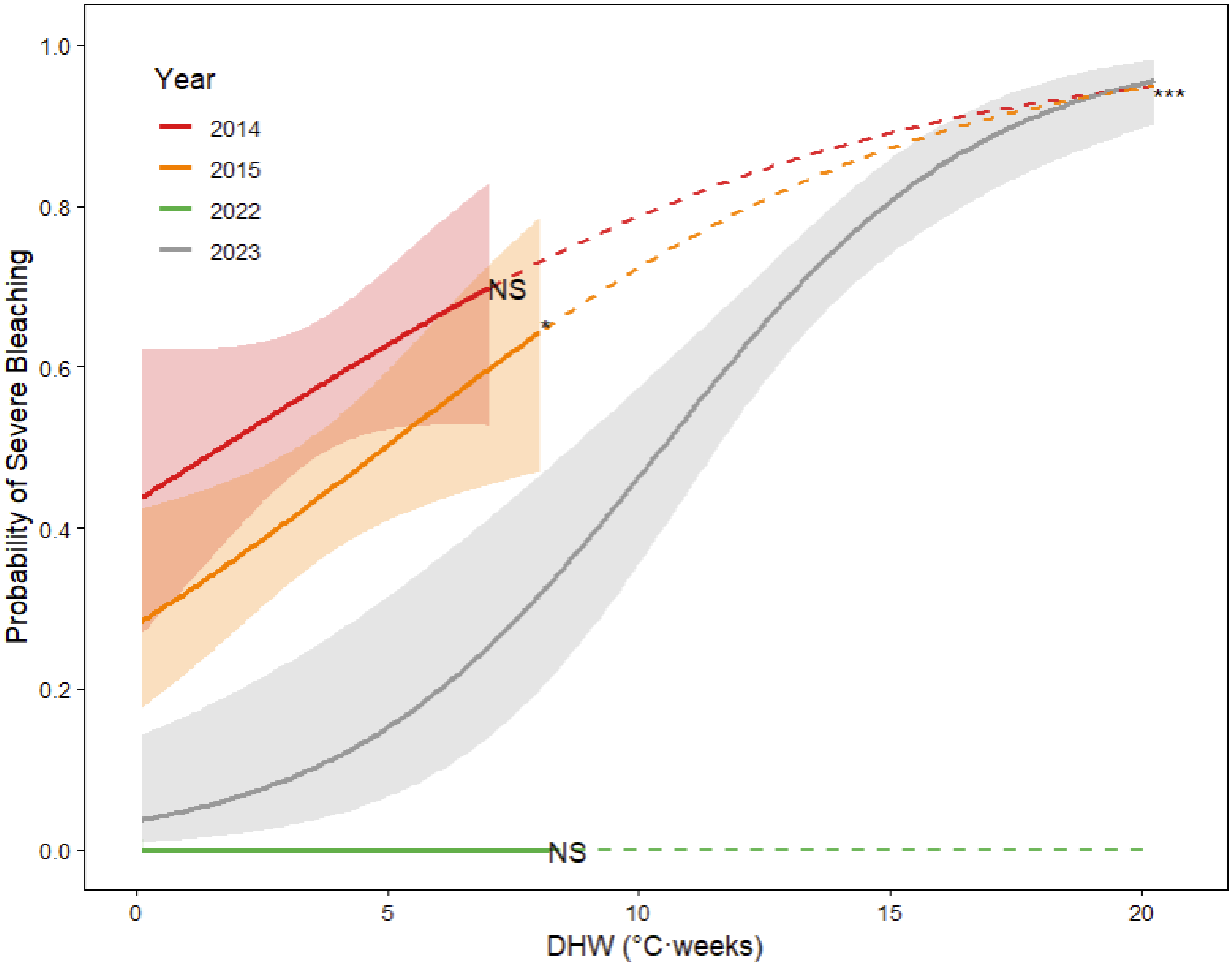

**Figure S4. Colony size-frequency distributions supporting adult size thresholds**

Frequency distributions of colony diameter (cm) for 22 coral species in 2014 and 2023, used to justify adult size thresholds for demographic analyses. Size distributions are shown for individual taxa and illustrate differences in growth strategies between life-history groups. Weedy species exhibit greater representation in smaller size classes, whereas stress-tolerant/generalist taxa are skewed toward larger sizes. These patterns support the use of life-history–specific thresholds (Weedy≥ 5 and Stress-tolerant/ Generalist ≥10, W5/S10, or Weedy≥ 10 and Stress-tolerant/ Generalist ≥15, W10/S15) to define adult colonies and justify the separation of adult vs juvenile demographic signals in trajectory classification and GLM analyses. See model statistics for different size thresholds in Table S5.

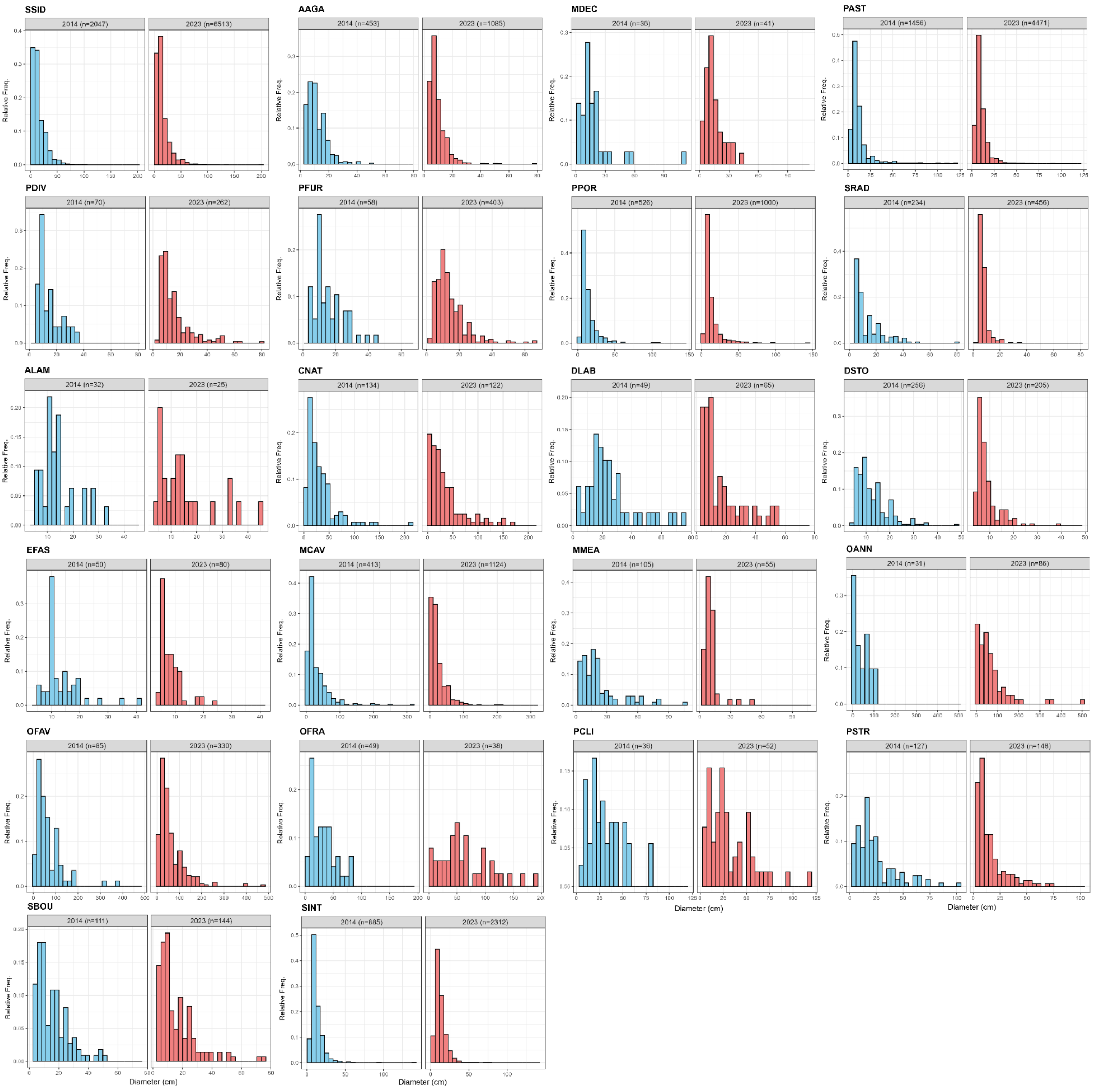

**Figure S5. Proportion of affected colonies by condition (Bleaching vs Disease) across years**
Temporal dynamics in colony condition across survey years (2014–2023), showing the proportion of colonies affected by bleaching, disease, or both. Values are reported as percentages of total surveyed colonies per year. “Bleaching condition” includes PB (partially bleached) and B (bleached) and P(Pale) but excludes N (normal) colonies. “Disease condition” was defined as all diseases observed during survey except DC (discolored) colonies. “Bleaching + Disease” condition includes colonies that are display both conditions (Bleaching AND Disease) (DRM-FRRP dataset, https://coraldrm.org/DataDownload/DrmDataDownload, accessed on 5/12/2024). The figure demonstrates that bleaching dominates observed stress responses during MHW years, while disease incidence remains comparatively low, supporting the decision to exclude diseased colonies from analyses to reduce confounding and maintain comparability across years. This exclusion ensures comparability, as new disease reporting protocols introduced in 2018 rendered earlier surveys incompatible (Supplementary Material SM2).

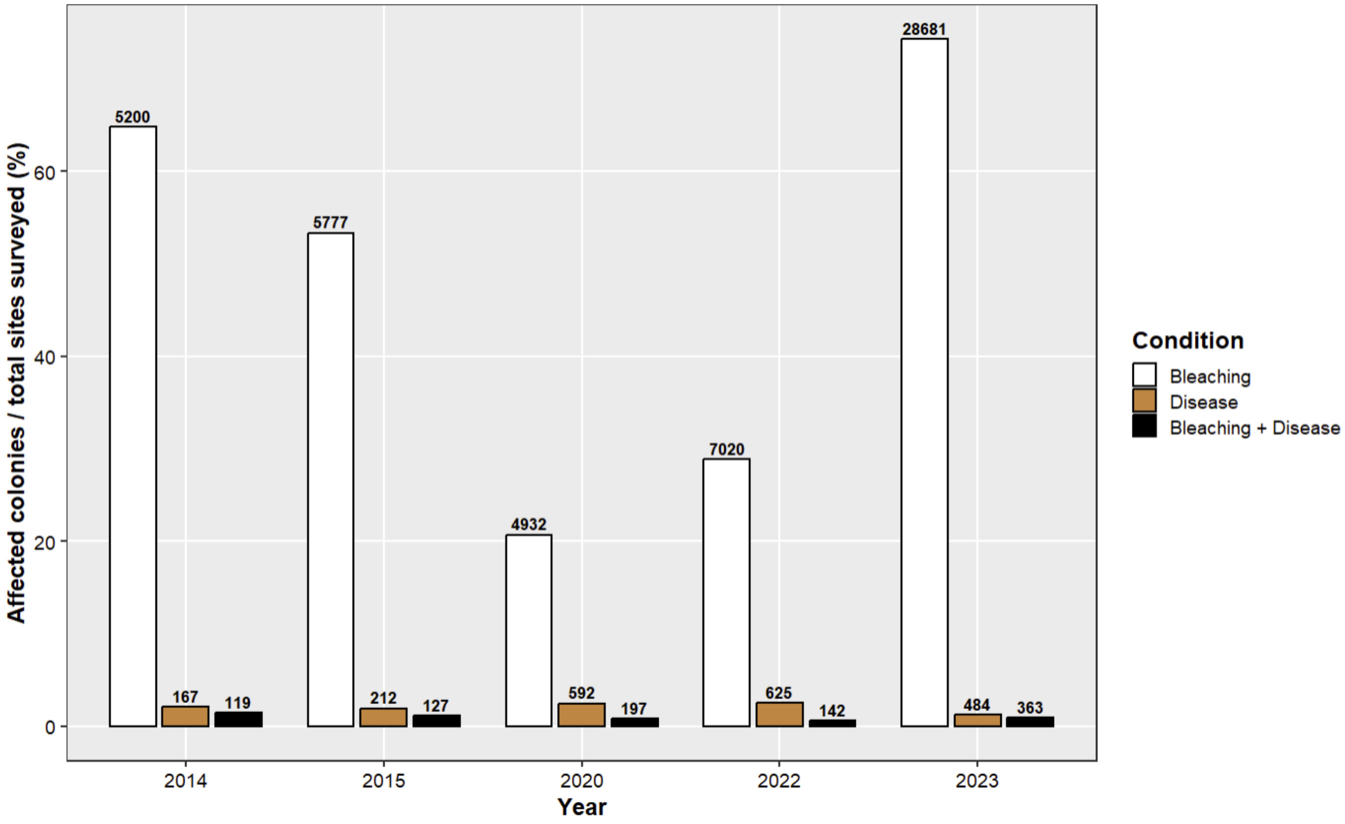

**Table S1. Summary of survey effort, bleaching severity, and heat stress across the FRT (2005–2023)**
Summary statistics of the DRM–FRRP dataset across all survey years, including number of sites, species richness, colony counts, bleaching severity metrics ($B_{site}$), and heat stress ($DHW_{site}$). DHW values represent the maximum heat exposure within the 14-day pre-survey window for each site. This table provides the full descriptive context for the dataset, including variation in sampling effort and thermal exposure, and supports selection of focal MHW years (2014, 2015, 2020, 2022 and 2023) used in subsequent analyses. The complete, unaggregated dataset comprising 216,080 colony-level observations has been deposited in Zenodo (https://zenodo.org/records/21464415?token=eyJhbGciOiJIUzUxMiJ9.eyJpZCI6IjJiMzRjMzNmLTI1ZmEtNDkxMC1hZGUyLWFiZDg0ZDE1NmJlYyIsImRhdGEiOnt9LCJyYW5kb20iOiI3ZjI5MDI2ODkzZjMwZWFjMmM1MWM3NzQyM2Q0MzJmYSJ9.PiNxZ-gpU_4T6tx-AqqUeszN-hipwA4bODO3MpLMkeLm0QTIdAX9QHovM7djxuOW0dehMMvdflnui-BxEbKt6w). The source code used for data analysis is hosted on the same Zenodo link. All data and code are currently restricted for peer review and will be made fully publicly accessible upon final peer-reviewed publication.

| Survey years | Data type | Total number of sites | Total number of species | Total number of colonies | Number species per site (Average ± SEM) | Number colonies per species - site (Average ±SEM ) | Bleaching severity per site (Bsite), sBRIk (Average ± SEM) | Number Severe Sites (Bsite>=0.3) Fig. 2 | % Recent mortality, RM, per colony-site (Average ± SEM) | colony density per site (Average ± SEM) | Numberof 5 x 5 km grid cells with DHW (NOAA CRW) | Degree Heating Weeks (DHW, °C-weeks) (Average ± SEM) | Severe Grids (Bgrid>= 0.3) | Slope of linear correlation (Bsite vs DHW), Fig. S1 | R2 linear correlation (Bsite vs DHW), Fig. S1 | Significance of linear correlation (Bsite vs DHW), p-value, Fig. S1 | criterium 2; 30% sites DHW >=4 AND 50% DHW> 4 | criterium 1; years with > 150 surveyed sites | Year included in this study |
|---|---|---|---|---|---|---|---|---|---|---|---|---|---|---|---|---|---|---|---|
| 2005 | entire FRT | 95 | 39 | 3451 | 8.29 ± 0.42 | 4.38 ± 0.22 | 0.17 ± 0.01 | 9 | 0.60 ± 0.15 | 36.33 ± 3.42 | 60 | 3.93 ± 0.17 | 9 | -0.339 | 0.003 | 0.631 | Pass | Fail | Excluded |
| 2006 | entire FRT | 121 | 43 | 4705 | 8.21 ± 0.36 | 4.74 ± 0.22 | 0.10 ± 0.01 | 5 | 0.66 ± 0.11 | 38.88 ± 3.01 | 62 | 0.02 ± 0.01 | 3 | -35.081 | 0.037 | 0.043 | Fail | Fail | Excluded |
| 2007 | entire FRT | 149 | 47 | 6031 | 8.17 ± 0.31 | 4.96 ± 0.24 | 0.15 ± 0.01 | 15 | 0.75 ± 0.13 | 40.48 ± 3.14 | 81 | 3.10 ± 0.30 | 13 | -0.79 | 0.035 | 0.026 | Pass | Fail | Excluded |
| 2008 | entire FRT | 204 | 43 | 10118 | 8.19 ± 0.28 | 6.06 ± 0.26 | 0.09 ± 0.01 | 7 | 0.72 ± 0.16 | 49.60 ± 3.65 | 85 | 0.16 ± 0.03 | 7 | -6.84 | 0.034 | 0.011 | Fail | Pass | Excluded |
| 2009 | entire FRT | 257 | 45 | 12622 | 8.82 ± 0.25 | 5.57 ± 0.19 | 0.13 ± 0.01 | 14 | 0.51 ± 0.08 | 49.11 ± 2.81 | 100 | 0.96 ± 0.12 | 13 | -1.982 | 0.04 | 0.002 | Fail | Pass | Excluded |
| 2010 | entire FRT | 200 | 41 | 7975 | 8.46 ± 0.28 | 4.71 ± 0.18 | 0.10 ± 0.01 | 7 | 0.51 ± 0.07 | 39.88 ± 2.59 | 83 | 1.81 ± 0.15 | 7 | -0.92 | 0.022 | 0.038 | Fail | Pass | Excluded |
| 2011 | entire FRT | 237 | 42 | 12481 | 9.39 ± 0.26 | 5.61 ± 0.20 | 0.14 ± 0.01 | 13 | 0.65 ± 0.12 | 52.66 ± 3.03 | 89 | 2.22 ± 0.20 | 11 | 1.466 | 0.076 | 0.000 | Fail | Pass | Excluded |
| 2012 | entire FRT | 233 | 52 | 10398 | 8.97 ± 0.25 | 4.97 ± 0.16 | 0.06 ± 0.01 | 4 | 0.73 ± 0.15 | 44.63 ± 2.31 | 85 | 0.25 ± 0.04 | 4 | -0.278 | 0 | 0.855 | Fail | Pass | Excluded |
| 2013 | entire FRT | 100 | 45 | 4241 | 8.39 ± 0.35 | 5.05 ± 0.32 | 0.09 ± 0.01 | 7 | 0.65 ± 0.25 | 42.41 ± 3.95 | 57 | 0.02 ± 0.01 | 7 | -4.37 | 0.001 | 0.792 | Fail | Fail | Excluded |
| 2014 | entire FRT | 174 | 47 | 7850 | 8.65 ± 0.30 | 5.22 ± 0.23 | 0.33 ± 0.01 | 86 | 1.16 ± 0.19 | 45.11 ± 3.05 | 80 | 3.64 ± 0.21 | 52 | 2.32 | 0.065 | 0.001 | Pass | Pass | Included |
| 2015 | entire FRT | 249 | 44 | 10613 | 7.84 ± 0.24 | 5.44 ± 0.22 | 0.29 ± 0.01 | 106 | 1.32 ± 0.18 | 42.62 ± 2.88 | 91 | 3.57 ± 0.24 | 59 | 1.827 | 0.095 | 0.000 | Pass | Pass | Included |
| 2016 | entire FRT | 162 | 46 | 7651 | 8.06 ± 0.33 | 5.86 ± 0.28 | 0.11 ± 0.01 | 7 | 0.75 ± 0.17 | 47.23 ± 3.59 | 84 | 1.33 ± 0.16 | 7 | -2.66 | 0.182 | 0.000 | Fail | Pass | Excluded |
| 2017 | entire FRT | 74 | 35 | 3947 | 8.77 ± 0.51 | 6.08 ± 0.43 | 0.12 ± 0.02 | 8 | 1.08 ± 0.25 | 53.34 ± 5.88 | 36 | 0.78 ± 0.14 | 8 | -4.081 | 0.043 | 0.091 | Fail | Fail | Excluded |
| 2018 | entire FRT | 195 | 47 | 8837 | 7.79 ± 0.26 | 5.81 ± 0.26 | 0.08 ± 0.00 | 2 | 0.07 ± 0.03 | 45.32 ± 3.15 | 92 | 1.23 ± 0.16 | 2 | -0.848 | 0.029 | 0.019 | Fail | Pass | Excluded |
| 2019 | entire FRT | 284 | 50 | 14242 | 7.94 ± 0.24 | 6.31 ± 0.24 | 0.07 ± 0.00 | 0 | 0.00 ± 0.00 | 50.15 ± 2.91 | 106 | 2.35 ± 0.17 | 0 | -0.348 | 0.009 | 0.133 | Fail | Pass | Excluded |
| 2020 | entire FRT | 388 | 45 | 19589 | 7.47 ± 0.19 | 6.75 ± 0.21 | 0.06 ± 0.00 | 0 | 0.00 ± 0.00 | 50.49 ± 2.46 | 118 | 2.77 ± 0.22 | 0 | -0.339 | 0.019 | 0.010 | Pass | Pass | Included |
| 2021 | entire FRT | 374 | 48 | 17995 | 7.59 ± 0.16 | 6.34 ± 0.21 | 0.05 ± 0.00 | 2 | 0.00 ± 0.00 | 48.11 ± 2.30 | 119 | 1.61 ± 0.13 | 2 | -0.523 | 0.02 | 0.007 | Fail | Pass | Exclude |
| 2022 | entire FRT | 444 | 47 | 19943 | 6.98 ± 0.15 | 6.44 ± 0.20 | 0.12 ± 0.00 | 25 | 0.00 ± 0.00 | 44.92 ± 2.04 | 136 | 3.91 ± 0.19 | 20 | 0.195 | 0.002 | 0.385 | Pass | Pass | Included |
| 2023 | entire FRT | 606 | 48 | 33391 | 7.80 ± 0.13 | 7.07 ± 0.18 | 0.45 ± 0.01 | 409 | 1.55 ± 0.16 | 55.10 ± 1.97 | 148 | 13.74 ± 0.31 | 107 | 2.932 | 0.17 | 0.000 | Pass | Pass | Included |
| 2014 | Repeatedly surveyed grids | 107 | 44 | 4965 | 8.626 ± 0.39 | 5.38 ± 0.30 | 0.35 ± 0.02 | 54 | 1.07 ± 0.23 | 46.40 ± 4.31 | 48 | 3.58 ± 0.19 | 26 | – | – | – | – | – | – |
| 2015 | Repeatedly surveyed grids | 153 | 40 | 5937 | 7.621 ± 0.29 | 5.09 ± 0.24 | 0.28 ± 0.01 | 64 | 1.26 ± 0.22 | 38.80 ± 3.16 | 48 | 3.05 ± 0.20 | 22 | – | – | – | – | – | – |
| 2020 | Repeatedly surveyed grids | 190 | 40 | 7597 | 6.547 ± 0.25 | 6.11 ± 0.30 | 0.06 ± 0.01 | 0 | 0.00 ± 0.00 | 39.98 ± 3.10 | 48 | 2.07 ± 0.18 | 0 | – | – | – | – | – | – |
| 2022 | Repeatedly surveyed grids | 225 | 42 | 8768 | 6.258 ± 0.22 | 6.23 ± 0.29 | 0.13 ± 0.01 | 15 | 0.00 ± 0.00 | 38.97 ± 2.85 | 48 | 2.99 ± 0.15 | 1 | – | – | – | – | – | – |
| 2023 | Repeatedly surveyed grids | 253 | 41 | 13316 | 7.395 ± 0.21 | 7.12 ± 0.29 | 0.39 ± 0.02 | 138 | 1.25 ± 0.20 | 52.63 ± 3.09 | 48 | 12.21 ± 0.23 | 28 | – | – | – | – | – | – |
| 2014 | repeatedly surveyed grids (Pair 2014-2015 ) | 134 | 44 | 5857 | 8.47 ± 0.34 | 5.16 ± 0.25 | 0.33 ± 0.01 | 64 | 1.00 ± 0.70 | 43.71 ± 3.56 | 63 | 3.55 ± 0.24 | 31 | – | – | – | – | – | – |
| 2015 | repeatedly surveyed grids (Pair 2014-2015 ) | 180 | 42 | 6838 | 7.63 ± 0.27 | 4.98 ± 0.21 | 0.28 ± 0.01 | 79 | 1.17 ± 0.66 | 37.99 ± 2.76 | 63 | 3.45 ± 0.28 | 29 | – | – | – | – | – | – |
| 2014 | repeatedly surveyed grids (Pair 2014-2020) | 155 | 47 | 7133 | 8.66 ± 0.32 | 5.31 ± 0.25 | 0.34 ± 0.01 | 77 | 1.21 ± 0.73 | 46.02 ± 3.35 | 68 | 3.60 ± 0.23 | 35 | – | – | – | – | – | – |
| 2020 | repeatedly surveyed grids (Pair 2014-2020) | 251 | 43 | 10978 | 7.06 ± 0.23 | 6.20 ± 0.25 | 0.06 ± 0.00 | 0 | 0.00 ± 0.42 | 43.74 ± 2.78 | 68 | 2.66 ± 0.30 | 0 | – | – | – | – | – | – |
| 2014 | repeatedly surveyed grids (Pair 2014-2022) | 160 | 47 | 7268 | 8.62 ± 0.31 | 5.27 ± 0.24 | 0.34 ± 0.01 | 82 | 1.21 ± 0.72 | 45.43 ± 3.27 | 71 | 3.53 ± 0.22 | 37 | – | – | – | – | – | – |
| 2022 | repeatedly surveyed grids (Pair 2014-2022) | 290 | 45 | 12029 | 6.51 ± 0.19 | 6.37 ± 0.25 | 0.13 ± 0.01 | 21 | 0.00 ± 0.46 | 41.48 ± 2.48 | 71 | 3.60 ± 0.25 | 1 | – | – | – | – | – | – |
| 2014 | repeatedly surveyed grids (Pair 2014-2023) | 165 | 46 | 7459 | 8.65 ± 0.30 | 5.22 ± 0.24 | 0.34 ± 0.01 | 83 | 1.21 ± 0.70 | 45.21 ± 3.19 | 74 | 3.61 ± 0.22 | 38 | – | – | – | – | – | – |
| 2023 | repeatedly surveyed grids (Pair 2014-2023) | 367 | 42 | 19353 | 7.63 ± 0.16 | 6.91 ± 0.23 | 0.44 ± 0.01 | 241 | 1.40 ± 0.47 | 52.73 ± 2.53 | 74 | 12.79 ± 0.38 | 51 | – | – | – | – | – | – |

**Table S2. Binomial GLM summaries of severe-bleaching reaction norms**

Model outputs for binomial GLMs relating the probability of severe bleaching ($B_{site} \geq 0.30$) to heat stress ($DHW_{site}$) for each focal MHW year (2014, 2015, 2020, 2022 and 2023, Eq.1). Models were fitted at both assemblage (site-level), grid-level, and species-specific scales. Reported statistics include coefficients, standard errors, and significance of the thermal sensitivity parameter (β1, Wald z-test). These models quantify year-specific reaction norms and demonstrate shifts in bleaching sensitivity consistent with ecological memory effects.

| | | | | | **Binomial GLM (logit link), Eq. 1** | | | | | | | |
|---|---|---|---|---|---|---|---|---|---|---|---|---|
| MHW | species | Number sites | Number grid | Number colonies | β0 intercept estimate | p- value (β0) | β1 thermal sensitivity slope estimate | β1 SE | β1 wald-z-score | p-value (β1) | β1 Significance | Supporting Figures |
| 2014 | – | 174 | – | – | -0.834 | 3.8E-17 | 0.258 | 0.094 | 2.740 | 1.8E-27 | ** | Figure 3a |
| 2015 | – | 249 | – | – | -0.902 | 2.7E-20 | 0.203 | 0.056 | 3.604 | 7.0E-27 | *** | Figure 3a |
| 2020 | – | 388 | – | – | -26.566 | 2.6E-43 | 0.000 | 8308.311 | 0.000 | 6.2E-02 | NS | Figure 3a |
| 2022 | – | 444 | – | – | -2.578 | 5.4E-65 | -0.036 | 0.094 | -0.387 | 8.9E-01 | NS | Figure 3a |
| 2023 | – | 606 | – | – | -3.882 | 6.3E-60 | 0.368 | 0.041 | 8.977 | 1.5E-115 | *** | Figure 3a |
| 2014 | – | 107 | – | – | -1.019 | 1.8E-02 | 0.312 | 0.107 | 2.925 | 6.2E-29 | ** | Figure S2a |
| 2015 | – | 153 | – | – | -1.261 | 1.7E-05 | 0.325 | 0.073 | 4.424 | 1.1E-41 | *** | Figure S2a |
| 2020 | – | 190 | – | – | -26.566 | 1.0E+00 | 0.000 | 11003.930 | 0.000 | 2.2E-01 | NS | Figure S2a |
| 2022 | – | 225 | – | – | -2.101 | 2.7E-06 | -0.138 | 0.134 | -1.024 | 6.5E-02 | NS | Figure S2a |
| 2023 | – | 253 | – | – | -8.072 | 1.4E-11 | 0.689 | 0.100 | 6.901 | 9.1E-67 | *** | Figure S2a |
| 2014 | – | – | 80 | – | -0.268 | 6.0E-01 | 0.158 | 0.127 | 1.246 | 2.1E-01 | NS | Figure S3 |
| 2015 | – | – | 91 | – | -0.948 | 2.5E-02 | 0.191 | 0.097 | 1.970 | 4.9E-02 | * | Figure S3 |
| 2022 | – | – | 136 | – | 0.000 | 1.0E+00 | -85.244 | 12243.470 | -0.007 | 9.9E-01 | NS | Figure S3 |
| 2023 | – | – | 148 | – | -3.291 | 8.2E-04 | 0.314 | 0.074 | 4.232 | 2.3E-05 | *** | Figure S3 |
| 2014 | AAGA | – | – | 459 | 1.799 | 4.0E-04 | 0.050 | 0.124 | 0.399 | 6.9E-01 | NS | Figure 4 |
| 2015 | AAGA | – | – | 418 | 1.380 | 8.9E-04 | 0.132 | 0.092 | 1.433 | 1.5E-01 | NS | Figure 4 |
| 2020 | AAGA | – | – | 1111 | -2.127 | 3.7E-32 | 0.136 | 0.048 | 2.851 | 4.4E-03 | ** | Figure 4 |
| 2022 | AAGA | – | – | 1170 | -0.604 | 3.7E-03 | 0.244 | 0.043 | 5.701 | 1.2E-08 | *** | Figure 4 |
| 2023 | AAGA | – | – | 1926 | 0.577 | 1.2E-01 | 0.145 | 0.027 | 5.347 | 9.0E-08 | *** | Figure 4 |
| 2014 | MCAV | – | – | 456 | -2.150 | 3.8E-11 | 0.193 | 0.071 | 2.703 | 6.9E-03 | ** | Figure 4 |
| 2015 | MCAV | – | – | 709 | -1.282 | 2.6E-14 | -0.077 | 0.039 | -1.969 | 4.9E-02 | * | Figure 4 |
| 2020 | MCAV | – | – | 1442 | -2.496 | 1.5E-61 | -0.081 | 0.049 | -1.666 | 9.6E-02 | NS | Figure 4 |
| 2022 | MCAV | – | – | 1123 | -2.836 | 1.1E-22 | -0.061 | 0.062 | -0.970 | 3.3E-01 | NS | Figure 4 |
| 2023 | MCAV | – | – | 1906 | -2.433 | 4.7E-30 | 0.205 | 0.015 | 13.803 | 2.4E-43 | *** | Figure 4 |
| 2014 | OFAV | – | – | 93 | -1.107 | 9.1E-02 | 0.207 | 0.140 | 1.483 | 1.4E-01 | NS | Figure 4 |
| 2015 | OFAV | – | – | 181 | -1.313 | 3.9E-03 | 0.174 | 0.080 | 2.191 | 2.8E-02 | * | Figure 4 |
| 2020 | OFAV | – | – | 304 | -1.721 | 5.1E-09 | -0.131 | 0.085 | -1.535 | 1.2E-01 | NS | Figure 4 |
| 2022 | OFAV | – | – | 303 | -1.811 | 1.5E-05 | -0.096 | 0.086 | -1.126 | 2.6E-01 | NS | Figure 4 |
| 2023 | OFAV | – | – | 606 | 0.253 | 5.2E-01 | 0.065 | 0.025 | 2.594 | 9.5E-03 | * | Figure 4 |
| 2014 | PAST | – | – | 1503 | -2.250 | 1.5E-29 | 0.460 | 0.042 | 11.050 | 2.2E-28 | *** | Figure 4 |
| 2015 | PAST | – | – | 1820 | -1.009 | 2.7E-18 | 0.127 | 0.023 | 5.438 | 5.4E-08 | *** | Figure 4 |
| 2020 | PAST | – | – | 3547 | -3.642 | 1.2E-88 | -0.074 | 0.053 | -1.390 | 1.6E-01 | NS | Figure 4 |
| 2022 | PAST | – | – | 4467 | -3.193 | 9.3E-64 | 0.003 | 0.039 | 0.064 | 9.5E-01 | NS | Figure 4 |
| 2023 | PAST | – | – | 7415 | -1.171 | 1.1E-22 | 0.163 | 0.009 | 19.080 | 3.7E-81 | *** | Figure 4 |
| 2014 | PPOR | – | – | 538 | 0.376 | 2.0E-01 | 0.071 | 0.067 | 1.066 | 2.9E-01 | NS | Figure 4 |
| 2015 | PPOR | – | – | 686 | -0.457 | 8.0E-03 | 0.228 | 0.041 | 5.567 | 2.6E-08 | *** | Figure 4 |
| 2020 | PPOR | – | – | 714 | -3.138 | 7.8E-24 | -0.008 | 0.082 | -0.097 | 9.2E-01 | NS | Figure 4 |
| 2022 | PPOR | – | – | 1082 | -1.282 | 3.7E-12 | 0.038 | 0.039 | 0.980 | 3.3E-01 | NS | Figure 4 |
| 2023 | PPOR | – | – | 1532 | -1.042 | 1.1E-04 | 0.206 | 0.021 | 9.854 | 6.6E-23 | *** | Figure 4 |
| 2014 | SINT | – | – | 935 | -0.929 | 2.0E-05 | 0.028 | 0.043 | 0.655 | 5.1E-01 | NS | Figure 4 |
| 2015 | SINT | – | – | 1358 | -0.287 | 2.7E-02 | -0.165 | 0.026 | -6.371 | 1.9E-10 | *** | Figure 4 |
| 2020 | SINT | – | – | 3131 | -2.480 | 6.0E-76 | -0.171 | 0.038 | -4.510 | 6.5E-06 | *** | Figure 4 |
| 2022 | SINT | – | – | 2686 | -1.630 | 1.8E-25 | -0.121 | 0.031 | -3.863 | 1.1E-04 | *** | Figure 4 |
| 2023 | SINT | – | – | 4126 | -0.585 | 1.2E-05 | 0.028 | 0.009 | 3.192 | 1.4E-03 | ** | Figure 4 |
| 2014 | SRAD | – | – | 267 | -0.812 | 1.9E-02 | 0.077 | 0.078 | 0.989 | 3.2E-01 | NS | Figure 4 |
| 2015 | SRAD | – | – | 414 | -0.813 | 7.2E-04 | -0.051 | 0.045 | -1.118 | 2.6E-01 | NS | Figure 4 |
| 2020 | SRAD | – | – | 325 | -1.163 | 4.0E-09 | -0.129 | 0.060 | -2.146 | 3.2E-02 | * | Figure 4 |
| 2022 | SRAD | – | – | 373 | -0.702 | 2.5E-03 | -0.140 | 0.057 | -2.442 | 1.5E-02 | * | Figure 4 |
| 2023 | SRAD | – | – | 688 | -1.795 | 1.3E-12 | 0.096 | 0.018 | 5.385 | 7.3E-08 | *** | Figure 4 |
| 2014 | SSID | – | – | 2150 | -0.531 | 2.6E-04 | 0.028 | 0.029 | 0.950 | 3.4E-01 | NS | Figure 4 |
| 2015 | SSID | – | – | 3087 | -0.826 | 5.6E-13 | -0.156 | 0.023 | -6.695 | 2.2E-11 | *** | Figure 4 |
| 2020 | SSID | – | – | 6953 | -1.499 | 5.3E-109 | -0.240 | 0.020 | -12.259 | 1.5E-34 | *** | Figure 4 |
| 2022 | SSID | – | – | 6582 | -0.769 | 5.3E-20 | -0.253 | 0.018 | -14.179 | 1.2E-45 | *** | Figure 4 |
| 2023 | SSID | – | – | 11579 | -0.513 | 4.0E-12 | 0.035 | 0.005 | 7.223 | 5.1E-13 | *** | Figure 4 |
| 2014 | CNAT | – | – | 138 | -4.605 | 6.89E-06 | 1.018 | 0.202 | 5.053 | 4.36E-07 | *** | Figure 4 |
| 2015 | CNAT | – | – | 205 | -0.476 | 0.442 | -0.089 | 0.102 | -0.875 | 0.382 | NS | Figure 4 |
| 2020 | CNAT | – | – | 131 | -2.252 | 2.75E-04 | -0.640 | 0.483 | -1.325 | 0.185 | NS | Figure 4 |
| 2022 | CNAT | – | – | 115 | -2.725 | 0.006 | 0.070 | 0.171 | 0.411 | 0.681 | NS | Figure 4 |
| 2023 | CNAT | – | – | 194 | -0.351 | 0.523 | 0.049 | 0.035 | 1.408 | 0.159 | NS | Figure 4 |
| 2014 | DSTO | – | – | 271 | -1.796 | 5.40E-04 | 0.003 | 0.114 | 0.025 | 0.980 | NS | Figure 4 |
| 2015 | DSTO | – | – | 357 | -2.150 | 2.81E-09 | 0.066 | 0.073 | 0.905 | 0.366 | NS | Figure 4 |
| 2020 | DSTO | – | – | 192 | -3.526 | 2.63E-08 | 0.096 | 0.177 | 0.539 | 0.590 | NS | Figure 4 |
| 2022 | DSTO | – | – | 217 | -2.905 | 1.61E-05 | -0.006 | 0.147 | -0.042 | 0.967 | NS | Figure 4 |
| 2023 | DSTO | – | – | 348 | -3.352 | 1.72E-08 | 0.181 | 0.039 | 4.650 | 3.33E-06 | *** | Figure 4 |
| 2014 | PSTR | – | – | 131 | -1.541 | 0.002 | 0.186 | 0.112 | 1.667 | 0.096 | NS | Figure 4 |
| 2015 | PSTR | – | – | 138 | -0.598 | 0.208 | 0.019 | 0.088 | 0.215 | 0.830 | NS | Figure 4 |
| 2020 | PSTR | – | – | 147 | -3.058 | 2.22E-10 | 0.230 | 0.126 | 1.820 | 0.069 | NS | Figure 4 |
| 2022 | PSTR | – | – | 144 | -3.288 | 3.51E-04 | 0.123 | 0.173 | 0.713 | 0.476 | NS | Figure 4 |
| 2023 | PSTR | – | – | 220 | -0.228 | 0.731 | 0.054 | 0.047 | 1.146 | 0.252 | NS | Figure 4 |

**Table S3. Statistical comparisons of DHW distributions among MHW years**

Non-parametric tests comparing distributions of heat stress (DHW) among focal MHW years (2014, 2015, 2020, 2022 and 2023). For full FRT analyses, Kruskal–Wallis tests followed by pairwise post-hoc Wilcoxon rank-sum tests with Bonferroni correction were applied. For repeatedly surveyed grids, Friedman test followed by post-hoc paired Wilcoxon signed-rank tests with Bonferroni correction were applied. These analyses establish whether bleaching responses occurred under comparable or distinct thermal regimes, supporting interpretation of ecological memory effects. The complete set of repeatedly surveyed grids, with 76,915 colony-level observations for focal years 2014, 2015, 202, 2022 and 2023 has been deposited in Zenodo (https://zenodo.org/records/21464415?token=eyJhbGciOiJIUzUxMiJ9.eyJpZCI6IjJiMzRjMzNmLTI1ZmEtNDkxMC1hZGUyLWFiZDg0ZDE1NmJlYyIsImRhdGEiOnt9LCJyYW5kb20iOiI3ZjI5MDI2ODkzZjMwZWFjMmM1MWM3NzQyM2Q0MzJmYSJ9.PiNxZ-gpU_4T6tx-AqqUeszN-hipwA4bODO3MpLMkeLm0QTIdAX9QHovM7djxuOW0dehMMvdflnui-BxEbKt6w). The source code used for data analysis is hosted on the same Zenodo link. All data and code are currently restricted for peer review and will be made fully publicly accessible upon final peer-reviewed publication.

| Dataset | Year / Comparison | Number grids | Number sites | Mean DHWgrid ± SEM | Test | Statistic | p-value | Significance | Supporting Figure |
|---|---|---|---|---|---|---|---|---|---|
| Entire FRT | 2014 | 80 | 174 | 3.65 ± 0.209 | – | – | – | – | Figure 3 |
| Entire FRT | 2015 | 91 | 249 | 3.55 ± 0.241 | – | – | – | – | Figure 3 |
| Entire FRT | 2020 | 118 | 388 | 2.76 ± 0.218 | – | – | – | – | Figure 3 |
| Entire FRT | 2022 | 136 | 444 | 3.93 ± 0.188 | – | – | – | – | Figure 3 |
| Entire FRT | 2023 | 148 | 606 | 13.8 ± 0.305 | – | – | – | – | Figure 3 |
| Repeatedly surveyed Grids | 2014 | 48 | 107 | 3.56 ± 0.293 | – | – | – | – | Figure S2 |
| Repeatedly surveyed Grids | 2015 | 48 | 153 | 3.53 ± 0.352 | – | – | – | – | Figure S2 |
| Repeatedly surveyed Grids | 2020 | 48 | 190 | 2.72 ± 0.374 | – | – | – | – | Figure S2 |
| Repeatedly surveyed Grids | 2022 | 48 | 225 | 3.50 ± 0.322 | – | – | – | – | Figure S2 |
| Repeatedly surveyed Grids | 2023 | 48 | 253 | 12.9 ± 0.475 | – | – | – | – | Figure S2 |
| Entire FRT | All years | – | – | – | Kruskal–Wallis | 308 | 1.90E-65 | *** | |
| Entire FRT | 2014 vs 2015 | – | – | – | Wilcoxon rank-sum (Bonferroni) | 3810 | 1 | ns | Figure 3b |
| Entire FRT | 2014 vs 2020 | – | – | – | Wilcoxon rank-sum (Bonferroni) | 5966 | 0.016 | * | Figure 3b |
| Entire FRT | 2014 vs 2022 | – | – | – | Wilcoxon rank-sum (Bonferroni) | 5094 | 1 | ns | Figure 3b |
| Entire FRT | 2014 vs 2023 | – | – | – | Wilcoxon rank-sum (Bonferroni) | 306 | 3.53E-31 | *** | Figure 3b |
| Entire FRT | 2015 vs 2020 | – | – | – | Wilcoxon rank-sum (Bonferroni) | 6514 | 0.083 | ns | |
| Entire FRT | 2015 vs 2022 | – | – | – | Wilcoxon rank-sum (Bonferroni) | 5606 | 1 | ns | |
| Entire FRT | 2015 vs 2023 | – | – | – | Wilcoxon rank-sum (Bonferroni) | 333 | 6.06E-34 | *** | |
| Entire FRT | 2020 vs 2022 | – | – | – | Wilcoxon rank-sum (Bonferroni) | 5642 | 4.52E-04 | *** | |
| Entire FRT | 2020 vs 2023 | – | – | – | Wilcoxon rank-sum (Bonferroni) | 357 | 3.58E-40 | *** | |
| Entire FRT | 2022 vs 2023 | – | – | – | Wilcoxon rank-sum (Bonferroni) | 550 | 4.50E-42 | *** | |
| Repeatedly surveyed Grids | All years | – | – | – | Friedman | 115 | 5.43E-24 | *** | |
| Repeatedly surveyed Grids | 2014 vs 2015 | – | – | – | Wilcoxon signed-rank (Bonferroni) | 753 | 0.916 | ns | Figure S2b |
| Repeatedly surveyed Grids | 2014 vs 2020 | – | – | – | Wilcoxon signed-rank (Bonferroni) | 1013 | 1.34E-04 | *** | Figure S2b |
| Repeatedly surveyed Grids | 2014 vs 2022 | – | – | – | Wilcoxon signed-rank (Bonferroni) | 573 | 1 | ns | Figure S2b |
| Repeatedly surveyed Grids | 2014 vs 2023 | – | – | – | Wilcoxon signed-rank (Bonferroni) | 0 | 7.11E-14 | *** | Figure S2b |
| Repeatedly surveyed Grids | 2015 vs 2020 | – | – | – | Wilcoxon signed-rank (Bonferroni) | 856 | 0.02 | * | |
| Repeatedly surveyed Grids | 2015 vs 2022 | – | – | – | Wilcoxon signed-rank (Bonferroni) | 566 | 1 | ns | |
| Repeatedly surveyed Grids | 2015 vs 2023 | – | – | – | Wilcoxon signed-rank (Bonferroni) | 0 | 7.11E-14 | *** | |
| Repeatedly surveyed Grids | 2020 vs 2022 | – | – | – | Wilcoxon signed-rank (Bonferroni) | 308 | 0.042 | * | |
| Repeatedly surveyed Grids | 2020 vs 2023 | – | – | – | Wilcoxon signed-rank (Bonferroni) | 0 | 7.11E-14 | *** | |
| Repeatedly surveyed Grids | 2022 vs 2023 | – | – | – | Wilcoxon signed-rank (Bonferroni) | 0 | 7.11E-14 | *** | |

**Table S4. Grid-level trajectory classifications and underlying metrics**

Complete grid-level dataset used to classify matched-event trajectories across pairwise comparisons (2014–2015, 2014–2020, 2014–2022, and 2014–2023). For each grid (5 × 5 km), the table reports: (i) Bleaching severity ($B_{grid}$) and adult colony density ($CD_{adult,grid}$); (ii) Changes relative to 2014 ($\Delta B_{grid}$ and $\Delta CD_{adult,grid}$; Eq. 3); (iii) Heat stress metrics ($DHW_{grid}$ values); (iv) Thermal classification [thermally typical (within regression envelope (±1.5 SD of residuals) vs ND_thermal (within regression envelope (±1.5 SD of residuals)]; (v) Trajectory assignment: H2 (sensitization: $\Delta B_{grid} > 0$); H3 (acclimatization/persistence: $\Delta B_{grid} \leq 0$ and $\Delta CD_{adult,grid} \geq 0$); H4 (mortality-filtered tolerance: $\Delta B_{grid} \leq 0$ and $\Delta CD_{adult,grid} < 0$). All grid-level values were derived using two-step averaging (colony density was calculated at site-level first and then averaged across all sites mapped to a grid-year) with true zeros retained. This table is the core classification dataset underpinning all trajectory, stability, and species-decomposition analyses. NA indicates that no sites were surveyed for a given grid-year combination.

| Grid ID_number | Grid ID [lat,long] | Subregion | Avg Depth (m) | Number Sites 2014 | ST/Gen Adults (>=10cm) per site, 2014 (AVG ± SEM) | Weedy Adults (>=5cm) per site, 2014 (AVG ± SEM) | Number Sites 2015 | ST/Gen Adults (>=10cm) per site, 2015 (AVG ± SEM) | Weedy Adults (>=5cm) per site, 2015 (AVG ± SEM) | Number Sites 2020 | ST/Gen Adults (>=10cm), per site, 2020 (AVG ± SEM) | Weedy Adults (>=5cm), per site, 2020 (AVG ± SEM) | Number Sites 2022 | ST/Gen Adults (>=10cm), per site, 2022 (AVG ± SEM) | Weedy Adults (>=5cm), per site, 2022 (AVG ± SEM) | Number Sites 2023 | ST/Gen Adults (>=10cm), per site, 2023 (AVG ± SEM) | Weedy Adults (>=5cm), per site, 2023 (AVG ± SEM) | 2014 v 2015 State | 2014 v 2020 State | 2014 v 2022 State | 2014 v 2023 State |
|---|---|---|---|---|---|---|---|---|---|---|---|---|---|---|---|---|---|---|---|---|---|---|
| 9 | (24.425, -81.975) | Lower Keys | 33.51 | 1 | 8 | 20 | 2 | 4.5 ± 2.5 | 12 ± 10 | 4 | 15.75 ± 7.97 | 12.25 ± 1.7 | 1 | 11 | 37 | 3 | 10.67 ± 3.18 | 14.33 ± 5.21 | ND_thermal | H3 | H3 | H2 |
| 10 | (24.425, -81.925) | Lower Keys | 31.25 | 1 | 20 | 9 | 2 | 0.5 ± 0.5 | 10 ± 6 | NA | NA | NA | NA | NA | NA | 1 | 5 | 9 | ND_thermal | NA | NA | H2 |
| 22 | (24.475, -81.925) | Lower Keys | 30.22 | 1 | 57 | 17 | NA | NA | NA | 1 | 18 | 30 | 2 | 29.5 ± 15.5 | 14 ± 3 | 2 | 88 ± 54 | 24.5 ± 18.5 | NA | H4 | H4 | H3 |
| 23 | (24.475, -81.875) | Lower Keys | 22.57 | 1 | 8 | 9 | 4 | 19.75 ± 9.44 | 24.25 ± 5.25 | 3 | 35 ± 3.06 | 17 ± 7.09 | 5 | 25 ± 15.21 | 8.8 ± 2.87 | 9 | 45.11 ± 6.84 | 22.89 ± 4.14 | H3 | H3 | H3 | ND_thermal |
| 24 | (24.475, -81.825) | Lower Keys | 25.70 | 2 | 27 ± 7 | 12 ± 5 | 2 | 28.5 ± 27.5 | 18 ± 14 | 6 | 10.33 ± 3.42 | 4.33 ± 0.95 | 2 | 13 ± 0 | 2 ± 2 | 2 | 57.5 ± 13.5 | 8 ± 1 | H2 | H4 | H4 | ND_thermal |
| 25 | (24.475, -81.775) | Lower Keys | 32.17 | 1 | 4 | 23 | 2 | 5 ± 5 | 8 ± 2 | 4 | 15.5 ± 8.29 | 11 ± 5.03 | 2 | 62.5 ± 10.5 | 14 ± 7 | 1 | 36 | 14 | H4 | H4 | ND_thermal | ND_thermal |
| 26 | (24.475, -81.725) | Lower Keys | 26.55 | 4 | 15.5 ± 11.51 | 15.25 ± 3.35 | 3 | 13 ± 9.5 | 12 ± 2.08 | 6 | 13.67 ± 4.67 | 11 ± 4.61 | 3 | 23.67 ± 10.09 | 10.33 ± 4.81 | 5 | 35 ± 18.89 | 33.2 ± 13.18 | H4 | H4 | H3 | H3 |
| 39 | (24.525, -81.725) | Lower Keys | 21.75 | 1 | 34 | 44 | 2 | 25.5 ± 11.5 | 10 ± 7 | 6 | 53.33 ± 10.9 | 7.5 ± 2.26 | 4 | 61.25 ± 5.25 | 14.75 ± 5.07 | 4 | 60.75 ± 8.77 | 8.75 ± 1.55 | ND_thermal | H4 | H3 | H4 |
| 42 | (24.525, -81.575) | Lower Keys | 21.40 | 6 | 60.83 ± 19.67 | 17.5 ± 2.36 | 3 | 6.33 ± 2.85 | 10.67 ± 3.53 | 6 | 42 ± 17.07 | 27.33 ± 12.75 | 6 | 33.33 ± 8.47 | 25.83 ± 9.71 | 8 | 37.75 ± 10.52 | 15.75 ± 5.28 | H4 | H4 | ND_thermal | H2 |
| 43 | (24.525, -81.525) | Lower Keys | 29.08 | 4 | 20.5 ± 11.21 | 36.25 ± 9.07 | 4 | 16 ± 4.51 | 11.5 ± 4.48 | 2 | 36 ± 1 | 21.5 ± 7.5 | 2 | 17 ± 15 | 24 ± 8 | 1 | 36 | 29 | H4 | H3 | H4 | H2 |
| 44 | (24.525, -81.475) | Lower Keys | 31.46 | 6 | 21.5 ± 6.15 | 14.5 ± 2.84 | NA | NA | NA | 4 | 9.75 ± 5.23 | 3 ± 1.73 | 5 | 28.6 ± 6.18 | 12.2 ± 3.95 | 1 | 7 | 6 | NA | H4 | H3 | H2 |
| 45 | (24.525, -81.425) | Lower Keys | 26.63 | 2 | 17 ± 11 | 24.5 ± 16.5 | 4 | 28.75 ± 7.62 | 35.25 ± 6.98 | 1 | 9 | 4 | 4 | 9 ± 5.58 | 13.25 ± 4.7 | 2 | 9 ± 0 | 34 ± 28 | H2 | H4 | ND_thermal | H2 |
| 47 | (24.575, -82.975) | Tortugas--Dry Tortugas NP | 46.66 | 2 | 7.5 ± 6.5 | 2 ± 1 | NA | NA | NA | NA | NA | NA | 2 | 20.5 ± 6.5 | 12.5 ± 2.5 | 6 | 12.33 ± 1.76 | 16.67 ± 4.58 | NA | NA | H3 | H2 |
| 48 | (24.575, -82.925) | Tortugas--Dry Tortugas NP | 38.67 | 2 | 18 ± 6 | 16.5 ± 3.5 | NA | NA | NA | 2 | 10.5 ± 5.5 | 18.5 ± 12.5 | 7 | 7.86 ± 0.67 | 15.71 ± 2.08 | 8 | 14.75 ± 3.75 | 20.5 ± 2.95 | NA | ND_thermal | H4 | H2 |
| 53 | (24.575, -81.575) | Lower Keys | 16.19 | 1 | 40 | 26 | 3 | 14.67 ± 14.67 | 5.33 ± 1.2 | 2 | 62.5 ± 5.5 | 13.5 ± 5.5 | 4 | 44 ± 9.01 | 11 ± 3.37 | 5 | 61.6 ± 10.68 | 19.8 ± 5.51 | H4 | H3 | H4 | H2 |
| 55 | (24.575, -81.475) | Lower Keys | 25.64 | 1 | 86 | 17 | 1 | 50 | 20 | 2 | 39 ± 29 | 14.5 ± 14.5 | 2 | 44.5 ± 11.5 | 16 ± 3 | 1 | 73 | 30 | H4 | H4 | H4 | ND_thermal |
| 56 | (24.575, -81.425) | Lower Keys | 21.93 | 2 | 32.5 ± 6.5 | 13 ± 5 | 1 | 16 | 7 | 2 | 24.5 ± 14.5 | 2.5 ± 0.5 | 1 | 8 | 8 | 3 | 22.33 ± 2.4 | 7 ± 1.15 | H4 | H4 | H4 | H4 |
| 57 | (24.575, -81.375) | Lower Keys | 24.35 | 4 | 53 ± 26.64 | 30 ± 12.26 | 2 | 7 ± 1 | 19 ± 11 | 5 | 31.8 ± 17.09 | 10.6 ± 5.13 | 9 | 37.56 ± 7.26 | 11.33 ± 2.01 | 5 | 65.8 ± 30.62 | 20 ± 7.26 | H4 | H4 | H4 | H2 |
| 58 | (24.575, -81.325) | Lower Keys | 25.40 | 2 | 22 ± 13 | 16.5 ± 4.5 | 4 | 8.75 ± 5.78 | 15.25 ± 6.87 | 1 | 1 | 13 | 3 | 10.33 ± 3.18 | 4.67 ± 2.33 | 1 | 13 | 4 | H2 | H4 | ND_thermal | H2 |
| 59 | (24.575, -81.275) | Middle Keys | 28.24 | 1 | 4 | 0 | 3 | 11 ± 3.51 | 5.67 ± 1.45 | 1 | 2 | 1 | 1 | 12 | 4 | NA | NA | NA | H2 | H4 | H3 | NA |
| 60 | (24.575, -81.225) | Middle Keys | 40.73 | 2 | 8 ± 8 | 11 ± 6 | 3 | 17.67 ± 10.2 | 16 ± 8.02 | 2 | 12.5 ± 0.5 | 4.5 ± 0.5 | 2 | 11 ± 6 | 4 ± 3 | 3 | 7.67 ± 2.33 | 9.33 ± 6.01 | H2 | H4 | ND_thermal | H2 |
| 62 | (24.625, -82.975) | Tortugas--Dry Tortugas NP | 44.34 | 3 | 17.33 ± 6.57 | 6 ± 1.53 | NA | NA | NA | 3 | 16.33 ± 9.21 | 17.67 ± 6.17 | 1 | 6 | 8 | 7 | 10.57 ± 4.39 | 9.14 ± 2.86 | NA | H2 | H2 | H2 |
| 63 | (24.625, -82.925) | Tortugas--Dry Tortugas NP | 35.66 | 3 | 31 ± 2.08 | 25 ± 0.58 | 4 | 15.25 ± 4.59 | 9 ± 0.82 | 5 | 14.6 ± 7.45 | 19.6 ± 3.74 | 5 | 9 ± 7.26 | 19.2 ± 2.08 | 12 | 22.25 ± 5.27 | 19.25 ± 3.57 | ND_thermal | H4 | H4 | H2 |
| 64 | (24.625, -82.875) | Tortugas--Dry Tortugas NP | 33.57 | 4 | 17 ± 13.67 | 14.75 ± 2.06 | 1 | 23 | 10 | 5 | 43.2 ± 12 | 17.8 ± 4.52 | NA | NA | NA | 10 | 13.1 ± 3.49 | 16.9 ± 4.88 | ND_thermal | ND_thermal | NA | H2 |
| 65 | (24.625, -82.825) | Tortugas--Dry Tortugas NP | 30.26 | 5 | 12 ± 5.22 | 12.8 ± 3.25 | 2 | 16 ± 0 | 12.5 ± 8.5 | 9 | 13.11 ± 4.21 | 22.78 ± 4.19 | 5 | 13.6 ± 4.3 | 27 ± 5.75 | 6 | 22.33 ± 5.44 | 26.67 ± 4.72 | ND_thermal | ND_thermal | H3 | H2 |
| 68 | (24.625, -81.375) | Lower Keys | 16.33 | 2 | 1.5 ± 1.5 | 0.5 ± 0.5 | NA | NA | NA | 2 | 12.5 ± 12.5 | 11.5 ± 8.5 | 5 | 16.6 ± 3.59 | 6.8 ± 5.58 | 3 | 25.67 ± 14.71 | 8.67 ± 3.76 | NA | H3 | H3 | ND_thermal |
| 70 | (24.625, -81.225) | Middle Keys | 28.18 | 2 | 12.5 ± 11.5 | 11 ± 11 | 1 | 25 | 24 | 1 | 5 | 3 | 1 | 0 | 1 | 6 | 58 ± 10.21 | 19.17 ± 7.73 | ND_thermal | ND_thermal | H4 | H2 |
| 72 | (24.625, -81.125) | Middle Keys | 32.49 | 1 | 1 | 3 | 3 | 17 ± 4.16 | 21 ± 11.15 | 3 | 17.33 ± 3.28 | 21.33 ± 8.84 | 4 | 20.75 ± 9.52 | 11.5 ± 3.43 | 2 | 3 ± 0 | 40 ± 10 | ND_thermal | ND_thermal | H3 | ND_thermal |
| 76 | (24.675, -82.925) | Tortugas--Dry Tortugas NP | 38.39 | 1 | 7 | 1 | 1 | 46 | 13 | 3 | 30 ± 18.5 | 24.33 ± 2.91 | 5 | 11.4 ± 3.85 | 17.6 ± 4.04 | 9 | 16.33 ± 4.54 | 17.89 ± 3.18 | ND_thermal | H3 | H2 | H2 |
| 77 | (24.675, -82.875) | Tortugas--Dry Tortugas NP | 24.06 | 2 | 29.5 ± 7.5 | 36.5 ± 0.5 | 5 | 24.2 ± 9.49 | 15.2 ± 2.4 | 4 | 29.25 ± 5.28 | 24 ± 4.71 | 4 | 24.25 ± 5.17 | 20.5 ± 2.5 | 5 | 15.2 ± 5.2 | 29.8 ± 7.62 | ND_thermal | ND_thermal | H4 | H2 |
| 78 | (24.675, -82.825) | Tortugas--Dry Tortugas NP | 21.91 | 4 | 19.5 ± 5.11 | 10.5 ± 1.32 | NA | NA | NA | 3 | 37.33 ± 3.18 | 29.67 ± 9.17 | 3 | 7.33 ± 6.36 | 15 ± 7.57 | 6 | 12.83 ± 6.82 | 15.5 ± 5.8 | NA | ND_thermal | H4 | H2 |
| 81 | (24.675, -81.075) | Middle Keys | 18.51 | 2 | 26 ± 12 | 43.5 ± 36.5 | NA | NA | NA | 3 | 51 ± 34.24 | 28 ± 16.17 | 2 | 15.5 ± 10.5 | 36.5 ± 36.5 | 3 | 46 ± 22.55 | 40.67 ± 11.32 | NA | ND_thermal | H4 | H2 |
| 82 | (24.675, -81.025) | Middle Keys | 18.88 | 2 | 3 ± 0 | 0 | 5 | 42.6 ± 16.44 | 10.6 ± 6.43 | 3 | 9.67 ± 8.69 | 11.33 ± 5.81 | 2 | 34.5 ± 7.5 | 38 ± 18 | 6 | 62.5 ± 22.37 | 42.5 ± 10.9 | H2 | H3 | ND_thermal | H2 |
| 83 | (24.675, -80.975) | Middle Keys | 16.94 | 3 | 6.33 ± 2.03 | 4.67 ± 2.4 | 2 | 99 ± 51 | 42 ± 21 | 4 | 11.75 ± 6.99 | 10 ± 5.18 | 5 | 23 ± 8.08 | 43 ± 6.4 | 1 | 15 | 16 | H2 | H3 | ND_thermal | ND_thermal |
| 87 | (24.725, -82.875) | Tortugas--Dry Tortugas NP | 39.35 | 3 | 11.33 ± 3.67 | 8.33 ± 2.19 | NA | NA | NA | 7 | 17.86 ± 2.54 | 21.86 ± 6.95 | 4 | 5 ± 2.74 | 11 ± 3.49 | 6 | 9 ± 2.29 | 14.17 ± 2.71 | NA | H3 | H4 | H2 |
| 91 | (24.725, -80.925) | Middle Keys | 20.05 | 2 | 32 ± 22 | 7.5 ± 4.5 | 1 | 71 | 25 | 3 | 67.33 ± 11.39 | 44.33 ± 23.38 | 7 | 44.43 ± 9.34 | 40 ± 14.44 | 2 | 93.5 ± 21.5 | 55.5 ± 20.5 | H2 | ND_thermal | H3 | H2 |
| 107 | (24.925, -80.525) | Upper Keys | 25.57 | 1 | 5 | 7 | NA | NA | NA | 1 | 11 | 13 | 4 | 12.25 ± 3.86 | 28.25 ± 6.18 | 6 | 27.5 ± 16.33 | 15.83 ± 7 | NA | H3 | H3 | H3 |
| 110 | (24.975, -80.525) | Upper Keys | 16.80 | 1 | 1 | 1 | NA | NA | NA | 1 | 8 | 0 | 1 | 42 | 4 | 5 | 34.4 ± 21.61 | 10.6 ± 4.4 | NA | ND_thermal | H3 | H2 |
| 112 | (24.975, -80.425) | Upper Keys | 21.46 | 1 | 8 | 2 | 4 | 6.5 ± 1.19 | 21.5 ± 8.41 | 5 | 10 ± 5.79 | 27.6 ± 9.06 | NA | NA | NA | 6 | 3.83 ± 1.49 | 13.83 ± 2.74 | H3 | H3 | NA | H2 |
| 115 | (25.025, -80.375) | Upper Keys | 21.67 | 1 | 14 | 20 | 5 | 6.4 ± 2.8 | 32.8 ± 6.09 | 2 | 19 ± 2 | 8.5 ± 2.5 | 5 | 4.8 ± 2.71 | 17.2 ± 8.05 | 14 | 7.14 ± 2.77 | 30.21 ± 5.97 | H3 | H4 | H4 | H3 |
| 117 | (25.075, -80.375) | Upper Keys | 8.92 | 1 | 14 | 1 | 1 | 25 | 19 | 4 | 16.25 ± 2.29 | 20 ± 3.49 | NA | NA | NA | 4 | 22.25 ± 9.34 | 18 ± 3.42 | H3 | H3 | NA | ND_thermal |
| 118 | (25.075, -80.325) | Upper Keys | 26.96 | 1 | 4 | 14 | NA | NA | NA | 4 | 3.25 ± 1.31 | 21.5 ± 10.19 | 2 | 4.5 ± 2.5 | 15 ± 5 | 3 | 6 ± 1.73 | 24.67 ± 10.4 | NA | H3 | H3 | H2 |
| 120 | (25.125, -80.375) | Upper Keys | 13.08 | 1 | 2 | 23 | 1 | 13 | 2 | NA | NA | NA | NA | NA | NA | NA | NA | NA | NA | NA | NA | NA |
| 121 | (25.125, -80.325) | Upper Keys | 11.83 | 1 | 2 | 8 | 2 | 21 ± 9 | 24.5 ± 8.5 | 2 | 15.5 ± 2.5 | 12.5 ± 4.5 | 6 | 6.5 ± 2 | 16.5 ± 5.46 | 5 | 26.2 ± 14.45 | 32.4 ± 3.26 | H2 | H3 | H3 | H2 |
| 126 | (25.225, -80.275) | Upper Keys | 8.32 | 3 | 35.67 ± 2.6 | 18.33 ± 4.7 | 1 | 54 | 35 | 1 | 23 | 35 | 6 | 14.17 ± 4.62 | 19 ± 3.45 | 8 | 20.75 ± 3.99 | 25.62 ± 3.97 | H3 | H3 | ND_thermal | H2 |
| 127 | (25.225, -80.225) | Upper Keys | 13.41 | 6 | 4.17 ± 1.58 | 53.17 ± 11.65 | NA | NA | NA | 2 | 12.5 ± 10.5 | 18.5 ± 2.5 | 3 | 2.33 ± 0.88 | 54 ± 14.53 | 5 | 5 ± 2.88 | 8.8 ± 1.46 | NA | H4 | H3 | H2 |
| 130 | (25.275, -80.225) | Upper Keys | 18.85 | 4 | 20.5 ± 5.95 | 38.75 ± 17.34 | 1 | 7 | 51 | 1 | 8 | 52 | 2 | 15 ± 1 | 38 ± 14 | 4 | 9 ± 3.72 | 20.25 ± 6.43 | H4 | H3 | H4 | H2 |
| 131 | (25.275, -80.175) | Upper Keys | 34.52 | 2 | 4.5 ± 1.5 | 7.5 ± 0.5 | 1 | 4 | 18 | NA | NA | NA | 3 | 2.33 ± 1.33 | 31 ± 14.05 | 1 | 3 | 75 | H3 | NA | H3 | H2 |
| 132 | (25.325, -80.225) | Biscayne | 16.90 | 2 | 20 ± 15 | 4 ± 4 | 1 | 16 | 31 | NA | NA | NA | 1 | 37 | 38 | 5 | 25.4 ± 4.32 | 22.4 ± 7.29 | H2 | NA | ND_thermal | H2 |
| 135 | (25.375, -80.175) | Biscayne | 12.72 | 3 | 9.67 ± 2.91 | 18.33 ± 4.37 | 2 | 13.5 ± 4.5 | 18.5 ± 7.5 | NA | NA | NA | 6 | 11.83 ± 2.9 | 64.67 ± 13.51 | 5 | 21 ± 12.08 | 36.8 ± 16.09 | H2 | NA | H3 | H2 |
| 138 | (25.425, -80.125) | Biscayne | 21.50 | 2 | 5.5 ± 5.5 | 10.5 ± 0.5 | 1 | 5 | 19 | NA | NA | NA | 1 | 4 | 29 | 10 | 8.6 ± 2.86 | 19 ± 4.98 | H2 | NA | H2 | H2 |
| 140 | (25.475, -80.125) | Biscayne | 17.04 | 2 | 9 ± 2 | 28.5 ± 0.5 | 2 | 14.5 ± 2.5 | 29 ± 1 | 1 | 4 | 7 | 5 | 8.8 ± 3.28 | 18.4 ± 4.28 | 9 | 22 ± 4.91 | 36.44 ± 5.45 | H3 | H4 | H4 | H2 |
| 142 | (25.525, -80.125) | Biscayne | 21.14 | 1 | 12 | 19 | 5 | 6.8 ± 0.73 | 8.4 ± 1.86 | 3 | 12.67 ± 5.55 | 13 ± 2.52 | 1 | 12 | 8 | 6 | 19.67 ± 6.38 | 14.67 ± 3.3 | H4 | H4 | ND_thermal | H2 |
| 144 | (25.575, -80.125) | Biscayne | 17.87 | 2 | 5.5 ± 0.5 | 14 ± 13 | 3 | 7.67 ± 1.45 | 5.67 ± 0.88 | 1 | 15 | 36 | NA | NA | NA | 3 | 14.67 ± 10.17 | 14.67 ± 10.68 | H4 | H3 | NA | H2 |
| 145 | (25.575, -80.075) | Biscayne | 24.20 | 2 | 10.5 ± 3.5 | 13.5 ± 6.5 | 1 | 6 | 4 | NA | NA | NA | NA | NA | NA | 1 | 6 | 12 | H4 | NA | NA | H2 |
| 147 | (25.625, -80.075) | Biscayne | 22.94 | 1 | 6 | 3 | 1 | 9 | 6 | NA | NA | NA | 2 | 7.5 ± 2.5 | 20 ± 17 | 1 | 6 | 10 | H3 | NA | H3 | H3 |
| 149 | (25.675, -80.075) | Biscayne | 26.27 | 1 | 6 | 44 | 2 | 3.5 ± 2.5 | 3 ± 2 | 1 | 5 | 15 | 2 | 4.5 ± 0.5 | 24 ± 14 | 3 | 3.67 ± 1.76 | 30.67 ± 15.6 | H4 | H4 | H4 | H4 |
| 150 | (25.725, -80.125) | Biscayne | 25.39 | 1 | 1 | 0 | 1 | 9 | 5 | NA | NA | NA | 1 | 0 | 0 | 1 | 2 | 0 | NA | NA | NA | NA |
| 151 | (25.725, -80.075) | Biscayne | 34.42 | 2 | 1 ± 0 | 12 ± 10 | NA | NA | NA | 1 | 10 | 6 | 2 | 3.5 ± 1.5 | 3.5 ± 0.5 | NA | NA | NA | NA | H3 | H4 | NA |
| 152 | (25.775, -80.125) | Broward-Miami | 22.72 | 2 | 6.5 ± 6.5 | 10.5 ± 4.5 | 2 | 12 ± 9 | 1.5 ± 1.5 | 4 | 1 ± 0.71 | 0.75 ± 0.75 | 1 | 0 | 4 | 4 | 0.75 ± 0.48 | 4.25 ± 1.25 | H4 | H4 | H4 | H4 |
| 153 | (25.775, -80.075) | Broward-Miami | 42.53 | 4 | 4.5 ± 1.5 | 3.5 ± 1.89 | 3 | 2.67 ± 2.19 | 4.67 ± 3.67 | 7 | 2.14 ± 0.63 | 7 ± 2.29 | 7 | 1.29 ± 0.47 | 1.43 ± 0.57 | 4 | 0.75 ± 0.48 | 2.75 ± 1.18 | H4 | H3 | H4 | H4 |
| 154 | (25.825, -80.125) | Broward-Miami | 20.96 | 2 | 3.5 ± 2.5 | 7 ± 3 | 5 | 3 ± 2.07 | 15.6 ± 3.82 | 8 | 1.12 ± 0.4 | 2 ± 0.57 | 5 | 1.4 ± 0.51 | 1.6 ± 0.93 | 5 | 0.6 ± 0.24 | 1 ± 0.32 | H3 | H4 | H4 | H4 |
| 155 | (25.825, -80.075) | Broward-Miami | 44.99 | 1 | 3 | 1 | 3 | 2 ± 1 | 0.33 ± 0.33 | 1 | 2 | 0 | 4 | 1.75 ± 1.11 | 3.75 ± 2.78 | 3 | 2 ± 2 | 2 ± 0 | H4 | H4 | H3 | H3 |
| 156 | (25.875, -80.125) | Broward-Miami | 27.42 | 3 | 2.67 ± 1.33 | 10.33 ± 10.33 | 6 | 3 ± 0.68 | 3.33 ± 1.17 | 9 | 5.44 ± 1.67 | 8.33 ± 2.73 | 9 | 3.22 ± 0.72 | 4.56 ± 1.8 | 6 | 0.83 ± 0.31 | 3.33 ± 0.67 | H4 | H3 | H4 | H4 |
| 157 | (25.875, -80.075) | Broward-Miami | 50.28 | 1 | 6 | 6 | 1 | 6 | 3 | 1 | 1 | 0 | 5 | 2.2 ± 0.66 | 2.6 ± 1.4 | 3 | 1.67 ± 1.2 | 6 ± 4.58 | H2 | NA | H2 | H4 |
| 158 | (25.925, -80.125) | Broward-Miami | 26.89 | 1 | 6 | 2 | 3 | 2.67 ± 0.88 | 2 ± 1 | 7 | 2.86 ± 0.77 | 2.43 ± 1.45 | 9 | 1.11 ± 0.59 | 4.89 ± 1.99 | 13 | 2.46 ± 0.62 | 5.08 ± 0.98 | H2 | H4 | H4 | H4 |
| 159 | (25.925, -80.075) | Broward-Miami | 50.27 | 2 | 2 ± 1 | 5 ± 3 | 2 | 2 ± 0 | 4.5 ± 4.5 | 3 | 1 ± 1 | 2.33 ± 1.2 | 5 | 3.6 ± 0.98 | 4 ± 1.38 | 1 | 0 | 0 | H2 | H4 | H3 | H4 |
| 161 | (25.975, -80.075) | Broward-Miami | 49.75 | 1 | 3 | 5 | 5 | 3.2 ± 0.58 | 2.4 ± 1.6 | 5 | 0.4 ± 0.24 | 0.8 ± 0.58 | 4 | 3 ± 2.68 | 2 ± 1 | 7 | 3.43 ± 1.13 | 4 ± 0.95 | H4 | H4 | H4 | H4 |
| 162 | (26.025, -80.125) | Broward-Miami | 22.94 | 1 | 9 | 2 | 1 | 2 | 0 | 6 | 3.83 ± 0.65 | 9.83 ± 2.39 | 6 | 2.33 ± 0.92 | 5 ± 2.66 | 7 | 3.43 ± 0.97 | 7.57 ± 1.77 | H4 | H3 | H4 | H3 |
| 165 | (26.075, -80.075) | Broward-Miami | 37.32 | 4 | 5.75 ± 1.65 | 6.25 ± 2.39 | 6 | 4.17 ± 0.7 | 5.67 ± 0.88 | 4 | 1.75 ± 0.48 | 3.25 ± 1.11 | 10 | 4.4 ± 0.83 | 3.9 ± 0.98 | 4 | 10.71 | 11.25 ± 2.02 | H4 | H4 | H4 | H3 |
| 167 | (26.125, -80.075) | Broward-Miami | 29.25 | 3 | 13 ± 6.66 | 1 ± 0.58 | 11 | 8.36 ± 3.31 | 11.73 ± 4.59 | 3 | 4 ± 0.58 | 1.33 ± 0.88 | 6 | 3.83 ± 1.58 | 5 ± 2.31 | 7 | 3 ± 1.13 | 6 ± 3.15 | H2 | H4 | H4 | H4 |
| 168 | (26.175, -80.075) | Broward-Miami | 34.09 | 4 | 6 ± 2.35 | 5.25 ± 2.56 | 8 | 4.38 ± 0.73 | 3.12 ± 0.9 | 7 | 1 ± 0.53 | 4.71 ± 1.52 | 10 | 0.9 ± 0.41 | 2.9 ± 0.92 | 8 | 1.5 ± 0.6 | 5.38 ± 1.52 | H4 | H4 | H4 | H4 |
| 169 | (26.225, -80.075) | Broward-Miami | 39.84 | 3 | 5.67 ± 1.86 | 2.67 ± 0.33 | 5 | 4.4 ± 1.69 | 1.4 ± 1.17 | 6 | 3 ± 1.1 | 3.5 ± 1.61 | 6 | 0.33 ± 0.21 | 3.17 ± 0.91 | 4 | 2.5 ± 0.87 | 7.25 ± 3.54 | H4 | H4 | H2 | H3 |
| 170 | (26.275, -80.075) | Broward-Miami | 42.05 | 3 | 11.33 ± 5.67 | 6.33 ± 3.48 | 3 | 8 ± 7.51 | 6.33 ± 5.84 | 1 | 7 | 15 | 2 | 4 ± 3 | 3 ± 1 | NA | NA | NA | H4 | H3 | H4 | NA |
| 178 | (26.625, -80.025) | South Palm Beach | 48.20 | 1 | 1 | 0 | NA | NA | NA | 8 | 1.62 ± 0.5 | 1.75 ± 1.06 | 3 | 1 ± 0.58 | 1.67 ± 0.88 | 5 | 4 ± 1.48 | 2.4 ± 1.6 | NA | NA | NA | NA |
| 179 | (26.675, -80.025) | South Palm Beach | 47.14 | 2 | 1 ± 1 | 0 | 3 | 1 ± 0.58 | 0.67 ± 0.67 | 7 | 2.14 ± 0.4 | 3.43 ± 0.65 | 3 | 5.33 ± 1.33 | 4 ± 3.06 | 6 | 3.5 ± 0.72 | 3.5 ± 1.06 | H3 | H3 | H3 | H3 |
| 180 | (26.725, -80.025) | South Palm Beach | 44.22 | 3 | 0.33 ± 0.33 | 1 ± 1 | 2 | 4.5 ± 3.5 | 3 ± 0 | 1 | 2 | 1 | 8 | 1.38 ± 0.56 | 1.88 ± 0.55 | 6 | 4.83 ± 2.29 | 4.5 ± 0.92 | H3 | NA | H3 | H3 |
| 185 | (27.125, -80.125) | Martin | 15.75 | 2 | 3.5 ± 3.5 | 7 ± 4 | 5 | 0.4 ± 0.24 | 5.4 ± 2.4 | 11 | 0 | 5.91 ± 1.97 | 10 | 0 | 2.6 ± 0.93 | 12 | 0.17 ± 0.11 | 3.25 ± 0.71 | H4 | H4 | H2 | ND_thermal |

**Table S5. Negative binomial GLMs of adult colony density trajectories**

Results of negative binomial GLMs modeling site-level adult colony density (counts) as a function of Timepoint, GridState (H3 vs H4), and their interaction (Eq. 4). Models account for overdispersion in count data and test whether temporal changes in density differ between trajectory classes. So, the 95% confidence intervals represent the uncertainty around the model-estimated proportional change in colony density (i.e., estimated effect size, β), not the distribution of the raw data. p-values are based on Wald z-tests of model coefficients.

Analyses are repeated across adult-size thresholds [W5/S10 (weedy ≥ 5cm; StressT/Gen ≥ 10 cm diameter); W10/S15 (weedy ≥ 10 cm; StressT/Gen ≥ 15 cm diameter); no-threshold) and complemented by juvenile-density models. Results confirm that H3 grids maintain or increase adult density, while H4 grids show systematic decline, supporting the mechanistic distinction between persistence and mortality filtering. For grid classification and descriptive statistics, see Table S4.

| Analysis_Window | Adult_Size_Criteria | Metric | negBGLM Model Term (Eq. 4) | Estimate | Wald 95% CI | p-value | Significance | Supporting Figures |
|---|---|---|---|---|---|---|---|---|
| 2014vs2015 | Weedy ≥5 cm; StressT/ Gen ≥10 cm | Adult Colony Density | Intercept (β0) | 3.6149 | [3.3657, 3.8640] | 6.98E-178 | *** | |
| 2014vs2015 | Weedy ≥5 cm; StressT/ Gen ≥10 cm | Adult Colony Density | Timepoint (β1) | -1.0141 | [-1.3502, -0.6780] | 3.34E-09 | *** | |
| 2014vs2015 | Weedy ≥5 cm; StressT/ Gen ≥10 cm | Adult Colony Density | GridState (β2) | -0.5932 | [-1.1087, -0.0777] | 0.0241 | * | |
| 2014vs2015 | Weedy ≥5 cm; StressT/ Gen ≥10 cm | Adult Colony Density | Interaction (β3) | 1.4186 | [0.7493, 2.0879] | 3.26E-05 | *** | Fig.5j |
| 2014vs2020 | Weedy ≥5 cm; StressT/ Gen ≥10 cm | Adult Colony Density | Intercept (β0) | 3.6667 | [3.4169, 3.9165] | 5.09E-182 | *** | |
| 2014vs2020 | Weedy ≥5 cm; StressT/ Gen ≥10 cm | Adult Colony Density | Timepoint (β1) | -0.6253 | [-0.9441, -0.3066] | 1.20E-04 | *** | |
| 2014vs2020 | Weedy ≥5 cm; StressT/ Gen ≥10 cm | Adult Colony Density | GridState (β2) | -0.4624 | [-0.8647, -0.0600] | 0.0243 | * | |
| 2014vs2020 | Weedy ≥5 cm; StressT/ Gen ≥10 cm | Adult Colony Density | Interaction (β3) | 0.805 | [0.2959, 1.3141] | 0.00194 | ** | Fig.5k |
| 2014vs2022 | Weedy ≥5 cm; StressT/ Gen ≥10 cm | Adult Colony Density | Intercept (β0) | 3.6113 | [3.3561, 3.8665] | 2.55E-169 | *** | |
| 2014vs2022 | Weedy ≥5 cm; StressT/ Gen ≥10 cm | Adult Colony Density | Timepoint (β1) | -0.5646 | [-0.8816, -0.2476] | 4.81E-04 | *** | |
| 2014vs2022 | Weedy ≥5 cm; StressT/ Gen ≥10 cm | Adult Colony Density | GridState (β2) | -0.3625 | [-0.7606, 0.0357] | 0.0743 | . | |
| 2014vs2022 | Weedy ≥5 cm; StressT/ Gen ≥10 cm | Adult Colony Density | Interaction (β3) | 1.0131 | [0.5174, 1.5088] | 6.18E-05 | *** | Fig.5l |
| 2014vs2023 | Weedy ≥5 cm; StressT/ Gen ≥10 cm | Adult Colony Density | Intercept (β0) | 2.8945 | [2.4673, 3.3217] | 3.01E-40 | *** | |
| 2014vs2023 | Weedy ≥5 cm; StressT/ Gen ≥10 cm | Adult Colony Density | Timepoint (β1) | -0.3172 | [-0.8232, 0.1889] | 0.219 | ns | |
| 2014vs2023 | Weedy ≥5 cm; StressT/ Gen ≥10 cm | Adult Colony Density | GridState (β2) | -0.0829 | [-0.7211, 0.5553] | 0.799 | ns | |
| 2014vs2023 | Weedy ≥5 cm; StressT/ Gen ≥10 cm | Adult Colony Density | Interaction (β3) | 0.9501 | [0.1989, 1.7013] | 0.0132 | * | Fig.6d |
| 2014vs2015 | Weedy ≥10; StressT/Gen ≥15 cm | Adult Colony Density | Intercept (β0) | 3.0994 | [2.7802, 3.4186] | 9.37E-81 | *** | no figure- sensitivity analysis |
| 2014vs2015 | Weedy ≥10; StressT/Gen ≥15 cm | Adult Colony Density | Timepoint (β1) | -0.9781 | [-1.4001, -0.5561] | 5.54E-06 | *** | |
| 2014vs2015 | Weedy ≥10; StressT/Gen ≥15 cm | Adult Colony Density | GridState (β2) | -0.3438 | [-0.9062, 0.2185] | 0.231 | ns | |
| 2014vs2015 | Weedy ≥10; StressT/Gen ≥15 cm | Adult Colony Density | Interaction (β3) | 0.9176 | [0.1608, 1.6743] | 0.0175 | * | |
| 2014vs2020 | Weedy ≥10; StressT/Gen ≥15 cm | Adult Colony Density | Intercept (β0) | 3.2105 | [2.9445, 3.4766] | 1.05E-123 | *** | |
| 2014vs2020 | Weedy ≥10; StressT/Gen ≥15 cm | Adult Colony Density | Timepoint (β1) | -0.8511 | [-1.1932, -0.5091] | 1.08E-06 | *** | |
| 2014vs2020 | Weedy ≥10; StressT/Gen ≥15 cm | Adult Colony Density | GridState (β2) | -0.9105 | [-1.3854, -0.4357] | 1.71E-04 | *** | |
| 2014vs2020 | Weedy ≥10; StressT/Gen ≥15 cm | Adult Colony Density | Interaction (β3) | 1.2448 | [0.6481, 1.8415] | 4.34E-05 | *** | |
| 2014vs2022 | Weedy ≥10; StressT/Gen ≥15 cm | Adult Colony Density | Intercept (β0) | 3.1172 | [2.8473, 3.3870] | 1.77E-113 | *** | |
| 2014vs2022 | Weedy ≥10; StressT/Gen ≥15 cm | Adult Colony Density | Timepoint (β1) | -0.6593 | [-0.9955, -0.3230] | 1.21E-04 | *** | |
| 2014vs2022 | Weedy ≥10; StressT/Gen ≥15 cm | Adult Colony Density | GridState (β2) | -0.5198 | [-0.9808, -0.0587] | 0.0271 | * | |
| 2014vs2022 | Weedy ≥10; StressT/Gen ≥15 cm | Adult Colony Density | Interaction (β3) | 1.0489 | [0.4746, 1.6232] | 3.44E-04 | *** | |
| 2014vs2023 | Weedy ≥10; StressT/Gen ≥15 cm | Adult Colony Density | Intercept (β0) | 2.3231 | [1.8125, 2.8336] | 4.73E-19 | *** | |
| 2014vs2023 | Weedy ≥10; StressT/Gen ≥15 cm | Adult Colony Density | Timepoint (β1) | -0.3301 | [-0.9194, 0.2591] | 0.272 | ns | |
| 2014vs2023 | Weedy ≥10; StressT/Gen ≥15 cm | Adult Colony Density | GridState (β2) | -0.0824 | [-0.8825, 0.7178] | 0.84 | ns | |
| 2014vs2023 | Weedy ≥10; StressT/Gen ≥15 cm | Adult Colony Density | Interaction (β3) | 0.9662 | [0.0035, 1.9289] | 0.0492 | * | |
| 2014vs2015 | NoThreshold | Total Colony Density | Intercept (β0) | 3.8553 | [3.6464, 4.0641] | 1.33E-286 | *** | no figure- sensitivity analysis |
| 2014vs2015 | NoThreshold | Total Colony Density | Timepoint (β1) | -0.7921 | [-1.0759, -0.5084] | 4.45E-08 | *** | |
| 2014vs2015 | NoThreshold | Total Colony Density | GridState (β2) | -0.574 | [-1.0182, -0.1297] | 0.0113 | * | |
| 2014vs2015 | NoThreshold | Total Colony Density | Interaction (β3) | 1.1416 | [0.5740, 1.7092] | 8.08E-05 | *** | |
| 2014vs2020 | NoThreshold | Total Colony Density | Intercept (β0) | 3.9435 | [3.7260, 4.1611] | 1.74E-276 | *** | |
| 2014vs2020 | NoThreshold | Total Colony Density | Timepoint (β1) | -0.5635 | [-0.8482, -0.2789] | 1.04E-04 | *** | |
| 2014vs2020 | NoThreshold | Total Colony Density | GridState (β2) | -0.5326 | [-0.8692, -0.1959] | 0.00193 | ** | |
| 2014vs2020 | NoThreshold | Total Colony Density | Interaction (β3) | 0.8948 | [0.4702, 1.3194] | 3.62E-05 | *** | |
| 2014vs2022 | NoThreshold | Total Colony Density | Intercept (β0) | 3.7881 | [3.5773, 3.9990] | 1.33E-271 | *** | |
| 2014vs2022 | NoThreshold | Total Colony Density | Timepoint (β1) | -0.4891 | [-0.7512, -0.2271] | 2.54E-04 | *** | |
| 2014vs2022 | NoThreshold | Total Colony Density | GridState (β2) | -0.2137 | [-0.5563, 0.1289] | 0.222 | ns | |
| 2014vs2022 | NoThreshold | Total Colony Density | Interaction (β3) | 0.9943 | [0.5677, 1.4209] | 4.93E-06 | *** | |
| 2014vs2023 | NoThreshold | Total Colony Density | Intercept (β0) | 3.1336 | [2.7593, 3.5079] | 1.61E-60 | *** | |
| 2014vs2023 | NoThreshold | Total Colony Density | Timepoint (β1) | -0.3504 | [-0.8198, 0.1190] | 0.143 | ns | |
| 2014vs2023 | NoThreshold | Total Colony Density | GridState (β2) | 0.0837 | [-0.4296, 0.5970] | 0.749 | ns | |
| 2014vs2023 | NoThreshold | Total Colony Density | Interaction (β3) | 0.7418 | [0.1246, 1.3589] | 0.0185 | * | |
| 2014vs2015 | Weedy ≥5 cm; StressT/ Gen ≥10 cm | Juvenile Colony Density | Intercept (β0) | 2.3859 | [2.1538, 2.6180] | 2.82E-90 | *** | no figure- sensitivity analysis |
| 2014vs2015 | Weedy ≥5 cm; StressT/ Gen ≥10 cm | Juvenile Colony Density | Timepoint (β1) | -0.3065 | [-0.6189, 0.0060] | 0.0546 | . | |
| 2014vs2015 | Weedy ≥5 cm; StressT/ Gen ≥10 cm | Juvenile Colony Density | GridState (β2) | -0.8417 | [-1.3431, -0.3403] | 0.001 | ** | |
| 2014vs2015 | Weedy ≥5 cm; StressT/ Gen ≥10 cm | Juvenile Colony Density | Interaction (β3) | 0.8287 | [0.1860, 1.4714] | 0.0115 | * | |
| 2014vs2020 | Weedy ≥5 cm; StressT/ Gen ≥10 cm | Juvenile Colony Density | Intercept (β0) | 2.4003 | [2.1873, 2.6134] | 5.09E-108 | *** | |
| 2014vs2020 | Weedy ≥5 cm; StressT/ Gen ≥10 cm | Juvenile Colony Density | Timepoint (β1) | 0.0088 | [-0.2620, 0.2797] | 0.949 | ns | |
| 2014vs2020 | Weedy ≥5 cm; StressT/ Gen ≥10 cm | Juvenile Colony Density | GridState (β2) | -0.5875 | [-0.9380, -0.2371] | 0.00102 | ** | |
| 2014vs2020 | Weedy ≥5 cm; StressT/ Gen ≥10 cm | Juvenile Colony Density | Interaction (β3) | 0.6528 | [0.2140, 1.0915] | 0.00354 | ** | |
| 2014vs2022 | Weedy ≥5 cm; StressT/ Gen ≥10 cm | Juvenile Colony Density | Intercept (β0) | 2.1941 | [1.9739, 2.4143] | 6.45E-85 | *** | |
| 2014vs2022 | Weedy ≥5 cm; StressT/ Gen ≥10 cm | Juvenile Colony Density | Timepoint (β1) | -0.2238 | [-0.4980, 0.0504] | 0.11 | ns | |
| 2014vs2022 | Weedy ≥5 cm; StressT/ Gen ≥10 cm | Juvenile Colony Density | GridState (β2) | -0.1122 | [-0.4562, 0.2319] | 0.523 | ns | |
| 2014vs2022 | Weedy ≥5 cm; StressT/ Gen ≥10 cm | Juvenile Colony Density | Interaction (β3) | 0.821 | [0.3945, 1.2475] | 1.61E-04 | *** | |
| 2014vs2023 | Weedy ≥5 cm; StressT/ Gen ≥10 cm | Juvenile Colony Density | Intercept (β0) | 1.8101 | [1.4737, 2.1465] | 5.33E-26 | *** | |
| 2014vs2023 | Weedy ≥5 cm; StressT/ Gen ≥10 cm | Juvenile Colony Density | Timepoint (β1) | 0.191 | [-0.2047, 0.5866] | 0.344 | ns | |
| 2014vs2023 | Weedy ≥5 cm; StressT/ Gen ≥10 cm | Juvenile Colony Density | GridState (β2) | 0.1614 | [-0.3364, 0.6593] | 0.525 | ns | |
| 2014vs2023 | Weedy ≥5 cm; StressT/ Gen ≥10 cm | Juvenile Colony Density | Interaction (β3) | 0.1504 | [-0.4324, 0.7331] | 0.613 | ns | |
| 2014vs2015 | Weedy ≥10; SressT/Gen ≥15 cm | Juvenile Colony Density | Intercept (β0) | 3.2917 | [3.0755, 3.5079] | 1.19E-195 | *** | |
| 2014vs2015 | Weedy ≥10; SressT/Gen ≥15 cm | Juvenile Colony Density | Timepoint (β1) | -0.4996 | [-0.7845, -0.2147] | 5.89E-04 | *** | |
| 2014vs2015 | Weedy ≥10; SressT/Gen ≥15 cm | Juvenile Colony Density | GridState (β2) | -0.6947 | [-1.0812, -0.3083] | 4.26E-04 | *** | |
| 2014vs2015 | Weedy ≥10; SressT/Gen ≥15 cm | Juvenile Colony Density | Interaction (β3) | 0.6063 | [0.0878, 1.1249] | 0.0219 | * | |
| 2014vs2020 | Weedy ≥10; SressT/Gen ≥15 cm | Juvenile Colony Density | Intercept (β0) | 3.299 | [3.1066, 3.4914] | 1.25E-247 | *** | |
| 2014vs2020 | Weedy ≥10; SressT/Gen ≥15 cm | Juvenile Colony Density | Timepoint (β1) | -0.1702 | [-0.4163, 0.0759] | 0.175 | ns | |
| 2014vs2020 | Weedy ≥10; SressT/Gen ≥15 cm | Juvenile Colony Density | GridState (β2) | -0.7203 | [-1.0648, -0.3758] | 4.16E-05 | *** | |
| 2014vs2020 | Weedy ≥10; SressT/Gen ≥15 cm | Juvenile Colony Density | Interaction (β3) | 0.8246 | [0.3937, 1.2556] | 1.77E-04 | *** | |
| 2014vs2022 | Weedy ≥10; SressT/Gen ≥15 cm | Juvenile Colony Density | Intercept (β0) | 3.1087 | [2.9017, 3.3158] | 2.34E-190 | *** | |
| 2014vs2022 | Weedy ≥10; SressT/Gen ≥15 cm | Juvenile Colony Density | Timepoint (β1) | -0.1236 | [-0.3806, 0.1334] | 0.346 | ns | |
| 2014vs2022 | Weedy ≥10; SressT/Gen ≥15 cm | Juvenile Colony Density | GridState (β2) | -0.1166 | [-0.4687, 0.2356] | 0.516 | ns | |
| 2014vs2022 | Weedy ≥10; SressT/Gen ≥15 cm | Juvenile Colony Density | Interaction (β3) | 0.6487 | [0.2110, 1.0864] | 0.00367 | ** | |
| 2014vs2023 | Weedy ≥10; SressT/Gen ≥15 cm | Juvenile Colony Density | Intercept (β0) | 2.6391 | [2.3199, 2.9582] | 4.63E-59 | *** | |
| 2014vs2023 | Weedy ≥10; SressT/Gen ≥15 cm | Juvenile Colony Density | Timepoint (β1) | 0.2532 | [-0.1134, 0.6198] | 0.176 | ns | |
| 2014vs2023 | Weedy ≥10; SressT/Gen ≥15 cm | Juvenile Colony Density | GridState (β2) | 0.0247 | [-0.4746, 0.5240] | 0.923 | ns | |
| 2014vs2023 | Weedy ≥10; SressT/Gen ≥15 cm | Juvenile Colony Density | Interaction (β3) | 0.2589 | [-0.3399, 0.8577] | 0.397 | ns | |

**Table S6. Baseline demographic and thermal context of stable pre-2023 trajectory networks**

Grid cells (5 × 5 km) were classified as Stable H2, Stable H3, or Stable H4 using only pre-2023 trajectory assignments (2014–2015, 2014–2020, 2014–2022, see Table S4) based on consistency criteria (≥2 concordant classifications, no opposing classifications). Baseline demographic metrics (2014 only) were computed using two-step averaging with true zeros retained. Thermal variability (Standard Deviation, SD) summarizes DHW across pre-2023 MHWs. Group differences between Stable H3 and Stable H4 are evaluated using Mann–Whitney U tests (equivalent to the Wilcoxon rank-sum tests), with Cliff's δ (95% CI) reported as a distribution-free effect size. Stable H2 grids were rare (n=3) and are provided just for comparison. Results indicate that baseline adult community composition differs between trajectory types, with suggestive but lower-power differences in thermal regimes. Stable H3 Grid ID numbers: 9, 23, 68, 107, 112, 117, 118, 126, 131, 147, 179, 180. Stable H4 Grid ID numbers: 22, 25, 42, 55, 56, 57, 63, 142, 149, 152, 161, 165, 168.

| Metric | Stable H3 grids (n=12) | Stable H4 grids (n=13) | p-value (Mann-Whitney U test) | Cliff's Delta | 95% CI | Stable H2 grids (n=3) (comparison only) |
|---|---|---|---|---|---|---|
| Avg_Depth (m) | 25.17 ± 12.52 | 29.11 ± 8.34 | 0.399 | -0.205 | [-0.667, 0.295] | 39.59 ± 16.40 |
| Total Surveyed Sites (per grid) | 1.58 ± 0.79 | 2.38 ± 1.66 | 0.275 | -0.244 | [-0.635, 0.186] | 2.00 ± 1.00 |
| Weedy Adults (≥5cm) | 6.94 ± 7.10 | 17.88 ± 10.98 | 0.011* | -0.603 | [-0.897, -0.218] | 7.50 ± 2.60 |
| Weedy Juvs (<5cm) | 0.17 ± 0.39 | 0.31 ± 0.60 | 0.49 | -0.128 | [-0.462, 0.192] | 0.33 ± 0.58 |
| ST/Gen Adults (≥10cm) | 8.00 ± 9.50 | 27.97 ± 27.87 | 0.064 | -0.442 | [-0.782, 0.019] | 9.61 ± 6.69 |
| ST/Gen Juvs (<10cm) | 4.81 ± 3.85 | 10.56 ± 8.80 | 0.096 | -0.397 | [-0.769, 0.051] | 4.06 ± 1.78 |
| 2014 DHW (°C-weeks) | 3.63 ± 2.07 | 3.77 ± 2.22 | 0.724 | -0.09 | [-0.526, 0.410] | 2.47 ± 0.89 |
| 2015 DHW (°C-weeks) | 3.28 ± 2.42 | 3.28 ± 2.37 | 0.644 | -0.13 | [-0.630, 0.407] | 1.79 ± 1.04 |
| 2020 DHW (°C-weeks) | 2.85 ± 2.14 | 2.77 ± 2.70 | 0.778 | -0.077 | [-0.538, 0.423] | 0.31 ± 0.44 |
| 2022 DHW (°C-weeks) | 3.35 ± 2.38 | 4.07 ± 2.38 | 0.598 | -0.138 | [-0.615, 0.369] | 3.21 ± 1.06 |
| 2023 DHW (°C-weeks) | 11.12 ± 4.46 | 13.67 ± 3.29 | 0.149 | -0.346 | [-0.744, 0.090] | 12.70 ± 1.23 |
| DHW Standard Dev: 4 MHWs | 0.76 ± 0.48 | 1.04 ± 0.48 | 0.183 | -0.321 | [-0.756, 0.179] | 1.07 ± 0.43 |
| DHW Standard Dev: 5 MHWs | 3.81 ± 1.71 | 4.68 ± 0.95 | 0.265 | -0.269 | [-0.718, 0.205] | 5.14 ± 0.39 |

**Table S7. Ultimate refugia: persistence of stable trajectory networks under the 2023 stress test**

This table identifies "ultimate" grids (i.e., Stable H3 or Stable H4 based on pre-2023 trajectories) (see Table S4 and Table S6) and that retained their trajectory classification in the 2014–2023 comparison. Metrics include baseline demographic properties, thermal variability across pre-2023 and full MHW series, and 2023 heat exposure. Group differences (Ultimate H3 vs Ultimate H4) are evaluated using Mann–Whitney U tests (equivalent to the Wilcoxon rank-sum tests) and Cliff's δ (95% CI). Because sample sizes are small (n = 4 H3, n = 5 H4), inference emphasizes effect sizes and directional consistency rather than strict p-value thresholds. No ultimate H2 grids were identified. Results suggest that persistence is associated with lower multi-event thermal variability, consistent with the predictable exposure hypothesis. Ultimate H3 Grid ID number: 107, 117, 147, 179 (8 lost H3 status). Ultimate H4 Grid ID number: 56, 149, 152, 161, 168 (8 lost H4 status).

| Metric | Ultimate H3 grids (n=4) | Ultimate H4 grids (n=5) | p-value (Mann-Whitney U test) | Cliff's Delta | 95% CI |
|---|---|---|---|---|---|
| Weedy Adults (≥5cm) | 2.75 ± 3.10 | 15.55 ± 16.27 | 0.063 | -0.8 | [-1.000, -0.200] |
| Weedy Juvs (<5cm) | 0.00 ± 0.00 | 0.55 ± 0.87 | 0.24 | -0.4 | [-0.800, 0.000] |
| ST/Gen Adults (≥10cm) | 3.08 ± 2.83 | 10.80 ± 12.21 | 0.105 | -0.7 | [-1.000, -0.100] |
| ST/Gen Juvs (<10cm) | 2.92 ± 1.77 | 7.10 ± 6.80 | 0.325 | -0.45 | [-1.000, 0.400] |
| 2014 DHW (°C-weeks) | 2.34 ± 1.79 | 2.53 ± 2.13 | 0.73 | -0.2 | [-1.000, 0.600] |
| 2015 DHW (°C-weeks) | 0.75 ± 1.06 | 1.76 ± 1.96 | 0.393 | -0.467 | [-1.000, 0.600] |
| 2020 DHW (°C-weeks) | 1.61 ± 2.79 | 1.40 ± 2.88 | 0.751 | -0.2 | [-1.000, 0.600] |
| 2022 DHW (°C-weeks) | 2.67 ± 2.08 | 2.93 ± 2.37 | 1 | 0 | [-0.902, 0.800] |
| 2023 DHW (°C-weeks) | 9.99 ± 1.57 | 12.69 ± 3.00 | 0.111 | -0.7 | [-1.000, 0.000] |
| DHW Standard Dev: 4 MHWs Only | 0.52 ± 0.15 | 0.83 ± 0.26 | 0.063 | -0.8 | [-1.000, -0.200] |
| DHW Standard Dev: 5 MHWs Only | 3.71 ± 0.40 | 4.77 ± 0.38 | 0.015* | -1 | [-1.000, -1.000] |

**Table S8. Species-level decomposition of H3–H4 divergence (2014–2023)**

Species-specific changes in adult colony density ($\Delta CD_{adult,grid}$ 2014-2023) were evaluated for 22 ecologically important coral taxa across H3 and H4 grids (see Table S4). Density was calculated at the site level with true zeros retained, then aggregated to grid-level means using two-step averaging. Differences between H3 and H4 were assessed using Welch's t-tests (unequal variances), with mean differences, 95% confidence intervals, and p-values reported. This table supports Figure 7 and demonstrates that network-level divergence is driven by selective gains in a subset of taxa (e.g., PAST, PPOR), rather than uniform community-wide change.

| Species Acronym | Full species names | Number H3 adult_colonies_ 2014 | Number H3 adult_colonies_ 2023 | Number H4 adult_colonies_ 2014 | Number H4 adult_colonies_ 2023 | Number H3 sites_2014 | Number H3 sites_2023 | Number H4_sites _2014 | Number H4_sites _2023 | Number H3_Grids _2014 | Number H3_Grids_ 2023 | Number H4_Grids _2014 | Number H4_Grids _2023 | Mean ± SE ΔCDgrid, adult (H3) | Mean ± SE ΔCDgrid, adult (H4) | Mean Difference; ΔCDgrid,adult (H3- H4) | Welch's t-test [95% CI] | Welch's t-test, p-value | Significance | Supporting Figure |
|---|---|---|---|---|---|---|---|---|---|---|---|---|---|---|---|---|---|---|---|---|
| PAST | *Porites astreoides* | 84 | 670 | 145 | 250 | 15 | 50 | 21 | 57 | 7 | 11 | 12 | 12 | 5.8444 ± 1.7566 | -3.5655 ± 1.8638 | 9.41 | [4.10, 14.72] | 0.00 | ** | Figure 7 |
| SSID | *Siderastrea siderea* | 101 | 482 | 85 | 269 | 15 | 36 | 19 | 29 | 9 | 10 | 12 | 10 | 5.1145 ± 2.7854 | 0.9642 ± 1.8810 | 4.15 | [-2.91, 11.21] | 0.23 | ns | Figure 7 |
| PPOR | *Porites porites* | 9 | 144 | 57 | 53 | 5 | 30 | 11 | 26 | 4 | 8 | 7 | 10 | 1.1307 ± 0.5470 | -1.2002 ± 0.8017 | 2.33 | [0.31, 4.35] | 0.03 | * | Figure 7 |
| MCAV | *Montastraea cavernosa* | 26 | 109 | 44 | 36 | 11 | 29 | 15 | 20 | 7 | 10 | 8 | 10 | 1.3481 ± 0.9929 | -0.7757 ± 0.3529 | 2.12 | [-0.16, 4.41] | 0.07 | . | Figure 7 |
| SINT | *Stephanocoenia intersepta* | 36 | 143 | 65 | 139 | 9 | 27 | 15 | 34 | 6 | 10 | 8 | 12 | 1.6266 ± 0.8305 | 0.1181 ± 1.0127 | 1.51 | [-1.21, 4.23] | 0.26 | ns | Figure 7 |
| DSTO | *Dichocoenia stokesii* | 3 | 8 | 18 | 4 | 3 | 8 | 9 | 4 | 3 | 4 | 7 | 4 | -0.0249 ± 0.1228 | -0.8595 ± 0.3927 | 0.83 | [-0.05, 1.72] | 0.06 | . | Figure 7 |
| SRAD | *Siderastrea radians* | 45 | 47 | 27 | 40 | 10 | 14 | 8 | 20 | 6 | 7 | 8 | 8 | -0.5301 ± 0.3701 | -1.3194 ± 1.3393 | 0.79 | [-2.19, 3.77] | 0.58 | ns | Figure 7 |
| OFAV | *Orbicella faveolata* | 4 | 34 | 6 | 9 | 3 | 14 | 4 | 6 | 3 | 5 | 3 | 4 | 0.3550 ± 0.3465 | -0.0082 ± 0.1484 | 0.36 | [-0.45, 1.17] | 0.35 | ns | Figure 7 |
| DLAB | *Diploria labyrinthiformis* | 3 | 5 | 6 | 2 | 2 | 5 | 3 | 2 | 1 | 4 | 3 | 2 | 0.0887 ± 0.1232 | -0.2473 ± 0.1698 | 0.34 | [-0.10, 0.77] | 0.12 | ns | Figure 7 |
| AAGA | *Agaricia agaricites* | 22 | 158 | 24 | 87 | 6 | 32 | 9 | 16 | 5 | 9 | 8 | 8 | 0.7119 ± 0.9335 | 0.4001 ± 0.4011 | 0.31 | [-1.87, 2.50] | 0.76 | ns | Figure 7 |
| PSTR | *Pseudodiploria strigosa* | 9 | 5 | 12 | 5 | 7 | 5 | 7 | 5 | 5 | 4 | 6 | 4 | -0.2067 ± 0.1211 | -0.4194 ± 0.2428 | 0.21 | [-0.36, 0.78] | 0.44 | ns | Figure 7 |
| MDEC | *Madracis decactis* | 1 | 7 | 2 | 3 | 1 | 5 | 2 | 2 | 1 | 4 | 2 | 2 | 0.1115 ± 0.0712 | -0.0082 ± 0.0461 | 0.12 | [-0.06, 0.30] | 0.18 | ns | Figure 7 |
| PCLI | *Pseudodiploria clivosa* | 3 | 4 | 4 | 5 | 3 | 3 | 3 | 4 | 3 | 1 | 3 | 4 | -0.0920 ± 0.1131 | -0.1082 ± 0.1124 | 0.02 | [-0.31, 0.35] | 0.92 | ns | Figure 7 |
| CNAT | *Colpophyllia natans* | 15 | 12 | 5 | 3 | 5 | 6 | 3 | 2 | 5 | 4 | 3 | 1 | -0.3788 ± 0.2790 | -0.2771 ± 0.2323 | -0.10 | [-0.86, 0.65] | 0.78 | ns | Figure 7 |
| ALAM | *Agaricia lamarcki* | 3 | 2 | 3 | 2 | 3 | 2 | 2 | 2 | 3 | 2 | 2 | 2 | -0.2197 ± 0.1550 | -0.0916 ± 0.0870 | -0.13 | [-0.51, 0.25] | 0.48 | ns | Figure 7 |
| PDIV | *Porites divaricata* | 0 | 17 | 0 | 24 | 0 | 5 | 0 | 5 | 0 | 3 | 0 | 3 | 0.1526 ± 0.0914 | 0.3538 ± 0.2761 | -0.20 | [-0.82, 0.42] | 0.50 | ns | Figure 7 |
| EFAS | *Eusmilia fastigiata* | 6 | 2 | 3 | 6 | 5 | 2 | 2 | 5 | 4 | 2 | 2 | 3 | -0.2511 ± 0.1336 | -0.0201 ± 0.0850 | -0.23 | [-0.56, 0.10] | 0.16 | ns | Figure 7 |
| OFRA | *Orbicella franksi* | 4 | 0 | 0 | 0 | 2 | 0 | 0 | 0 | 2 | 0 | 0 | 0 | -0.2955 ± 0.2714 | 0.0000 ± 0.0000 | -0.30 | [-0.90, 0.31] | 0.30 | ns | Figure 7 |
| PFUR | *Porites furcata* | 6 | 14 | 0 | 0 | 3 | 4 | 0 | 0 | 2 | 2 | 0 | 0 | -0.3095 ± 0.3824 | 0.0000 ± 0.0000 | -0.31 | [-1.16, 0.54] | 0.44 | ns | Figure 7 |
| SBOU | *Solenastrea bournoni* | 14 | 5 | 15 | 23 | 9 | 4 | 9 | 17 | 7 | 3 | 8 | 11 | -0.5753 ± 0.2495 | -0.2233 ± 0.2115 | -0.35 | [-1.03, 0.33] | 0.29 | ns | Figure 7 |
| MMEA | *Meandrina meandrites* | 11 | 6 | 12 | 13 | 7 | 6 | 9 | 12 | 6 | 4 | 7 | 5 | -0.7143 ± 0.3304 | -0.3088 ± 0.1728 | -0.41 | [-1.20, 0.39] | 0.29 | ns | Figure 7 |
| OANN | *Orbicella annularis* | 8 | 5 | 0 | 0 | 2 | 3 | 0 | 0 | 2 | 2 | 0 | 0 | -0.5803 ± 0.6456 | 0.00 | -0.58 | [-2.02, 0.86] | 0.39 | ns | Figure 7 |

**Table S9. Species sensitivity analysis using pre-2023 stable network labels**

Out-of-sample validation of species-level patterns using network labels defined exclusively from pre-2023 trajectories (Stable H3 vs Stable H4, see Table S4 and Table S6), thereby avoiding circularity with 2023 outcomes. Analyses focus on four focal taxa (PAST, PPOR, MCAV, SSID) representing key ecological strategies. Species-specific changes in adult colony density (2014-2023) were compared between stable network types using Welch's t-tests), with mean differences, 95% confidence intervals, and p-values reported. Results confirm that key patterns identified in Table S8 persist under independent labeling, strengthening inference that species-level drivers reflect underlying system dynamics rather than classification artifacts.

| Species | Number colonies in stable H3 grids (n=12) | Number colonies in stable H4 grids (n=13) | Mean ± SE ΔCDgrid,adult (Stable H3) | Mean ± SE ΔCDgrid,adult (Stable H4) | Mean Difference; ΔCDgrid,adult (H3-H4) | Welch's t-test [95% CI] | p_value (Welch's t-test) |
|---|---|---|---|---|---|---|---|
| MCAV (ST/Gen) | 106 | 248 | 0.67 ± 0.61 | 1.01 ± 0.95 | -0.34 | [-2.7, 2.02] | 0.77 |
| PAST (Weedy) | 638 | 1009 | 5.85 ± 1.75 | -0.76 ± 1.96 | 6.62 | [1.17, 12.07] | 0.02 |
| PPOR (Weedy) | 197 | 187 | 1.41 ± 1.06 | -1.76 ± 0.79 | 3.18 | [0.42, 5.95] | 0.03 |
| SSID (ST/Gen) | 779 | 1330 | 5.63 ± 2.55 | 3.031 ± 3.17 | 2.60 | [-5.84, 11.04] | 0.53 |
